# Development of novel flavonoid senolytics through phenotypic drug screening and drug design

**DOI:** 10.1101/2024.10.27.620529

**Authors:** Lei Justan Zhang, Rahagir Salekeen, Carolina Soto-Palma, Yuanjun He, Osama Elsallabi, Brian Hughes, Allancer Nunes, Wandi Xu, Borui Zhang, Abdalla Mohamed, Sara J. McGowan, Luise Angelini, Ryan O’Kelly, Theodore M. Kamenecka, Laura J. Niedernhofer, Paul D. Robbins

## Abstract

Accumulation of senescent cells drives aging and age-related diseases. Senolytics, which selectively kill senescent cells, offer a promising approach for treating many age-related diseases. Using a senescent cell-based phenotypic drug discovery approach that combines drug screening and drug design, we developed two novel flavonoid senolytics, SR29384 and SR31133, derived from the senolytic fisetin. These compounds demonstrated enhanced senolytic activities, effectively eliminating multiple senescent cell types, reducing tissue senescence *in vivo*, and extending healthspan in a mouse model of accelerated aging. Mechanistic studies utilizing RNA-Seq, machine learning, network pharmacology, and computational simulation suggest that these novel flavonoid senolytics target PARP1, BCL-xL, and CDK2 to induce selective senescent cell death. This phenotype-based discovery of novel flavonoid senolytics, coupled with mechanistic insights, represents a key advancement in developing next-generation senolyticss with potential clinical applications in treating aging and age-related diseases.

## Introduction

Aging is associated with declines in health and physiological functions over time, leading to an increased prevalence of various chronic diseases. One of the key hallmarks or underlying mechanisms of aging is cellular senescence.^1–3^ Senescence is a state in which cells lose their ability to proliferate in response to various stress conditions, resulting in stable cell cycle arrest.^4^ With age, senescent cells accumulate in tissues and release a range of deleterious signals known as the senescence-associated secretory phenotype (SASP).^5,6^ The phenotype includes diverse pro-inflammatory cytokines, chemokines, proteases, nucleic acid, and metabolites that can have adverse effects both locally or systemically through paracrine and endocrine signaling. The accumulative effect of senescent cells and the SASP results in tissue dysfunction, accelerated aging, and the promotion of various age-related diseases.^7–9^

The use of transgenic mouse models, in which p16^Ink4a+^ or p21^Cip1+^ senescent cells can be selectively ablated, have demonstrated the critical role of senescent cells in driving aging and age-related diseases.^10–12^ More importantly, pharmacological approaches that target and eliminate senescent cells using a class of drugs known as senolytics have shown promise in extending healthspan and treating many age-related diseases.^13–15^ Despite the promise, the number of effective senolytics remains limited, with most being discovered through focused library screening of known compounds, such as dasatinib combined with quercetin, navitoclax, fisetin, and others.^16–18^

Fisetin, a naturally occurring flavonoid, was identified as a senotherapeutic capable of reducing cellular senescence, suppressing age-related pathology, and extending healthspan in aged mice.^19,20^ Based on these preclinical results, fisetin is currently undergoing clinical trials for various conditions, such as chronic kidney disease (NCT03325322), skeletal health (NCT04313634), osteoarthritis (NCT04210986), COVID- 19 (NCT04476953, NCT04537299), and frailty (NCT03675724). However, fisetin is not a strong senolytic, requiring high doses to reduce the senescent cell burden in animal studies^19^. Moreover, fisetin suffers from low bioavailability, likely due to its four hydroxyl groups, which result in low aqueous solubility and poor gut absorption.^21,22^ Therefore, there is a clear need to develop safer and more effective fisetin analogs that can be translated into clinical applications for the treatment of age-related diseases.

Here we report the development of novel flavonoid-based senolytics by a phenotypic drug discovery approach that integrates senescent cell-based drug screening and drug design. Starting without a predefined drug target, we leveraged the relative senolytic activities of fisetin and related flavonoids to obtain phenotype-driven structure-activity relationship (SAR) insights to guide the design of novel flavonoid senolytics with enhanced efficacy. The optimized flavonoid analogs were evaluated for their ability to reduce cellular senescence and extend healthspan in mice. To investigate the potential mechanisms underlying these novel senolytics, we employed a comprehensive approach combining transcriptomics, machine learning, network pharmacology, and flexible molecular docking.

## Results

### Phenotypic drug screening

Flavonoids are a diverse class of polyphenolic natural products found in various fruits, vegetables, and plants. These compounds are characterized by a diphenylpropane scaffold, commonly referred to as a C_6_-C_3_-C_6_ structural framework and are categorized into several subgroups based on their chemical structures, including flavones, isoflavones, flavonols, flavanols, and anthocyanidins. Flavonoids have been extensively studied for their wide range of biological activities, such as antioxidant, anti-inflammatory, antidiabetic, antimicrobial, antimalarial, antiviral, anticancer, and neuroprotective effects.^23^ As a member of this family, fisetin also exhibits diverse activities, including senolytic properties,^19^ which extend beyond the common activities of flavonoids.^24–26^ However, fisetin is known for its promiscuity by targeting multiple pathways, which makes its molecular mechanisms complex.^24–26^ For example, fisetin can activate or suppress various molecular targets and pathways, enhancing antioxidant status, inducing apoptosis, and inhibiting proliferation, inflammation, and angiogenesis. Due to these complexities, traditional structure-based drug design approaches, which rely on targeting a predefined protein or biomolecule, are not well-suited for developing novel senolytic flavonoid analogs (FAs). To overcome this challenge, we employed a phenotypic drug discovery approach, leveraging senescent cell-based screening to evaluate various flavonoids for their senolytic activity.

We previously developed an efficient senotherapeutic screening platform based on senescence-associated beta-galactosidase (SA-β-gal) activity as a primary marker and phenotype of senescent cells.^8,18^ This platform, equipped with a high-content fluorescent imaging system, allows quick evaluation of both senolytics and senomorphics. Using this platform, we screened a focused library of various natural flavonoids, including flavones, flavonols, flavanols, anthocyanidins, and other subclasses, based on their ability to modulate the phenotype of SA-β-gal activity (Figures 1A and S1).

**Figure 1.**
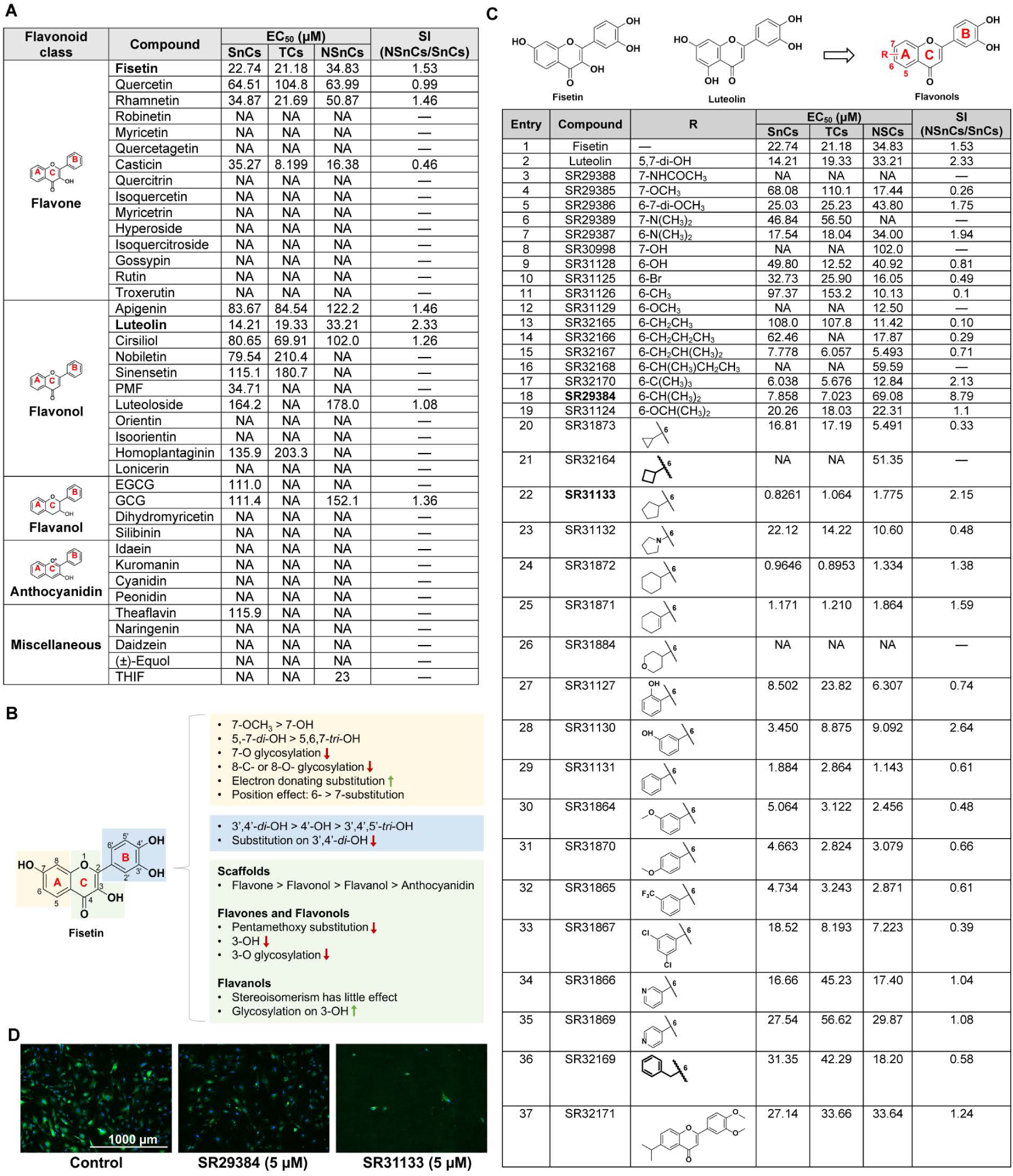
Discovery of novel flavonoid senolytics by drug screening and drug design. (A) Summary of the prelimianray senescence screening of flavonoid analogs of fisetin in senescent *Ercc1*^-/-^ MEFs. (B) Summary of the preliminary structure-activity relationship from the phenotypic drug screening of natural flavonoids. (C) Summary of the structural optimization for improved flavonoid analogs. (D) Representative images of the senescence assay showing SR29384 and SR31133 reduced SA-β-gal positive senescent *Ercc1*^-/-^ MEF cells after 48 hours of treatment. Blue fluorescence indicates nucleus stained with Hoechst 33324, and green fluorescence indicates SA-β-gal positive senescent cells stained with C_12_FDG. Images were captured using Cytation 1 at 4X magnification. Abbreviations: SnCs: Senescent *Ercc1*^-/-^ MEF cells; TCs: Total *Ercc1*^-/-^ MEF cells; NSnCs: Non-senescent WT MEF cells; SI: Selectivity index. EC_50_: the concentration that results in a 50% decrease in the number of cells compared with that of the control. NA: EC_50_ values are not available due to inactivity. PMF: 3′,4′,5′,5,7-pentamethoxyflavone; EGCG: (-)- epigallocatechin gallate; GCG: (-)-gallocatechin gallate; THIF: 3’,4’,7-trihydroxyisoflavone.

The senolytic activity of these compounds was assessed initially in *Ercc1*^-/-^ senescent mouse embryonic fibroblasts (MEFs). The ERCC1 protein is a subunit of the heterodimeric ERCC1/XPF DNA repair endonuclease essential for repairing multiple types of DNA damage. The *Ercc1*^-/Δ^ mice, with reduced expression of ERCC1, show increased sensitivity to DNA damage, elevated senescent cell burden, accelerated aging, and a shortened lifespan^27,28^. Accordingly, primary *Ercc1*^-/-^ MEFs with impaired DNA damage repair readily become senescent under oxidative stress by passage at 20% O_2_. The selectivity of each compound was determined using a selectivity index (SI), which compares the EC_50_ values in non-senescent proliferating MEFs to those in senescent *Ercc1*^-/-^ MEFs. The screening results showed that fisetin had a moderate senolytic EC_50_ value of approximately 22.74 μM and a low SI of 1.53 (Figure 1A), indicating the need for further optimization. Among the screened compounds, luteolin emerged as the most potent and selective natural flavonoid, at least in MEFs, with a senolytic EC_50_ value of 14.21 μM and an SI of 2.33 (Figure 1A).

### Preliminary SAR analysis

The primary objective of this phenotype-based senolytic screening was to gather preliminary structure-activity relationship (SAR) data, which is essential for the rational design of more potent and selective flavonoid senolytics. The majority of the flavonoids tested in this study share a common structural framework, consisting of two aromatic rings A and B linked by a central three-carbon ring C (Figures 1A and S1). Based on the phenotypic screening results, we conducted a detailed SAR analysis and established fundamental SAR guidelines for each moiety of the flavonoid structure (Figure 1B).

### Scaffold

In general, the flavonol scaffold demonstrated the most senolytic activity, followed by the flavone scaffold lacking the 3-hydroxyl group in the central C-ring. This trend was supported by the stronger senolytic activity of luteolin compared to quercetin. Flavanols and anthocyanidins were inactive even at higher concentrations, possibly due to the absence of the C2-C3 double bond in conjunction with the 4-oxo group in the C-ring. Compounds with other scaffolds, including isoflavones (daidzein and THIF), isoflavanes (equol), and flavanones (naringenin), were not effective as senolytics, suggesting the importance of the C2-C3 double bond and the C-4-oxo group in the C-ring for senolytic activity.

### Ring A

The SAR analysis revealed that the presence of a 5-hydroxyl group in the A ring decreased senolytic activity, as seen in the less potent quercetin compared to fisetin (Figures 1B and S1). Electron-donating substituents, such as methoxy groups in the A ring, were found to be favorable, evidenced by the increased potency of rhamnetin compared to quercetin. However, excessive methoxy substitutions, such as pentamethoxy groups, led to compound precipitation in cell culture media and reduced activity, as observed in nobiletin, sinensetin, and PMF. Quercetin outperformed quercetagetin, indicating that the presence of hydroxyl groups at positions 5 and 7 in the A ring is more favorable than three hydroxyl groups at positions 3, 5, and 7 in flavones. Any glycosylation in the A ring diminished senolytic activity, as seen with gossypin compared to quercetin. This pattern was also observed for flavonols. For example, glycosylation on the 7-OH of luteoloside substantially reduced senolytic activity compared to luteolin, the most potent flavonoid in the screening. Similarly, disaccharide glycosylation on 7-OH (lonicerin), 6-C-glycosylation (isoorientin), or 8-C-glycosylation (orientin) in the A ring of flavonol luteolin was detrimental to senolytic activity.

### Ring B

The number, position, and substitution of hydroxyl groups in the B ring are crucial for senolytic activity. This suggests that the presence of the 3’,4’-dihydroxyl groups in the B ring is essential (Figures 1B and S1), as having only one (as in apigenin < luteolin) or three (as in myricetin < quercetin) hydroxyl groups was less desirable. Although not definitive, the data also suggest that substitution on the 3’,4’-dihydroxyl groups in the B ring might decrease activity, as observed with troxerutin.

### Ring C

The presence of the 3-hydroxyl group in the C ring was found to be unfavorable for senolytic activity. This was demonstrated by the reduced activity of flavone quercetin compared with flavonol luteolin. Further SAR analysis of flavones revealed that glycosylation on the 3-hydroxyl group resulted in diminished activity. For example, various saccharide substitutions on the 3-OH of quercitrin, isoquercetin, hyperoside, isoquercitroside, and rutin decreased their activity compared with quercetin. This may be due, at least in part, to the steric factors of the saccharide substituents. In contrast, for flavanols, glycosylation on the 3-OH group increased activity, as evidenced by the slightly improved senolytic activity of EGCG and GCG compared with cyanidin. Unlike flavones or flavonols, the lack of conjugation between rings A and C in flavanols EGCG and GCG may tolerate the steric hindrance of the galloyl moiety on the 3-OH, thus increasing their senolytic activity compared with dihydromyricetin. Additionally, for the flavanol scaffold, stereoisomerism appeared to have little effect on senolytic activity, as the activities of GCG and EGCG are comparable.

### Drug design for enhanced flavonoid senolytics

The SAR analysis underscored the importance of the flavonol scaffold in developing senolytic FAs, with luteolin emerging as the most potent and selective compound in the preliminary screening. Based on the SAR information, we hypothesized that the absence of a hydroxyl group at the 3-position in the C ring, combined with the presence of unsubstituted 3’,4’-dihydroxyl groups in the B ring, could be responsible for the enhanced senolytic activity and selectivity observed in luteolin. We conducted an iterative process of structural optimization, encompassing drug design, chemical synthesis, biological evaluation, and further SAR analysis, to identify novel senolytic FAs with improved efficacy and selectivity.

In total, we designed, synthesized, and tested 35 novel flavonol analogs for senolytic activity in senescent *Ercc1*^-/-^ MEF cells and for selectivity against non-senescent WT MEF cells (Figure 1C). The majority of these analogs demonstrated significantly enhanced senolytic activity compared to fisetin or luteolin. These results also highlighted the superiority of the flavonol scaffold over the flavone structure. The SAR analysis provided additional insights consistent with the preliminary findings. For instance, compounds featuring electron-donating groups on the A ring (compounds 4-7) were more active than those with electron-withdrawing groups (compound 3). Moreover, substituents at the 6- position on the A ring were generally more favorable for senolytic activity than those at the 7-position (compound 7 > compound 6, compound 9 > compound 8). These observations led us to design additional FAs with 6-substitutions, including aliphatic, alicyclic, and aromatic groups.

Among the acyclic aliphatic substituents, bulky tert-butyl (compound 17) and isopropyl (compound 18 or SR29384) groups notably enhanced the senolytic activity of the flavonols. The isopropyl-substituted compound SR29384 emerged as the most selective senolytic agent among all designed FAs. The introduction of five- or six-member all-carbon alicyclic substituents (compounds 22, 24, and 25) further boosted the senolytic activity. Notably, SR31133 (compound 22) exhibited the highest potency, with a senolytic EC_50_ value of 0.8 µM, resulting in a more than 27-fold improvement in senolytic activity compared to fisetin. However, it had reduced specificity for senescent cells compared to SR29384. Substituted phenyl (compounds 27-33), pyridine (compounds 34 and 35), or benzyl (compound 36) groups at the 6-position of the A ring were tolerated but exhibited lower selectivity. Furthermore, the presence of free 3’,4’-dihydroxyl groups in the B ring proved critical for senolytic bioactivity, as substituting these hydroxyl groups (compound 18 or SR29384) with methoxy groups (compound 37) significantly reduced both senolytic activity and selectivity.

Collectively, our phenotype-guided drug design efforts led to the successful discovery of two promising senolytic FAs: SR29384, the most selective, and SR31133, the most potent. In comparison, fisetin has a diphenylpropane flavone scaffold with four hydrogen bond donors and a 3-hydroxy group in the central ring, whereas the two FAs possess a flavonol scaffold lacking the 3-hydroxy group and have only two hydrogen bond donors. As a result of these structural modifications, SR29384 and SR31133 exhibited 2.9- and 27.5-fold greater senolytic activity than fisetin, respectively (Figure 1C). Beyond the SA-β-gal activity (Figure 1D), the senolytic efficacy of these compounds was further validated by assessing their effects on key senescence and SASP markers by RT-qPCR. Treatment with SR29384 and SR31133 resulted in significant downregulation of *Cdkn2a* (p16^INK4a^), *Cdkn1a* (p21^Cip1^), and the SASP factor IL-6 compared to fisetin (Figure S2C). Regarding the physicochemical and molecular properties, both FAs exhibited increased lipophilicity, with WLogP values of 3.99 for SR29384 and 4.53 for SR31133, compared to 2.28 for fisetin (Figure S2A). The two FAs have lower topological polar surface area (TPSA) values than fisetin, each less than 90 angstroms squared, suggesting better permeability of the blood-brain barrier (BBB).^29^ Moreover, both FAs demonstrated better computational molecular properties compared to fisetin in terms of drug score, drug-likeness, LD_50_ values, and predicted toxicity, including a substantially reduced probability of carcinogenicity risk (Figure S2A, Supplementary Table 1E).

### Broad spectrum senolytic activity of the optimized flavonoid analogs

To further validate the senolytic activity of the newly designed FAs SR29384 and SR31133, we assessed their potential across a range of senescent cell models induced by various inducers. These models include WT MEF cells subjected to oxidative (H_2_O_2_) and genotoxic (etoposide) stress, human IMR90 fibroblast cells induced to senescence by genotoxic stress (etoposide), and human umbilical vein endothelial cells (HUVEC) undergoing replicative senescence. The results showed both SR29384 and SR31133 effectively reduced the number of C_12_FDG-positive senescent cells in these senescent cell models (Figures S2B and S2D). Therefore, we selected SR29384, the most selective, and SR31133, the most potent, for further evaluation.

### Analysis of senolytic activity *in vivo*

To assess the *in vivo* senolytic potential of SR29384 and SR31133, we used acute treatment of old WT C57BL/6J mice (27 months) in which fisetin would not be expected to be significantly effective according to previous studies.^19^ Instead of the 100 mg/kg dose used in earlier fisetin studies, the old wild-type C57BL/6J mice were acutely treated with a lower dose of 20 mg/kg of body weight of the compounds for five consecutive days. Mice were sacrificed two days after the last administration and tissues were collected for analysis (Figure 2A). The RT-qPCR analysis of different tissues found that fisetin had a marginal effect on senescence and SASP. However, the two FAs, SR29384 in particular, significantly decreased the expression of senescence markers *Cdkn2a*, and *Cdkn1* as well as multiple SASP factors, such as *IL6*, *IL1β*, *Mcp1*, and *Pai1*, in different tissues including kidney, brain, lung, spleen, fat, and muscle (Figures 2B-E, S3, and S4).

**Figure 2.**
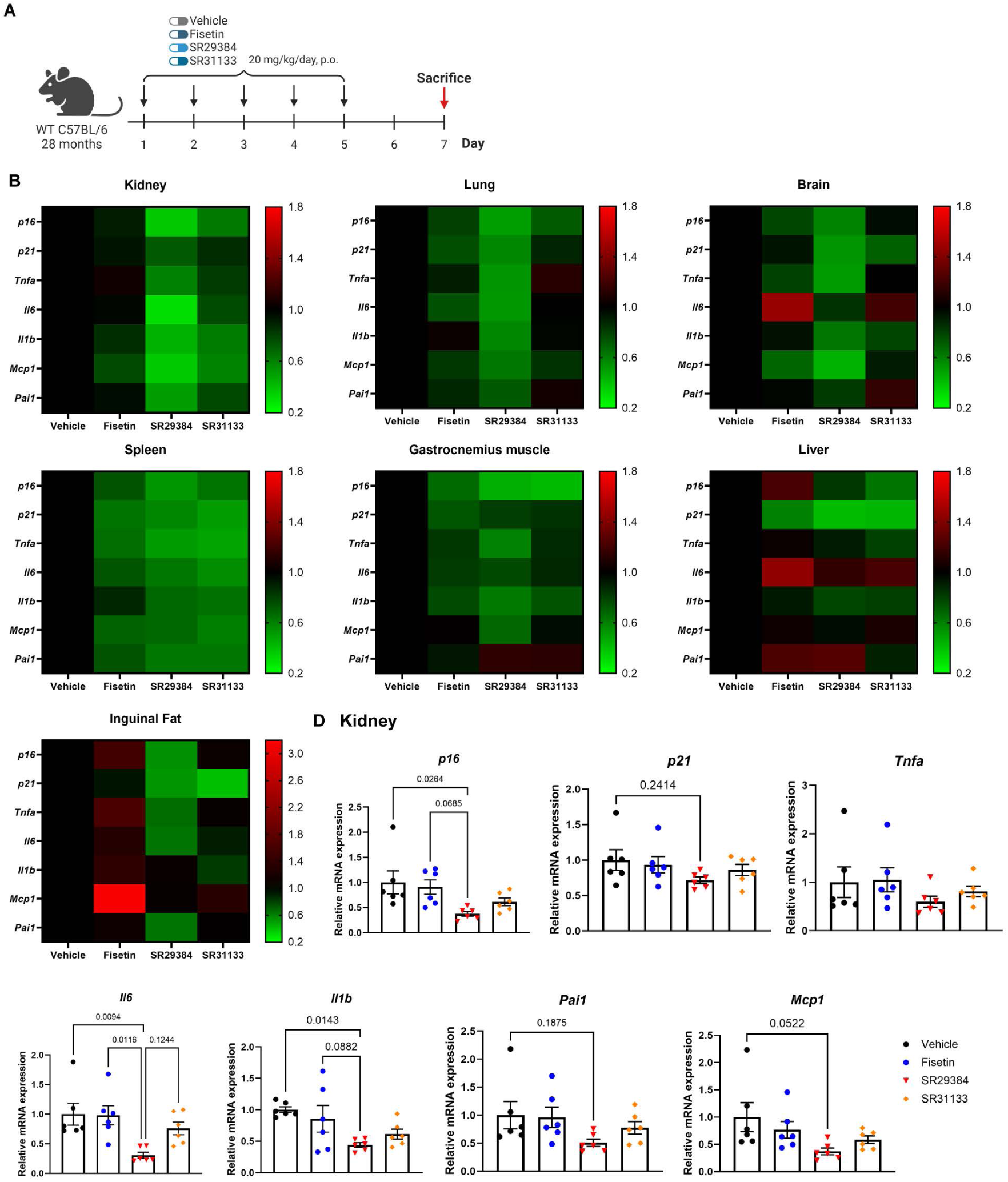
Acute treatment with the flavonoid analogs reduced tissue senescence in naturally aged mice. (A) At 27-months of age, wild-type C57BL/6J mice were administered vehicle, fisetin, SR29384, or SR31133 for five consecutive days by oral gavage at a dosage of 20 mg/kg/day. Mice were sacrificed two days after the final dose, and tissues were collected for analysis. (B) Heatmap showing mRNA expression changes of senescence and SASP factors in various tissues following treatment with vehicle, fisetin, SR29384, or SR31133. (C) RT-qPCR analysis of kidney tissue showing relative mRNA expression of senescence and SASP markers across treatment groups. Error bars represent the standard error of the mean (SEM) for n = 6 mice per group.

### Chronic treatment of SR29384 extends healthspan in accelerated aging mice

The most effective fisetin analog, SR29384, identified in the acute mouse study, was selected for testing in the *Ercc1*^-/Δ^ progeria mouse model of accelerated aging to determine its effect on tissue senescence and healthspan. *Ercc1*^-/Δ^ progeria mice were treated with 20 mg/kg of SR29384 for five weeks *via* oral administration (Figure 3A). Health assessments were conducted weekly to score age-related symptoms, including tremor, kyphosis, dystonia, ataxia, gait disorder, and body condition. Results showed that SR29384 effectively suppressed the composite score of aging symptoms, particularly dystonia and ataxia (Figure 3B). Although there were no significant changes in the composite score for aging symptoms by the end of the healthspan evaluation at week 16, gene expression analysis in tissues showed a significant reduction or trend towards reduction in senescence and SASP factors across multiple tissues, including the kidney, liver, lung, and brain (Figure 3C). These results further confirmed the promising senolytic activity of the optimized flavonoid analog SR29384.

**Figure 3.**
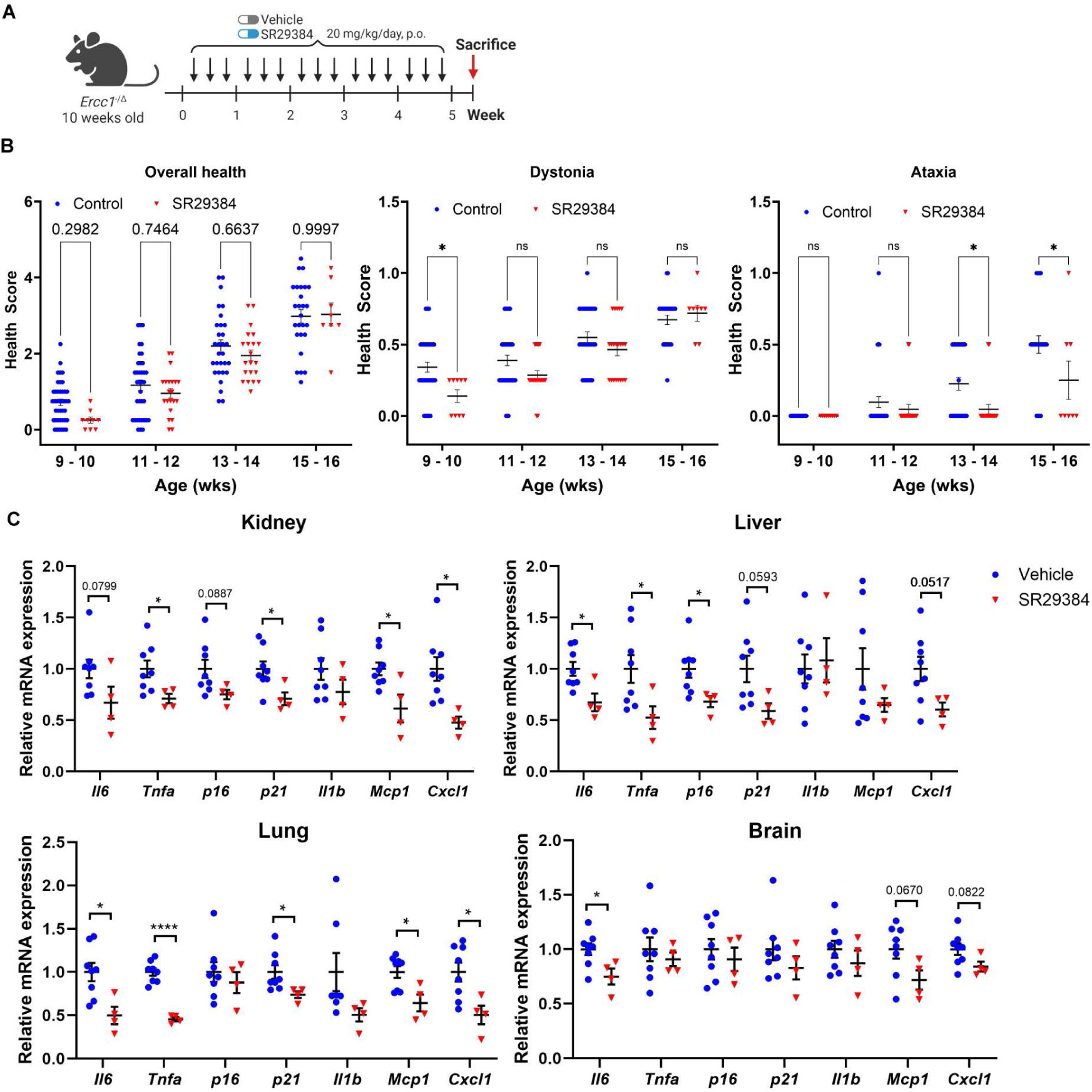
SR29384 extended healthspan and reduced tissue senescence in *Ercc1*^-/Δ^ progeria mice. (A) At 10 weeks of age, *Ercc1*^-/Δ^ progeria mice were treated with SR29384 or vehicle by oral gavage for five consecutive weeks. Mice were sacrificed two days after the final dose, and tissues were collected for further analysis. (B) Health assessments were performed weekly to score diverse age symptoms including tremor, kyphosis, dystonia, ataxia, gait disorder, and body condition. (C) RT-qPCR analysis of senescence and SASP markers in tissues collected from vehicle-treated and SR29384-treated mice. Error bars represent SEM for n = 8 (vehicle) and n = 4 (SR29384).

Collectively, these *in vivo* studies demonstrated the significant therapeutic potential of SR29384 as a senolytic agent, capable of reducing senescence markers, alleviating age-related symptoms, and extending healthspan in mice.

### Prediction of mechanistic processes underlying senolytic effects of FAs using transcriptomics and *in-silico* modeling

To elucidate the mechanisms underlying the improved senolytic activity of FAs SR29384 and SR31133, we first examined whether differences in antioxidant capability could be a contributing factor. Using the DPPH radical scavenging assay^30^, we compared the antioxidant activity of fisetin and the two analogs. The results showed that all compounds exhibited similar free radical scavenging activity, as demonstrated by their comparable IC_50_ values (Figure S2E). This finding suggests that the enhanced senolytic activity of SR29384 and SR31133 is not due to differences in antioxidant properties.

Next, we conducted bulk RNA-Seq analysis to identify gene expression changes and key signaling pathways in non-senescent (NS) and etoposide-induced senescent (S-Eto) cells following treatment with fisetin, SR29384 and SR31133 (Figure S5A-B). We used the SenMayo gene set^31^ enrichment as an independent validation of senescence induction in the S-Eto cells. We also observed a reduction of SenMayo enrichment in the S-Eto cells treated with fisetin (NES: −0.68), SR29384 (NES: −1.05), and SR31133 (NES: −1.27) (Figure S5C, Source Data 2). To better understand the underlying biological processes, we carried out Gene Set Enrichment Analysis (GSEA) using the Hallmarks database to identify pathways associated with the differentially expressed genes (DEGs). The top pathways activated by SR29384 and SR31133 were associated with hypoxia, p53 signaling, and multiple stress signaling pathways, while downregulated processes included G2M transition, E2F targets and DNA replication processes (Figure S5D, Source Data 2).

To further explore the possible molecular mechanisms, we used a sequence of *in silico* structure-based machine learning models and network analyses. First, we used PathwayMap, a deep self-normalizing convolutional neural network (DSNNN) machine learning model, to predict compound-pathway interactions.^32^ In addition to the two FAs, we included 7 reported senolytics for comparison, including fisetin and luteolin. PathwayMap probability scores for interactions with human biological pathways reveal that SR29384 and SR31133 had highly similar interaction signatures (Figures S6A-C). In contrast to fisetin, which clustered closely with quercetin, SR29384 and SR31133 did not have a strong interaction probability for human longevity pathways. Instead, SR29384 and SR31133 cluster with luteolin and dasatinib, which were predicted to strongly interact with pathways associated with G2M transitioning, oxidative stress, cellular senescence, ER stress, TP53 signaling, ESR signaling, and caspase activation. Additional data for pathway enrichment using the PathwayMap algorithm is provided in Supplementary Tables 1B-D.

These preliminary interaction probabilities prompted the identification of specific molecular drug targets. We used structural features of the drugs to predict such drug-protein interactions in the human druggable proteome using SwissTargetPrediction, a similarity-based algorithm trained with curated experimental ligand-receptor binding assays.^33^ As negative controls, we included 10 compounds that showed no senolytic activity (“NC” in Supplementary Table 1A). This NC filtering excluded targets shared with non-senolytic FAs, and reduced biases introduced by the scaffold. After filtering, SR29384 and SR31133 were found to target 49 and 44 proteins, with 29 overlapping targets (Figure 4A). To understand the biological functions of these targets, we constructed clustered protein-protein interaction (PPI) networks of the shared and exclusive targets for the two candidates. Targets shared between SR29384 and SR31133 grouped into three major clusters: p53 signaling and cell cycle regulation, estrogen receptor (ESR) activity, and PARP signaling. In addition to these shared targets, nodes exclusive to SR29384 are associated with inflammatory signaling and ECM remodeling. Conversely, SR31133 exclusive targets were enriched for histone modification processes, particularly HDAC family proteins (Figure 4B). These enrichment trends for the networks corroborated our previous findings from the DSNNN-based PathwayMap analysis. The shared target network was enriched for pathways associated with G2M transitioning, ESR signaling, stress response, and apoptosis regulation. The SR29384 exclusive target network enriched for immune cell activation, ECM disassembly, and response to hypoxia. The SR31133 exclusive target network primarily enriched for deacetylation and transcriptional regulation (Figure S6D). Interestingly, after NC filtering, SR29384 and SR31133 shared more targets with each other than with their parent compounds fisetin or luteolin (Figure S6A). A similar clustering and enrichment approach distinguished the two candidate FAs from luteolin and fisetin by exclusive targets enriched in processes associated with inflammatory signaling, unsaturated fatty acid metabolism, and apoptosis (Figures S7B, S8A-B). Further details on SwissTargetPrediction scores are available in Supplementary Tables 2A-E.

**Figure 4.**
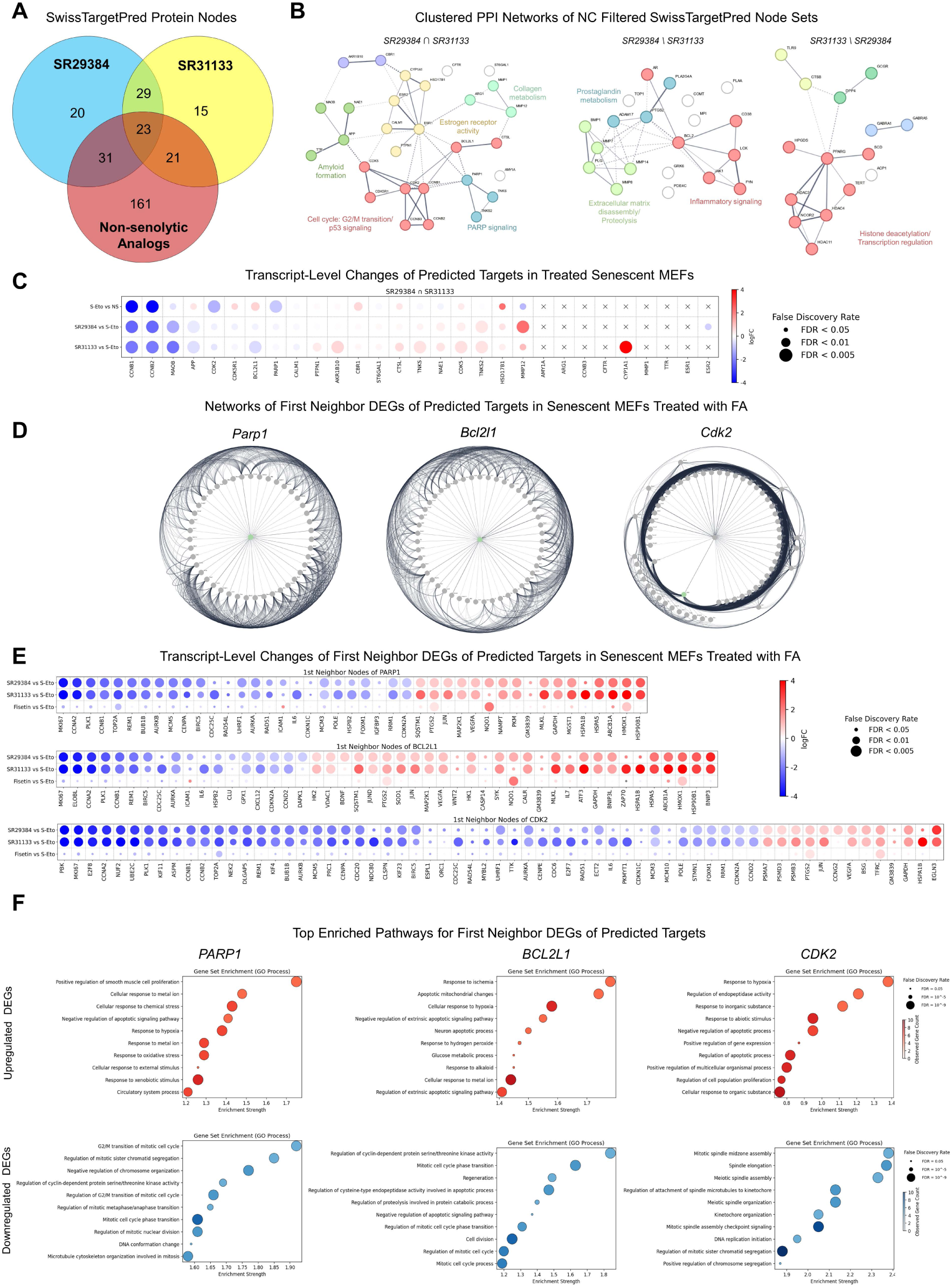
Prediction of direct molecular targets of top candidate FAs using ligand structure similarity and network-perturbation. (A) Venn diagram depicting overlaps between targets for SR29384, SR31133 and unique targets for negative control FAs (NC) using SwissTargetPrediction scores. (B) k-means clustered STRING PPI maps constructed from shared and exclusive targets for SR29384 and SR31133. Target sets were filtered to exclude targets shared with NCs. Nodes are clustered based on degree centrality. The top enriched GO-terms for each cluster are used as labels. (C) Bubble plot grid depicting changes in mRNA expression of predicted shared targets in etoposide induced senescence in MEFs, and senescent MEFs treated with SR29384, and SR31133. (D) Radial diagram of 1^st^ neighbor node DEGs of PARP1, BCL2L1, and CDK2. The target nodes are represented in green; hub genes within the sub-networks are in the center. (E) Bubble plot grid depicting changes in mRNA expression of 1^st^ neighbor node DEGs of PARP1, BCL2L1, and CDK2 in senescent MEFs treated with SR29384, SR31133, and Fisetin. (F) Ranked bubble plot depicting top 10 enriched pathways (GO Biological Processes) for upregulated and downregulated (log_2_FC > |1|, FDR< 0.05) 1^st^ Neighbor DEGs for PARP1, BCL2L1, and CDK2 in senescent MEFs treated with SR29384 and SR31133. (B) & (E) Connecting edges are defined as level of interaction confidence (probability > 0.9). (C) & (E) Genes are sorted by log_2_FC values in the SR29384 vs S-Eto dataset. Color and sizes of bubbles correspond to a set scale for log_2_FC and FDR values; “x” symbols represent genes not detected in dataset (CPM ≤ 0.5). (E) Bubble color intensity and sizes correspond to number of observed genes in GO term, and FDR values (< 0.05).

While these sets of targets provided us with an initial array of mechanistic targets, they required experimental support. For this, we first used detectable mRNA levels (CPM > 0.5) of each target in our RNA-Seq experiments as a proxy for cellular expression of targets. Nodes like ESR1, ESR2 and TTR that did not meet this criterion were excluded (Figure 4C). We hypothesized that true protein targets of the candidate drugs would not exhibit significant expression changes at the mRNA level upon treatment. However, for selective targeting of senescent cells, we expected that genes involved in senescence-related pathways would be altered during the senescence program. Using this rationale, we filtered for targets with log_2_FC > |0.5| and FDR < 0.05 in the non-senescent (NS) vs etoposide-induced senescent (S-Eto) MEF RNA-Seq datasets, with no significant change in expression between SR29384 or SR31133 treatment and the S-Eto groups (Figure 4C, S9A, Source Data 1). This analysis identified PARP1, CDK2, CDK5R1, HSD17B1, and BCL-xL (transcript name BCL2L1) as potential direct targets shared by the candidate FAs. Similarly, exclusive targets exhibiting this signature were TOP1, COMT, GRK6 for SR29384, and TERT, HDAC4, HDAC11 for SR31133.

If these shortlisted proteins are direct targets of the candidate FAs, they should have a substantial impact in perturbing their local gene network neighborhoods. For this, we isolated subnetworks consisting of nodes from 1^st^ neighbor interactor genes for each of the query proteins from a parent network of all DEGs from the FA-treated RNA-Seq datasets. Based on the number of DEGs in the local neighborhood and perturbation signatures, we determined PARP1, BCL-xL, and CDK2 as the best potential targets shared by the two FAs (Figure 4D, S9B). Indeed, the expression level changes of DEGs in their 1^st^ shell neighborhoods showed significant up- and downregulation without any changes of the targets themselves at the transcript level (Figure 4E). Interestingly, there was minimal change detected for these genes in fisetin-treated senescent MEFs. Conversely, there was a strong resemblance in magnitude and directionality of differential expressions of the DEGs between SR29384 and SR31133 treatments. Furthermore, these perturbations in shared FA target neighborhood DEGs functionally enriched many processes that mirror ontologies from both our RNA-Seq analyses and structure-based predictions (Figures S5D, S6D). Upregulated neighbor DEGs of PARP1, BCL-xL, and CDK2 showed strong enrichment for apoptosis, regulation of cell proliferation, response to hypoxia and oxidative stress. Conversely, downregulated neighbors enriched processes associated with DNA damage response and repair, cell division, G2M cell cycle transitioning, and anti-apoptotic processes (Figure 4F).

Additionally, individual queries of neighbor nodes using StringDB suggest the directional signatures of the DEGs to be associated with inhibited antagonists and upregulated agonists/co-expressors of PARP1, BCL-xL, and CDK2 proteins. This suggests that SR29384 and SR31133 may act as inhibitors of these target proteins. To validate this, we performed molecular docking simulations for the three top target proteins to assess energetic and conformational feasibilities of the candidate FAs to function as inhibitor compounds. Each set of ligands docked into pre-defined inhibitory conformational spaces include a collection of random decoy molecules (DUDs) as technical negative controls, a native substrate, two FAs with no experimental senolytic activity as experimental negative controls, two natural senolytic FAs (fisetin and luteolin), and three standard inhibitors of each protein. A list of crystal receptor structures, native substrates, positive controls, and relevant references is included in Supplementary Table 3A.

In case of the PARP1 catalytic cavity, an ideal competitive inhibitor should bind with a stronger binding affinity than the activating co-factor NAD+. The best docked ligand conformations for SR31133, SR29384, and luteolin, but not fisetin, have stronger interactions than native substrates or non-senolytic FAs within the receptor cavity (Figures 5A-B). This was further elucidated by the ligand orientations within the binding cavity, and the interaction characteristics of ligand atoms with surrounding amino acid residue. Fisetin, luteolin, SR29384, SR31133, and the top scoring standard talazoparib shared key favorable hydrogen bonding events with residues including GLY863, GLY904, GLN759, TYR889, TYR896, and SER904. Additionally, a set of hydrophobic interactions with TYR889, TYR896, ALA898, and TYR907 contributed to the favorable binding profiles. However, two donor-donor clashes with HIS862 and MET890 residues inhibit fisetin from strongly binding to the cavity. However, the benzamide core of talazoparib established a strong halogen bond with GLU988, a feature absent in the flavonoid analogs (Figure 5C).

**Figure 5.**
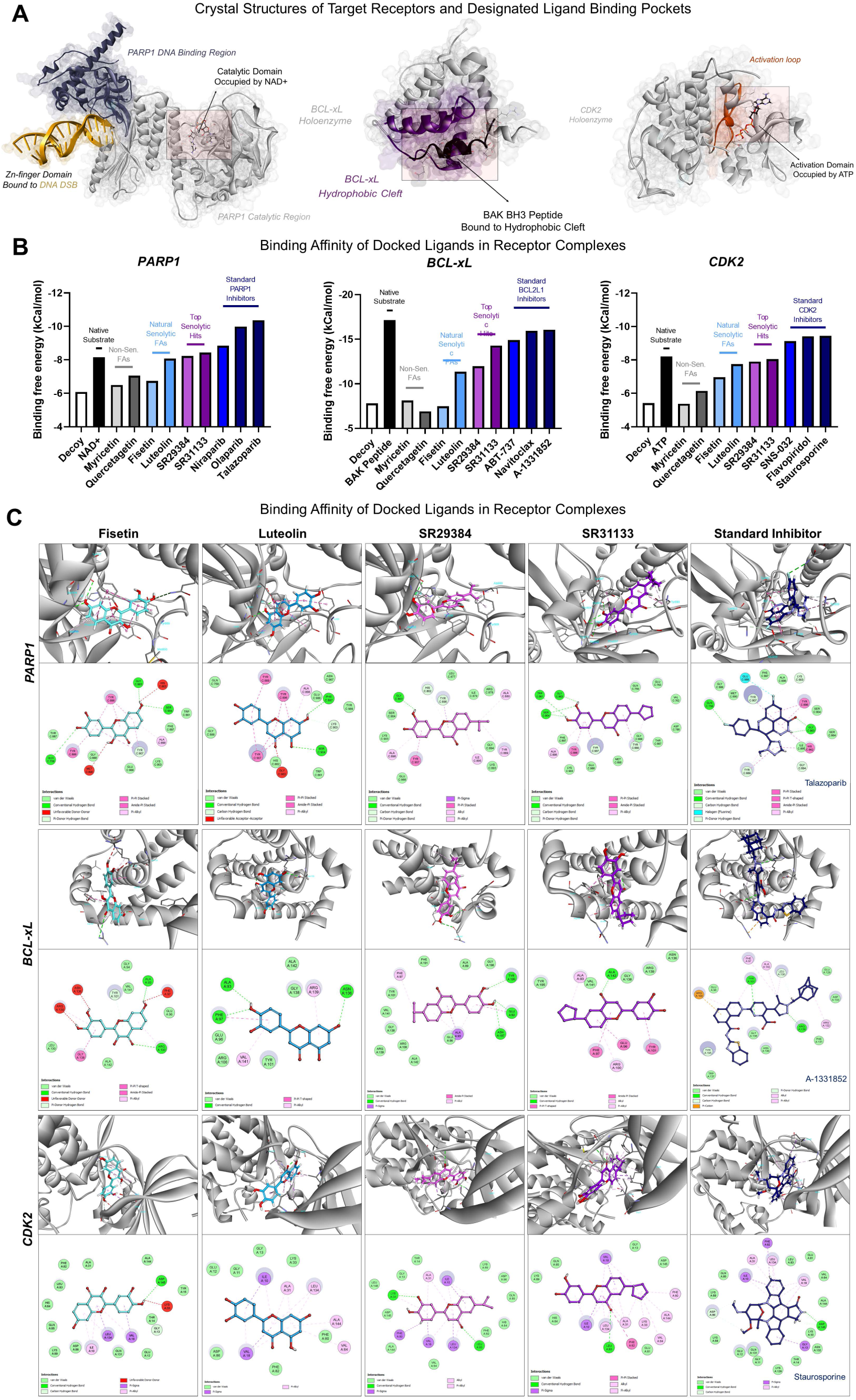
Flexible molecular docking results of predicted target proteins with candidate FAs and control compounds. Native substrate or ligand and known inhibitor or activator drug for each protein target was used along with a randomly generated structural decoy/DUD molecule. (A) 3D molecular model depictions of the receptor protein crystal structures used for docking simulations with surrounding native templates used for search area definition. (B) Binding free energy scores of best fitting conformers for each ligand within the selected receptor binding pockets; lower binding free energies correspond to stronger ligand-receptor interactions. (C) 3D and 2D interaction plots of ligands in the binding pockets with interacting pocket amino acid residues. Color identifiers correspond to panel (B); the best scoring standard inhibitor for each target was represented; ligand-residue interaction legends are provided under the 2D plots.

When docked into the navitoclax/BH3 binding site of BCL-xL, the native BAK peptide had a strong binding affinity threshold that is overcome by SR31133 and the standard compounds, but not SR29384 or the other FAs. SR31133 binding to BCL-xL hydrophobic cleft was primarily driven by a set of weak hydrophobic and strong electrostatic amide-pi bonds with residues like ALA93, GLU96, PHE97, ARG100, TYR101, and ALA142. In addition to some of these shared hydrophobic residues, the binding affinity for the standard A-1331852 was substantially enhanced by an established cationic bond with ARG100. SR29384 only shared one strong amide-pi bond with SR31133, while luteolin and fisetin had none. Fisetin-BCL-xL binding was also hampered by multiple donor-donor clashes with its functional group.

Finally, in the CDK2 activation domain, competitive inhibition of ATP binding will disrupt the cyclin binding and phosphorylation in the activation loop. Both SR29384 and SR31133 were predicted to outcompete the native substrate and bind to the ATP binding cavity. The strongest inhibitor for CDK2 is staurosporine, which relies on an array of weak hydrophobic interactions with ILE10, GLY13, VAL18, ALA31, PHE82, and LEU134. Luteolin, SR29384 and SR31133 shared many of these hydrophobic interactions, even establishing more interactions with LYS33, VAL64, PHE80, and ALA144. The best docked pose of fisetin had the lowest number of hydrophobic interactions, with unfavorable donor-donor clashes again making its ability to inhibit CDK2 energetically unfavorable.

Taken together, we have utilized machine learning algorithms, *in silico* structure-based prediction models, and network perturbation of expressional signatures in senescent MEFs to identify potential targets underlying the senolytic activities of SR29384 and SR31133. Our findings predict PARP1, BCL-xL, and CDK2 as key protein targets, with both FAs demonstrating distinct network perturbation of the local expressional neighborhoods and strong binding capacities, overcoming limitations of the parent compound (Figure 6). These results suggest that the candidate FAs are not only potential chemical antagonists of these proteins, but also target distinct biological processes, including DNA damage repair, hypoxic stress, and anti-apoptotic pathways, providing valuable insights into potential senolytic mechanisms.

**Figure 6.**
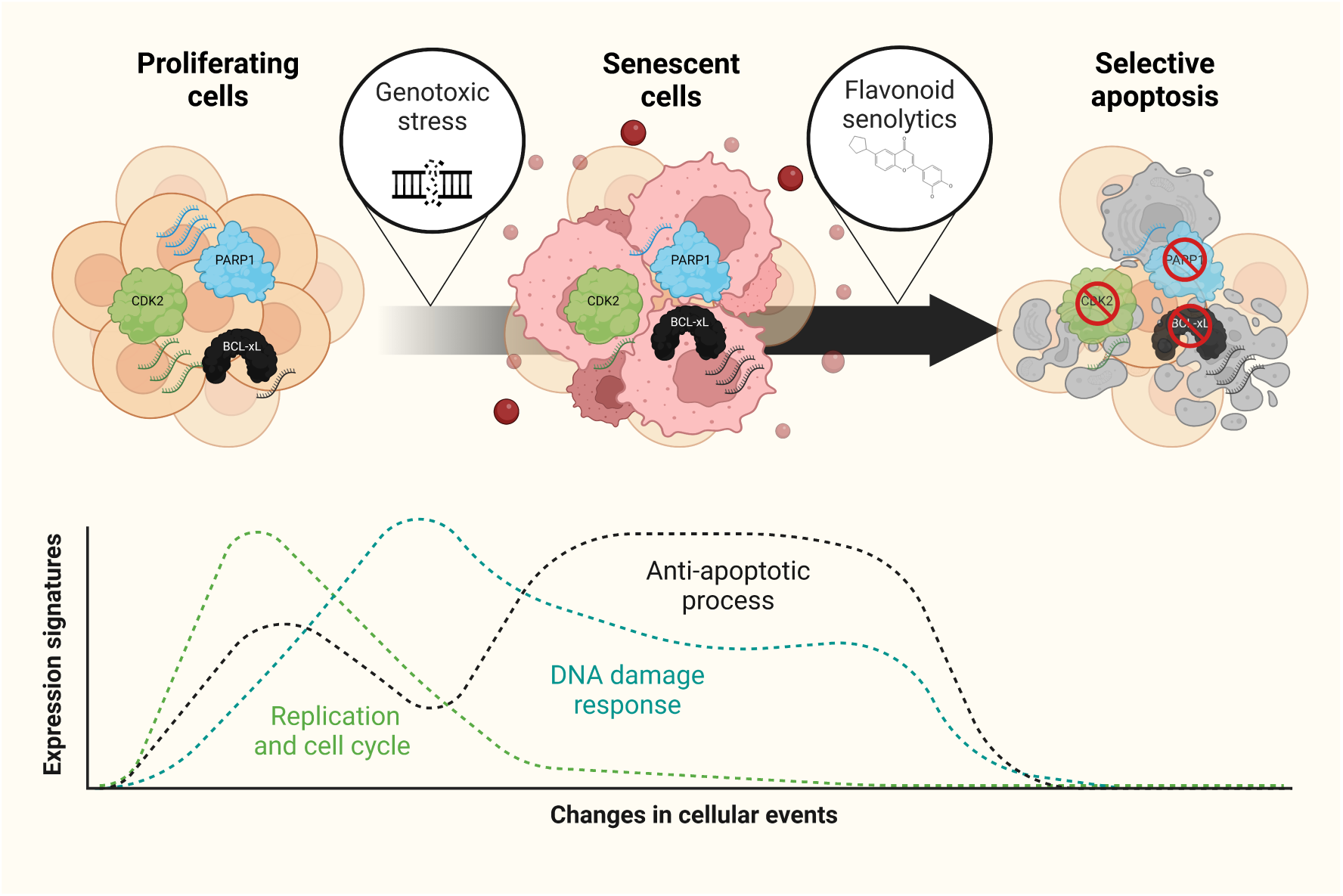
Proposed mechanism of flavonoid senolytic mediated apoptotic clearance of senescent cells. Senescent cells, induced by genotoxic stress, exhibit elevated levels of key proteins involved in cell cycle arrest, DNA damage response, and anti-apoptotic pathways, such as PARP1, CDK2, and BCL-xL. Flavonoid senolytics (SR29384 and SR31133) selectively target these proteins in senescent cells, disrupting their anti-apoptotic processes and promoting apoptosis, while sparing proliferating cells. The lower panel shows changes in cellular expression signatures, where replication and cell cycle processes decline, DNA damage response escalates, and anti-apoptotic signaling is elevated in senescent cells, which is later suppressed by the senolytic action of flavonoid compounds, leading to cell death.

## Discussion

Senolytic drug discovery has been challenging due to the limited identification of specific senolytic drug targets, with BCL-2/BCL-xL being one of the few convincingly identified. In this study, we aimed to develop novel flavonoid-based senolytics starting from fisetin, a compound with largely unknown targets. To overcome this challenge, we employed a phenotypic drug discovery approach, which focuses on identifying compounds based on their functional effects rather than their interactions with predetermined targets.^34–36^ This strategy allowed us to obtain valuable structure-activity relationship (SAR) insights, which guided the subsequent drug design process. As a result, we successfully identified two novel flavonoid analogs, SR29384 and SR31133, which demonstrated significantly improved senolytic activities compared to fisetin. Additionally, our initial screening also identified luteolin, another natural flanonoid, as a more potent natural senolytic than fisetin. Consistent with this, several studies showed that luteolin can supress H_2_O_2_-induced senescence in HEI-OC1 cells^37^ and TNF-α-induced senescence in human nucleus pulposus cells^38^. Moreover, luteolin is the most concentrated flavonoid in Haenkenium extract, which was shown to reduce senescence induced by aging or chemotherapy.^39^

During our senescent cell-based phenotypic drug screening, we tested various flavonoids, including fisetin and quercetin, for their senolytic activities. Our results found that neither fisetin nor quercetin exhibited significant senolytic activity at lower concentrations (Figure 1A). Specifically, fisetin showed only weak senolytic activity in senescent *Ercc1*^-/-^ MEFs and minimal senolytic effects in all other tested cell types, including senescent IMR90 cells, WT MEFs, and HUVECs (Figure S2B). Some studies have reported on the potential of fisetin as a senolytic agent. For example, Yousefzadeh et al. reported that fisetin reduced the burden of senescent cells and extended lifespan in naturally aged and accelerated aging mouse models.^19^ However, fisetin is not a pan-senolytic, displaying varying senolytic activity on different cell types and experimental conditions. For example, fisetin was found to kill senescent MEFs, but not sensecent human melanoma SK-MEL-103 cells.^40^ Additionally, fisetin, along with quercetin, was found to minimally reduce *p16*^Ink4a^+ fibroblasts isolated from fibrotic mouse lungs^41^, suggesting the senolytic activity of fisetin is context-dependent. It is also important to note that fisetin and other flavonoids have a wide range of cellular and systemic biological effects, including anti-inflammatory, antioxidant, anticancer, antidiabetic, and neuroprotective activities,^23,24,26,42^ which may contribute to their gerotherapeutic benefits independent of their specific and direct senolytic properties.

Our SAR analysis revealed key structural features that contribute to senolytic activity of flavanoids. The most potent flavonoid structures were characterized by a 3’,4’-O- dihydroxyl group in the B-ring, a 2,3-double bond combined with a 4-keto group in the C-ring, and electron-donating groups in the A-ring. Notably, removing the 3-hydroxyl group from the central ring of the flavone scaffold, resulting in a flavonol structure, significantly improved senolytic activity. This is examplified by luteolin, a flavonol that demonstrated higher senolytic activity compared to the flavone fisetin. Consistent with these SAR findings, the two flavonoid analogs (FAs), SR29384 and SR31133, share the flavonol scaffold with luteolin rather than the flavone scaffold with fisetin. The SAR information obtained in this study not only underscores the potential of flavonoids as senolytic agents if appropriately modified, but also provides valuable insights for the future design and discovery of more potent senolytic compounds within the flavonoid family.

Our study also found that fisetin and the two designed FAs have similar antioxidant activities, suggesting that the diiferences in their senolytic efficacy are less likely due to variations in their ability to scavenge free radicals. It is possible that the strucural differences between the compounds enhance their binding affinity and/or modulation of specific targets related to senolysis, contributing to the improved senolytic activity of SR29384 and SR31133. These structural modifications may improve the compounds’ ability to selectively induce apoptosis in senescent cells or interfere with specific signaling pathways involved in senescent cell death.

Using machine learning and structural similarity-based target prediction algorithms filtered by non-senolytic flavonoids and verification in our transcriptomic datasets allowed us to identify a set of potential drug targets for SR29384 and SR31133. Among these, PARP1, BCL-xL, and CDK2 were the most pronounced target proteins in our model. PARP1 is widely known in terms of cancer cell survival and DNA damage response processes. PARP inhibitors have been shown to induce a transient “senescence-like” phenotype in cancer cells,^43,44^ which makes them vulnerable to secondary modes of clearance, such as BCL-2 and BCL-xL inhibitors.^44,45^ Paradoxically, non-tumor senescent cells consistently have dampened expression and activity of PARP1 at the transcript and protein levels.^46^ In accordance, MEFs treated with DNA damage agents in our experiment exhibited reduced expression of PARP1 and downstream targets. Suppression of PARP1 with inhibitors have been suggested to reduce inflammatory signaling and induce apoptosis by bypassing PARP-mediated DNA SSB repair and trapping PARP on cytotoxic DNA complexes.^47^ Both SR29384 and SR31133 target PARP1 and significantly blunts neighboring nodes. This potentially exploits a therapeutic vulnerability in senescent cells by disrupting detection of DNA damage and promoting apoptosis. Similar to PARP1, evidence related to crosstalk between CDK2, senescence, and apoptosis components are conflicting. While cellular protein inhibitors of CDK2 such as p21^Cip1^ and p27^Kip1^ are associated with inducing senescence in cancer cells, small molecule inhibitors of CDK1/CDK2 such as flavopiridol and staurosporine show strong caspase cleavage and apoptosis induction via regulation of E2F and BCL-xL/MCL-1.^48,49^

Another key signature of senescent cells is resistance to apoptosis through upregulation of BCL-2 family proteins. BCL-2 and BCL-xL also have been suggested to directly participate in cell cycle arrest of senescent cells.^50^ Expectedly, in our study, etoposide treated cells had significant upregulation of BCL-xL. From our model and network analysis, SR29384 and SR31133 are predicted to be moderate to strong competitive inhibitors targeting BCL-xL and its functional processes. Based on expressional signatures and molecular interaction modelling, we speculate BCL-xL inhibition to be a key apoptotic mechanism in combination with PARP1 and CDK2 inhibition.

There are also substantial connections among the different targets. For instance, PARP1 is directly phosphorylated by CDK2. PARP1 inhibition or caspase-mediated cleavage has also been suggested to direct DNA-damaged cells towards apoptotic processes rather than necrotic cell death by disrupting PARP1-BCL-2 and PARP1-BCL-xL interactions.^51,52^ BCL-xL also acts in tandem with CDK2 to inhibit apoptosis, while inhibitors for both synergize and induce apoptotic cell death.^53^ A key process enriched throughout the study was hypoxia response, that is upregulated in senescent cells, and further promoted in MEFs treated with the candidate FAs. Both PARP1 and CDK2 have been reported to directly interact and regulate HIF1A and hypoxia response in different cell types.^54,55^ Conversely, hypoxic stress has also been shown to inhibit cell cycle progression and regulate BCL-xL.^56,57^ While tissue-wide hypoxia promotes cellular senescence and survival, cellular increase in hypoxic response is strongly associated with hypoxic death due to mitochondrial dysfunction, ROS generation, and loss of energy-demanding processes such as protein folding.^56,58,59^ In our transcriptomic and prediction models, these processes are consistently enriched in senescent cells treated with the candidate FAs. Taken together, these suggest a multi-hit potential of SR29384 and SR31133 for a robust induction of apoptotic processes in senescent cells encompassing DNA damage response, pro-death stress signaling, and inhibition of anti-apoptotic mechanisms (Figure 6).

Interestingly, in addition to shared targets for cell cycle regulation and anti-apoptosis, targets exclusive to SR29384 enriched for pathways involving prostaglandin and other pro-inflammatory mediators, ECM disassembly, and activation signaling for many immune cells. All of these processes are routinely associated with release of SASPs and pro-inflammatory signaling, characteristic to senescent cells and their microenvironment.^60,61^ This might present important clues into the remarkable selectivity by SR29384 observed in our senolysis experiments. Conversely, SR31133 exclusive targets enrich strongly for histone deacetylation processes. HDAC inhibition has been shown to induce apoptosis in numerous cancer cells;^62,63^ these findings prompt the need to further explore their role in selectively killing senescent cells as well. SR31133 also shows a significantly stronger binding affinity to Bcl-xL hydrophobic cleft not observed by SR29384 or other flavonoid analogs due to its unique functional moiety. This is supplemented by a network-wide increase in perturbation of Bcl-xL 1^st^ neighbors in SR31133 treated senescent MEFs compared to SR29384. Whether through exclusive mechanisms or stronger inhibition of a key upregulated apoptosis brake in senescent cells, SR31133 is estimated to provide better mechanistic elimination of senescent cells.

It is also important to consider that the targets primarily discussed here are highly filtered to identify specific and selective mechanisms for SR29384 and SR31133. However, due to their flavonoid scaffold, it is likely that they interact with many targets and biological processes that may not dictate their senolytic activity. Understanding these mechanisms could also provide valuable insights for the design, discovery, and optimization of more potent and safer senolytic agents.

In summary, by combining phenotypic drug screening and drug design, we successfully identified two novel flavonoid analogs, SR29384 and SR31133, which demonstrated significantly enhanced senolytic activities compared to fisetin. These analogs exhibited strong potential in targeting a broad spectrum of senescent cells, reducing tissue senescence, and extending healthspan in mice. These novel senolytic may hold great promise for future clinical translation in the treatment of age-related diseases driven by cellular senescence. The structure-activity relationship analysis also provided valuable insights into the key structural features that positively and negatively influence senolytic activity, paving the way for further optimization and development of more potent senolytic compounds.

### Experiments

#### Lead contact

Further information and requests for resources and reagents should be directed to and will be fulfilled by the lead contact, Paul D. Robbins.

#### Reagents

Natural flavonoids were purchased from Cayman Chemical (Michigan, USA). Hoechst 33342 was purchased from ThermoFisher (H1399). C_12_FDG was purchased from Setareh Biotech (7188). Formaldehyde 32% was purchased from Electron Microscopy Sciences (15714). DPPH (2,2-Diphenyl-1-picrylhydrazyl) was purchased from Sigma Aldrich (D9132). L-Ascorbic acid was purchased from Sigma Aldrich (255564).

#### Compound synthesis and characterization

Experimental protocols for compound synthesis and characterization were provided in the Supporting Information.

#### Cells and mice

Primary *Ercc1^-/-^* mouse embryonic fibroblasts (MEFs) and WT MEFs were isolated on embryonic day 12.5-13.5. In brief, mouse embryos were isolated from yolk sac followed by the complete removal of viscera, lung and heart if presented. Embryos were then minced into fine chunks, fed with MEFs medium, cultivated under 3% oxygen to reduce stresses. Cells were split at 1:3 when reaching confluence. MEFs were grown at a 1:1 ratio of Dulbecco’s Modification of Eagles Medium (supplemented with 4.5 g/L glucose and L-glutamine) and Ham’s F10 medium, supplemented with 10% fetal bovine serum, penicillin, streptomycin and non-essential amino acid. To induce oxidative stress-mediated DNA damage, *Ercc1^-/-^* MEFs were switched to 20% oxygen for three passages. WT MEFs were induced senescence by treating hydrogen peroxide H_2_O_2_ (200 μM) or etoposide (2 μM) for 24 h, followed by 5 days in normal culture media.

Human IMR90 lung fibroblasts were obtained from American Type Culture Collection (ATCC) and cultured in EMEM medium with 10% FBS and pen/strep antibiotics. To induce senescence, cells were treated with 20 μM etoposide for 24 h, followed by five days in normal culture media.

Human umbilical vein endothelial cells (HUVECs) were obtained from ATCC and cultured using Endothelial Cell Growth Media plus supplement (without vascular endothelial growth factor (VEGF)) and 1% pen/strep antibiotics. The cells were experimentally treated at late passages 13 to 15.

*Ercc1*^+/−^ and *Ercc1*^+/*Δ*^ mice from C57BL/6J and FVB/n backgrounds were crossed to generate *Ercc1*^−/*Δ*^ mice to prevent potential strain-specific pathology. Aged wild-type C57BL/6J:FVB/NJ mice were generated by crossing C57BL/6J and FVB/n inbred mice purchased from Jackson Laboratory. Mice were left to age for two years before being enrolled into the late life intervention study. Animal protocols used in this study were approved by the University of Minnesota Institutional Animal Care and Use Committees.

### Health evaluation of *Ercc1^-/Δ^* mice

Health assessment of *Ercc1^-/Δ^* mice was conducted twice per week to evaluate age-related symptoms, including body weight, tremor, forelimb grip strength, kyphosis, hindlimb paralysis, gait disorder, dystonia and ataxia. Kyphosis, body condition and coat condition were used to reflect general health conditions. Ataxia, dystonia, gait disorder and tremor were used as indicators of aging-related neurodegeneration. All aging symptoms were scored based on a scale of 0, 0.5 and 1, with the exception of dystonia that has a scale from 0 to 5. The sum of aging scores of each group was used to determine the overall aging conditions, with zero means no symptom presented.

### Senotherapeutic screening

Senescence was evaluated based on SA-β-gal activity using C_12_FDG staining assay. Specifically, senescent *Ercc1^−/−^* MEFs were passaged for three times at 20% O_2_ to induce senescence then seeded at 3000 cells per well in black wall, clear bottom 96 well plates at least 16 hours prior to treatment. Following the addition of drugs, the MEFs were incubated for 48 hours at 20% O_2_. After removing the medium, cells were incubated in 100 nM Bafilomycin A1 in culture medium for 60 min to induce lysosomal alkalinization, followed by incubation with 20 μM fluorogenic substrate C_12_FDG (7188, Setareh Biotech, USA) for 2 h and counterstaining with 2 μg/ml Hoechst 33342 (H1399, Thermo Fisher Scientific, MA, USA) for 15 min. Subsequently, cells were washed with PBS and fixed in 2% paraformaldehyde for 15 min. Finally, cells were imaged with 6 fields per well using a high content fluorescent image acquisition and analysis platform Cytation 1 (BioTek, VT, USA). Cells were seeded at 3000 cells per well in black wall, clear-bottom 96-well plates at least 16 hours prior to treatment. Following the addition of drugs, cells were incubated for 48 hours. After removing the medium, cells were incubated in 100 nM Bafilomycin A1 in culture medium for 1 hour to induce lysosomal alkalinization, followed by incubation with 20 μM fluorogenic substrate C_12_FDG (7188, Setareh Biotech, OR, USA) for 2 hours and counterstaining with 2 μg/mL Hoechst 33342 (H1399, Thermo Fisher Scientific, MA, USA) for 15 minutes. Subsequently, cells were washed with PBS and fixed in 2% paraformaldehyde for 15 minutes. Finally, cells were imaged with six fields per well using the Cytation 1 Cell Imaging Multi-Mode Reader (BioTek Instruments, VT, USA).

### DPPH antiradical assay

The antioxidant activity was evaluated in microplates using the purple free radical 2,2- diphenyl-1-picrylhydrazyl (DPPH•) radical assay. Methanolic solutions fisetin, SR29384, SR31133 at different concentrations (from 0.4 to 100 µM) were tested. Methanolic solutions of L-ascorbic acid at different concentrations were used as standards. The methanolic solution (50 μL) containing the tested compounds at different concentrations was added into 150 μL of methanolic solution of DPPH (100 μM) in 96-well plates. The plates were incubated for 30 min under dark conditions and then measured at 517 nm using a microplate reader. Calculate the percentage inhibition of DPPH radical by the test compounds and the positive control using the formula: %Inhibition = [(Abs_control - Abs_sample)/Abs_control] × 100. The IC_50_ value was calculated on the scavenging activity against DPPH radical.

### RT-qPCR analysis

Total RNA was extracted from cells or snap frozen tissues using Trizol reagent (Thermo Fisher, USA). cDNA was synthesized using High-Capacity cDNA Reverse Transcription Kit (Thermo Fisher, USA). Quantitative PCR reactions were performed with PowerUp™ SYBR™ Green Master Mix (ThermoFisher, USA). The experiments were performed according to the manufacturer’s instructions. The sequences of the primers used were listed below.

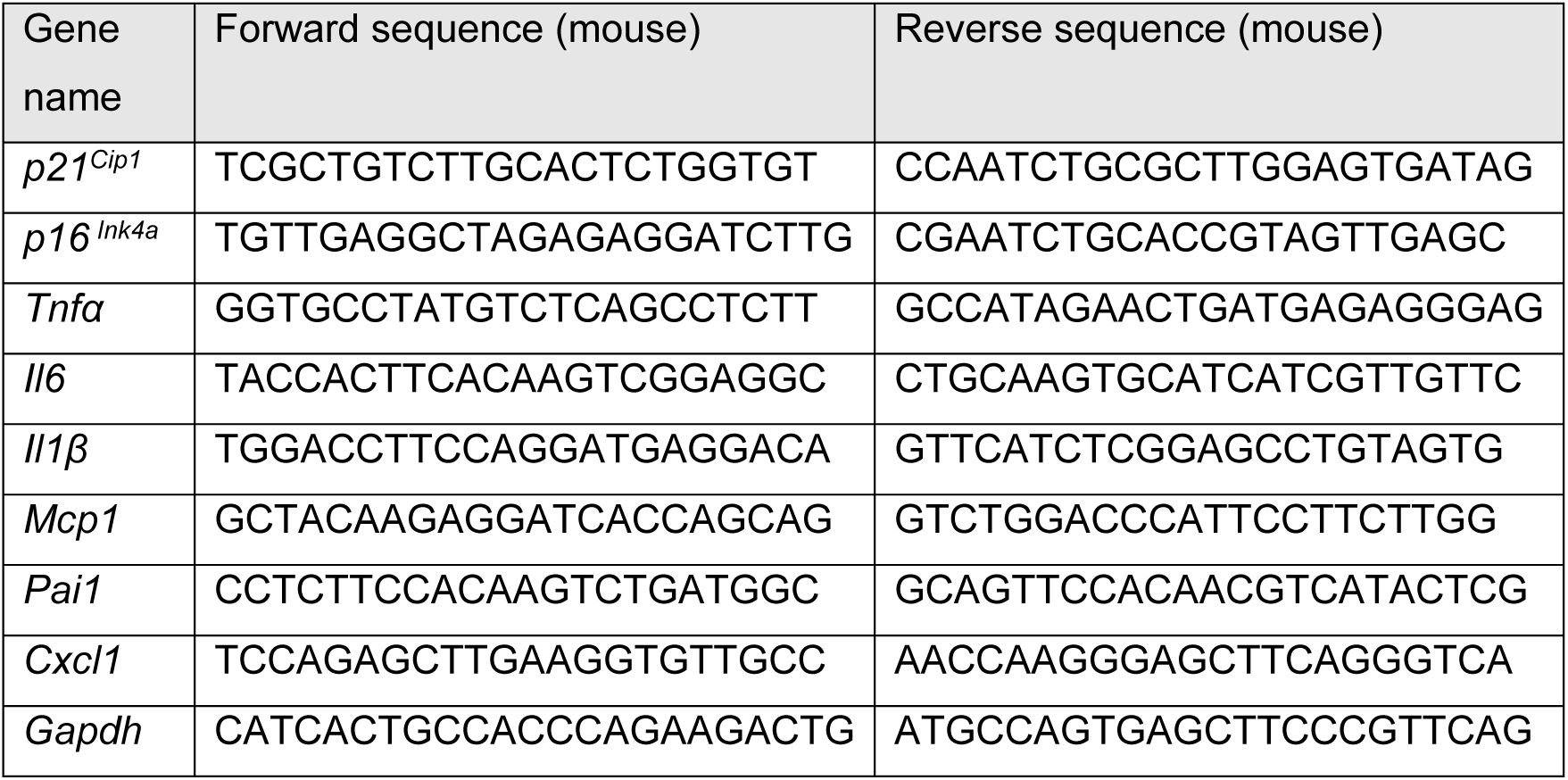

### Pharmacokinetic properties and *in-silico* toxicity screening of compounds

Canonical SMILES of the compounds were subjected to *in-silico* ADME prediction using the SWISSADME web server.^64^ We subsequently performed prediction algorithms for human oral toxicity parameters of the compounds using the OSIRIS Property Explorer v1.1 standalone software (https://www.organic-chemistry.org/prog/peo) and the ProTox-3.0 prediction tool.^65^

### RNA-Seq and enrichment analysis

Non-senescent and senescent MEF cells were treated with fisetin (5 µM), SR29384 (5 µM), or SR31133 (5 µM) for 24 hours. Cells were collected and RNA samples were extracted using Trizol reagent (Thermo Fisher, USA). Samples were quantified using fluorimetry (RiboGreen assay) and RNA integrity was assessed using capillary electrophoresis. All samples had at least 500ng mass and RNA Integrity Number (RIN) of at least 8. Library preparation was carried out using the Illumina TruSeq Stranded Total RNA Library Prep kit, followed by sequencing on the NovaSeq 6000 using 150 PE flow cell, with a sequencing depth of 20 million reads per sample. Quality control on raw sequence data for each sample was performed with FastQC. Read mapping was performed via Hisat2 (v2.1.0) using the mouse genome (GRCm38.94) as reference. Gene quantification was done via Feature Counts for raw read counts. Data has been deposited in the Gene Expression Omnibus under accession number GSE279349. Differentially expressed genes (DEGs) analysis was achieved using the R package edgeR (CPM > 0.05 cutoff). Gene Set Enrichment Analysis (GSEA) was performed using the R Clusterprofiler package.^66,67^ GSEA of the SenMayo panel^31^ was performed using the GSEA 4.3.3 software.^68,69^

### Molecular Pathway Interaction Probability Analysis

Self-normalizing neural-network (DSNNN) based metabolic pathway interaction probability predictions were simulated using the PathwayMap algorithm as previously described.^32^ Canonical SMILES strings for each compound was retrieved from https://pubchem.ncbi.nlm.nih.gov and submitted as inputs (listed in Supplementary Table 1A). Both class-wise and individual pathway results for KEGG and Reactome interaction databases were generated and visualized using using the <u>PlayMolecule</u> interface.

### Structure Based Virtual Target Prediction and Network Construction

The canonical SMILES strings for the ligands were used as inputs for the SwissTargetPrediction algorithm.^33^ Both 2D and 3D similarity measures were used as search parameters with *H. sapiens* selected as the target database. Probability score of > 0.2 was used as statistical cutoff for identified targets and the protein identifiers were retrieved. The set of identifiers for each compound was used as node inputs in the STRING interaction database (https://string-db.org). Results were restricted to *H. sapiens* and submitted list of proteins only. Molecular interaction modes (edges) with highest confidence score (> 0.90) were generated and used to analyze significant (FDR < 0.05) functional enrichment of biological processes (gene ontology). Subsequently, nodes and edges were imported to CytoScape v3.10.2 for visualization using the stringApp 1.7.0 add-on. Node values were assigned using weighted k-shell distribution via wk-shell-decomposition 1.1.0. Subsequently, nodes were clustered using clusterMaker2 2.0 taking undirected edges and wk-shell values as array input with granularity parameter of 2.5. Similarly, for identifying 1^st^ neighbor nodes of targets of interest, all shared DEGs (CPM > 0.5, log_2_FC > |1|, FDR < 0.05) between the SR29384 vs S-Eto and SR31133 vs S-Eto RNA-Seq datasets along with the target proteins were used as node inputs to construct the parent network restricted to *M. musculus*. The parent network was loaded into CytoScape, and 1^st^ neighbor nodes of each individual target were used to construct subnetworks, targets not meeting DEG criteria were excluded. Radial diagrams of the subnetworks were generated using the yFiles Layout Algorithms 1.1.4 add-on.

### Ligand and Receptor Crystal Structure Acquisition and 3D Structure Preparation

The 3D ligand structures of candidate compounds and control were retrieved from https://pubchem.ncbi.nlm.nih.gov. 20 decoy molecules per ligand were generated using the DUD-E database (https://dude.docking.org). Prior to docking runs, we optimized the ligand structures through minimization and polar protonation (pH = 7.4). 3D crystal structures of inhibitor-bound open conformations of the target receptors from the RCSB-PDB database (Supplementary Table 3B).

The structures were optimized using Biovia Discovery Studio Modeling Environment 2024. The crystals were titrated and protonated at pH = 7.4. Co-crystal water, non-cofactor ligand molecules were removed and the appropriate binding cavities were determined by the deep convolutional neural network (DCNN) based protein-binding site prediction algorithm DeepSite (Score ≥ 9.0),^70^ and verified with existing literature. All crystal structures were subjected to the PROCHECK algorithm for stereochemical quality assessment in order to ensure accurate docking pose predictions (https://www.ebi.ac.uk/thornton-srv/software/PROCHECK). Estimations of the whole-model reliability of the 3D structures was performed by evaluating the QMEAN Z-scores of the PDB structures using the ProSA server (https://prosa.services.came.sbg.ac.at/prosa.php). All structures were within acceptable resolution, and Z-scores for NMR and X-ray quality crystal references (Supplementary Table 3B).

### Flexible Molecular Docking Simulation

Structure-based flexible molecular docking of the ligands into the receptor binding cavities were done using the DockThor v2.0 platform.^71^ A receptor search grid of 25Å×25Å×25Å was considered near a pre-defined binding site of each target protein and post-docking OPLS forcefield minimization was carried out. Binding site definitions were set using inhibitors complexed with receptors in crystal structures as templates without restraints. Monomeric subunits of the proteins were used as receptor files and grids were manually defined around the binding cavity for running the docking simulations. DMRTS method was employed for the simulation with 100,000 evaluations per run with an initial population of 750 and 25 runs per ligand. In addition to drug molecules, 20 decoy molecules per complex were used to assess the specificity of the docking protocol and the best binding decoy was used for comparisons. BAK peptide was maintained in original crystal conformation and defined as a ligand molecule to calculate binding affinity. Ligand-receptor interactions were scored using the rDock master scoring function, and binding free energy was calculated as *Binding Free Energy (ΔG) = intermolecular+ ligand intramolecular+ site intramolecular+ external restraints* (https://rxdock.gitlab.io/documentation/devel/html/reference-guide/scoring-functions.html). Post-docking visualization and ligand-receptor interactions were analyzed using Biovia Discovery Studio Modeling Environment 2024 platform.

### Statistical analysis

Data were statistically analyzed by GraphPad Prism software. Two-tailed Student’s *t*-test was performed to determine differences between two groups, and one-way ANOVA with Tukey’s test was used for three groups. A value of *p* < 0.05 was considered as statistically significant, shown as \**p* < 0.05, \*\**p* < 0.01, \*\*\**p* < 0.001 and ****p < 0.0001.

## Funding

Funding for this research was provided by NIH grants R01 AG069819 (PDR), P01 AG043376 (PDR, LJN), U19 AG056278 (PDR, LJN), RO1 AG063543 (LJN), P01 AG062413 (LJN, PDR), U54 AG079754 (LJN, PDR), U54 AG076041 (LJN, PDR), and The Glenn Foundation for Medical Research Postdoctoral Fellowships in Aging Research (LJZ).

---

**Supplementary Figure S1.**
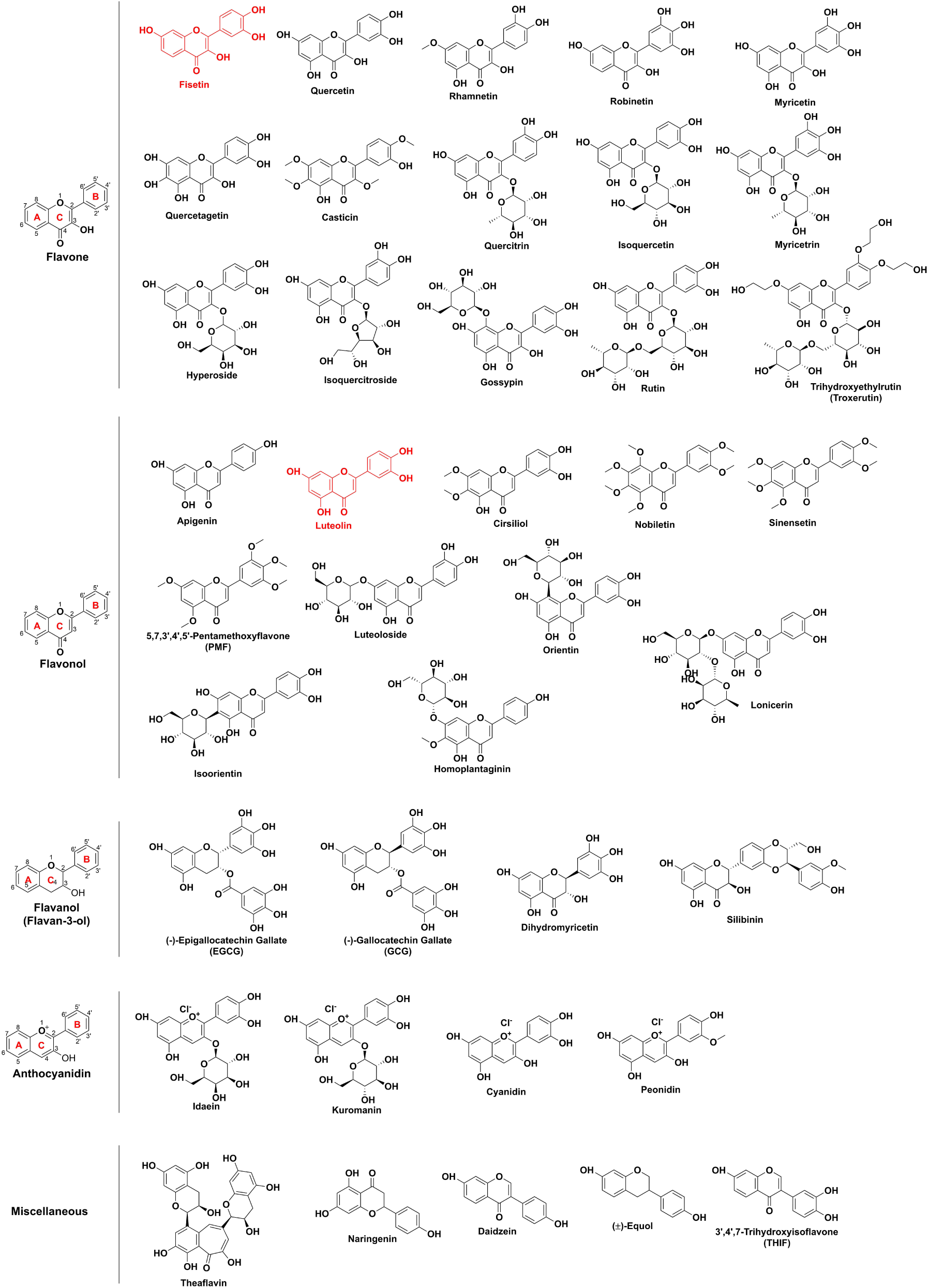
Chemical structures of natural flavonoids tested in the preliminary drug screening.

**Supplementary Figure S2.**
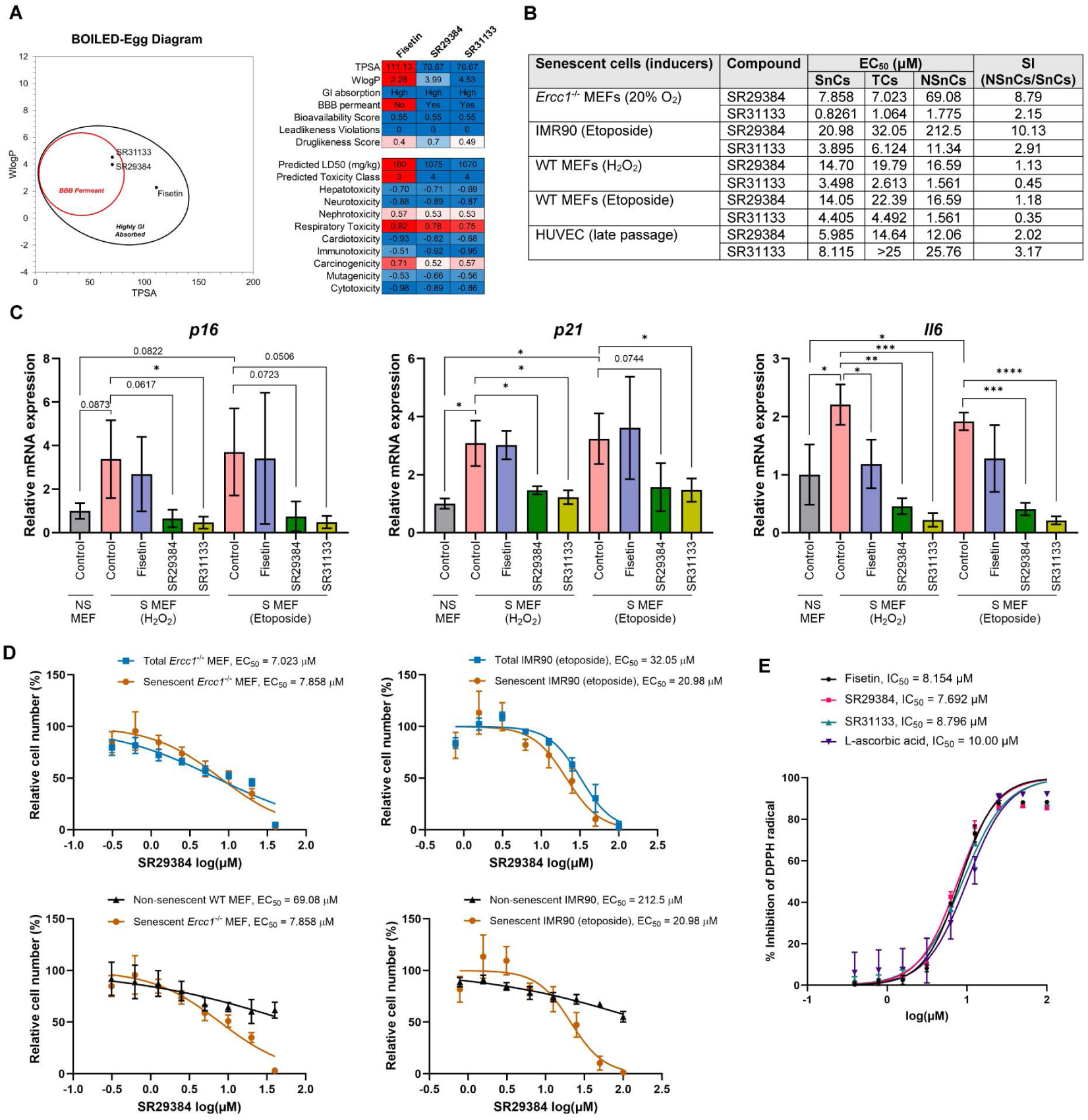
(A) BOILED-Egg diagram and heatmap summarizing the pharmacological and physicochemical properties of fisetin, SR29384, and SR31133, including bioavailability, blood-brain barrier (BBB) permeability, solubility, and predicted toxicity. (B) Table summarizing the senolytic activity (EC_50_) and selectivity index (SI) of SR29384 and SR31133 in various senescent cell models, including *Ercc1*^-/-^ MEFs, IMR90 fibroblasts, WT MEFs, and HUVECs. (C) RT-qPCR analysis showing the relative mRNA expression levels of p16, p21, and IL6 in non-senescent (NS) and senescent (S) MEF cells treated with fisetin (5 µM), SR29384 (5 µM), or SR31133 (5 µM) for 24 hours. Error bars represent SD, n = 3. (D) Selected dose-response curves for the senolytic activity of SR29384. Error bars represent SD, n = 3. (E) Antioxidant activity curves of fisetin, SR29384, and SR31133, measured using the DPPH radical scavenging assay. Error bars represent SD, n = 2.

**Supplementary Figure S3.**
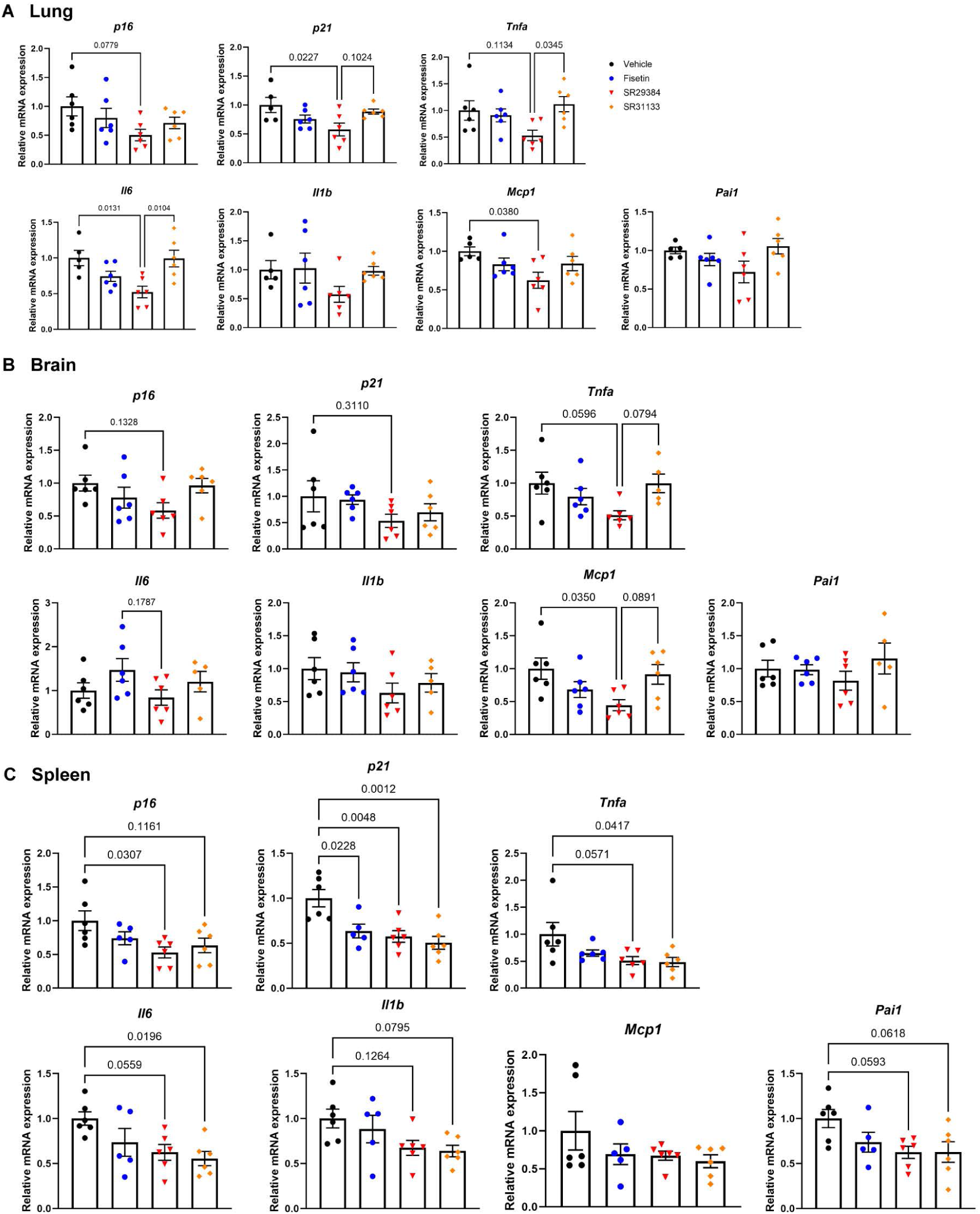
Acute treatment with the flavonoid analogs reduced tissue senescence in naturally aged mice. At 27 months of age, wild-type C57BL/6J mice were administered fisetin, SR29384, or SR31133 for five consecutive days via oral gavage. Two days after the last dose, tissues were collected and analyzed. RT-qPCR analysis of (A) lung, (B) brain, and (C) spleen showing relative mRNA expression levels of senescence markers and SASP factors across treatment groups. Error bars represent SEM for n = 6 mice per group.

**Supplementary Figure S4.**
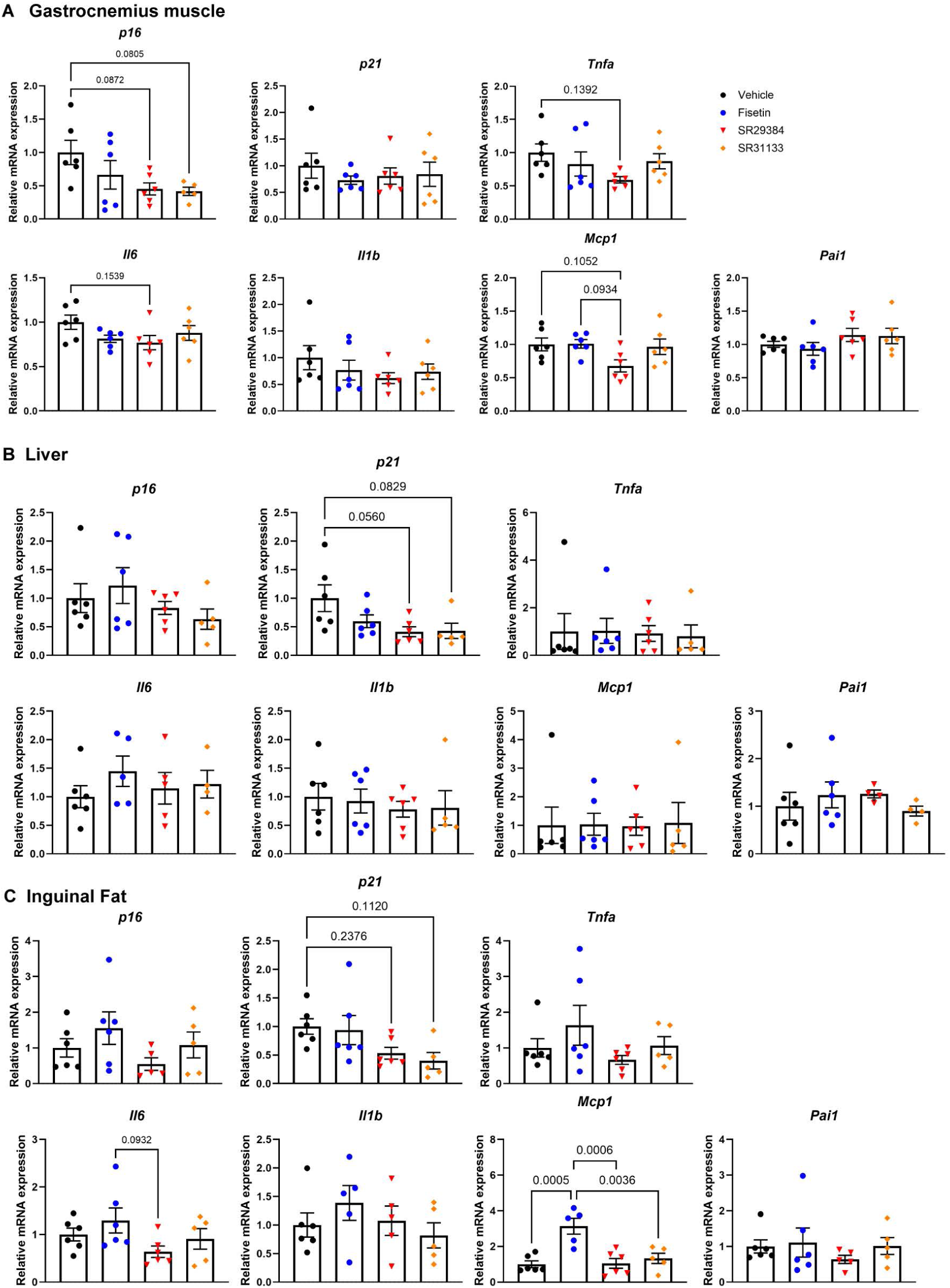
Acute treatment with the flavonoid analogs reduced tissue senescence in naturally aged mice (continued with Figure S3) At 27 months of age, wild-type C57BL/6J mice were administered fisetin, SR29384, or SR31133 for five consecutive days via oral gavage. Two days after the last dose, tissues were collected and analyzed. RT-qPCR analysis of (A) gastrocnemius muscle, (B) liver, and (C) inguinal fat showing relative mRNA expression levels of senescence markers and SASP factors across treatment groups. Error bars represent SEM for n = 6 mice per group.

**Supplementary Figure S5.**
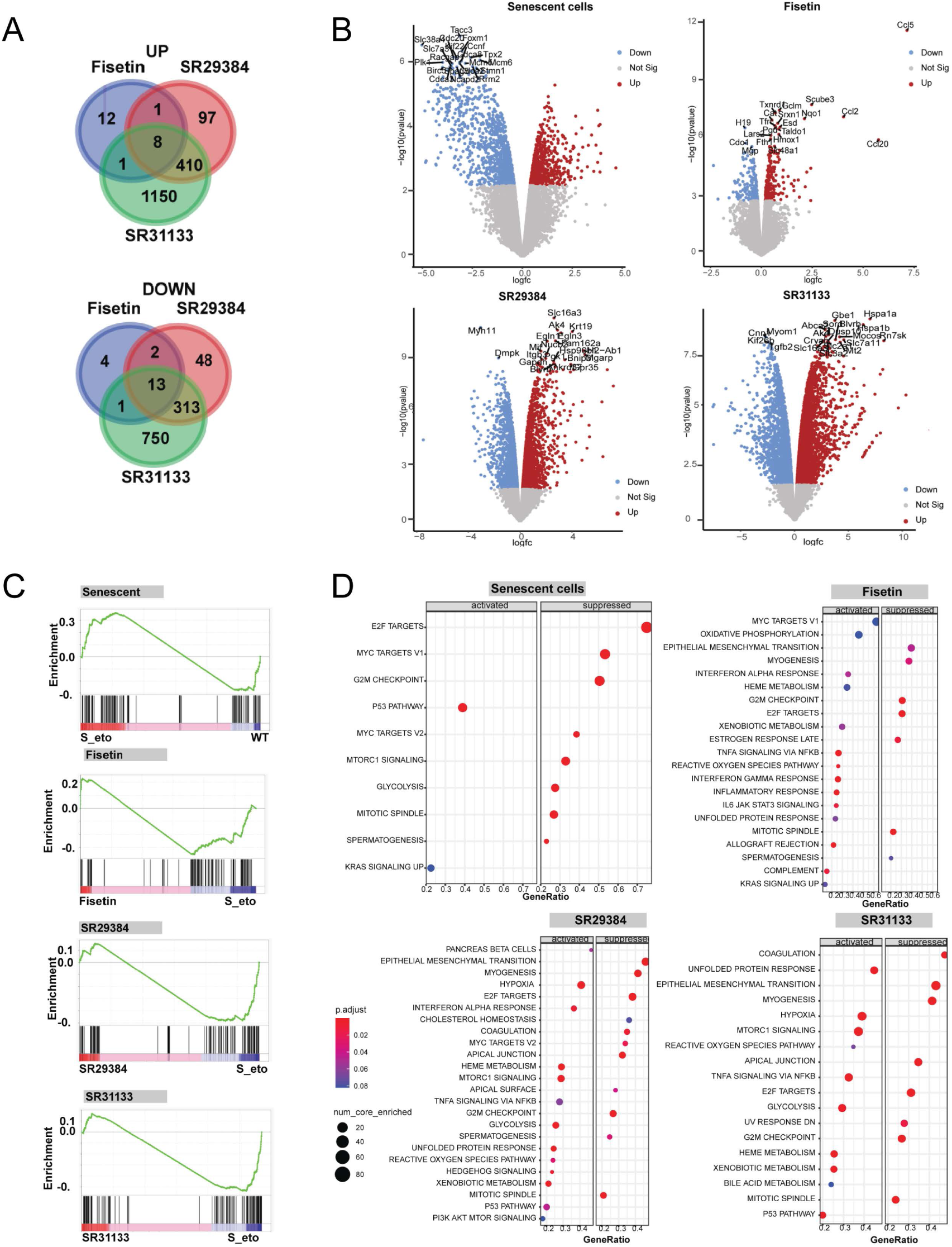
Bulk RNA-Seq analysis of cells treated with Fisetin, SR29384 and SR31133. (A) Venn diagrams showing the number of DEGs upregulated (UP) and downregulated (DOWN) that overlap between the different treatments (CPM > 0.05, log_2_FC > |1| and FDR < 0.05). Three independent biological replicates were analyzed per group. (B) Volcano plot comparing S-Eto (senescent) vs. NS (non-senescent) MEFs, and S-Eto treated with Fisetin, SR29384, or SR31133 vs. untreated S-Eto MEFs. The top 20 most significantly altered genes are labeled. (C) GSEA of the SenMayo panel in S-Eto vs. NS MEFs (NES: 0.84, p val: 0.85) and S-Eto treated with Fisetin (NES: −0.68, p val 0.995), SR29384 (NES: −1.05 FDR, p val 0.381), or SR31133 (NES: −1.27, p val 0.055) vs untreated S-Eto MEFs. Nominal p-values were calculated using a paired two-tailed t-test. (D) Bubble charts of enriched GSEA hallmark terms showing activated and suppressed pathways in S-Eto vs NS MEFs, and S-Eto MEFs treated with Fisetin, SR29384, or SR31133 vs untreated S- Eto MEFs.

**Supplementary Figure S6.**
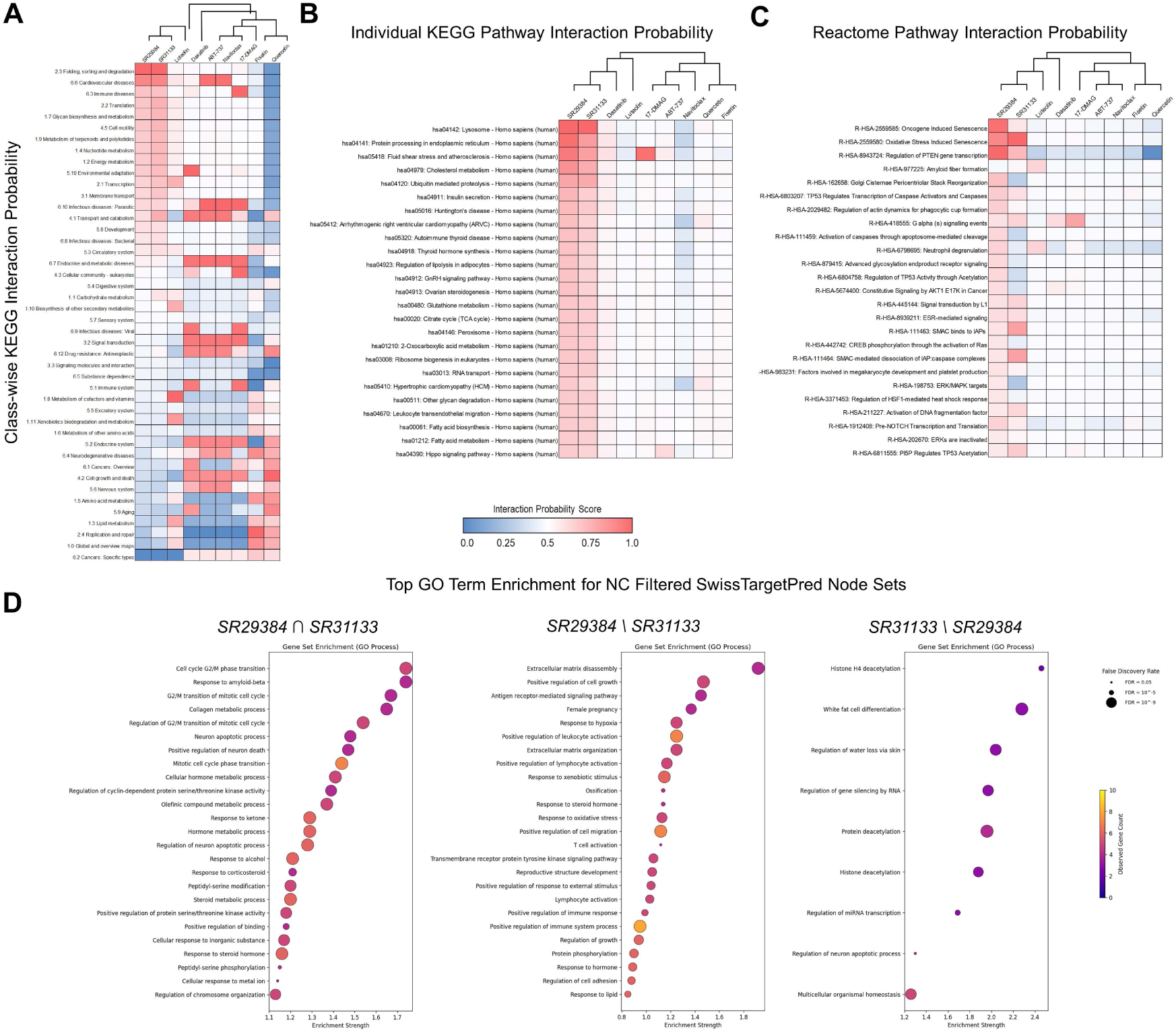
Prediction of top drug-pathway interactions using deep self-normalizing convolutional neural network model and ligand structure similarity. (A) PathwayMap KEGG class-wise pathway interaction probability scores of compound-pathway pairs for candidate FAs and standard senolytic compounds; pathway classes sorted by SR29384 probability scores. (B-C) Top PathwayMap KEGG individual pathway and PathwayMap Reactome interaction probability scores of compound-pathway pairs for candidate FAs and standard senolytic compounds; pathway classes sorted by SR29384 probability scores. (D) Ranked bubble plot of pathway (GO Biological Process) enrichment for constructed interaction network node sets for shared and exclusive SR29384 and SR31133 targets. Bubble color and sizes correspond to number of observed genes in GO term, and FDR values (< 0.05).

**Supplementary Figure S7.**
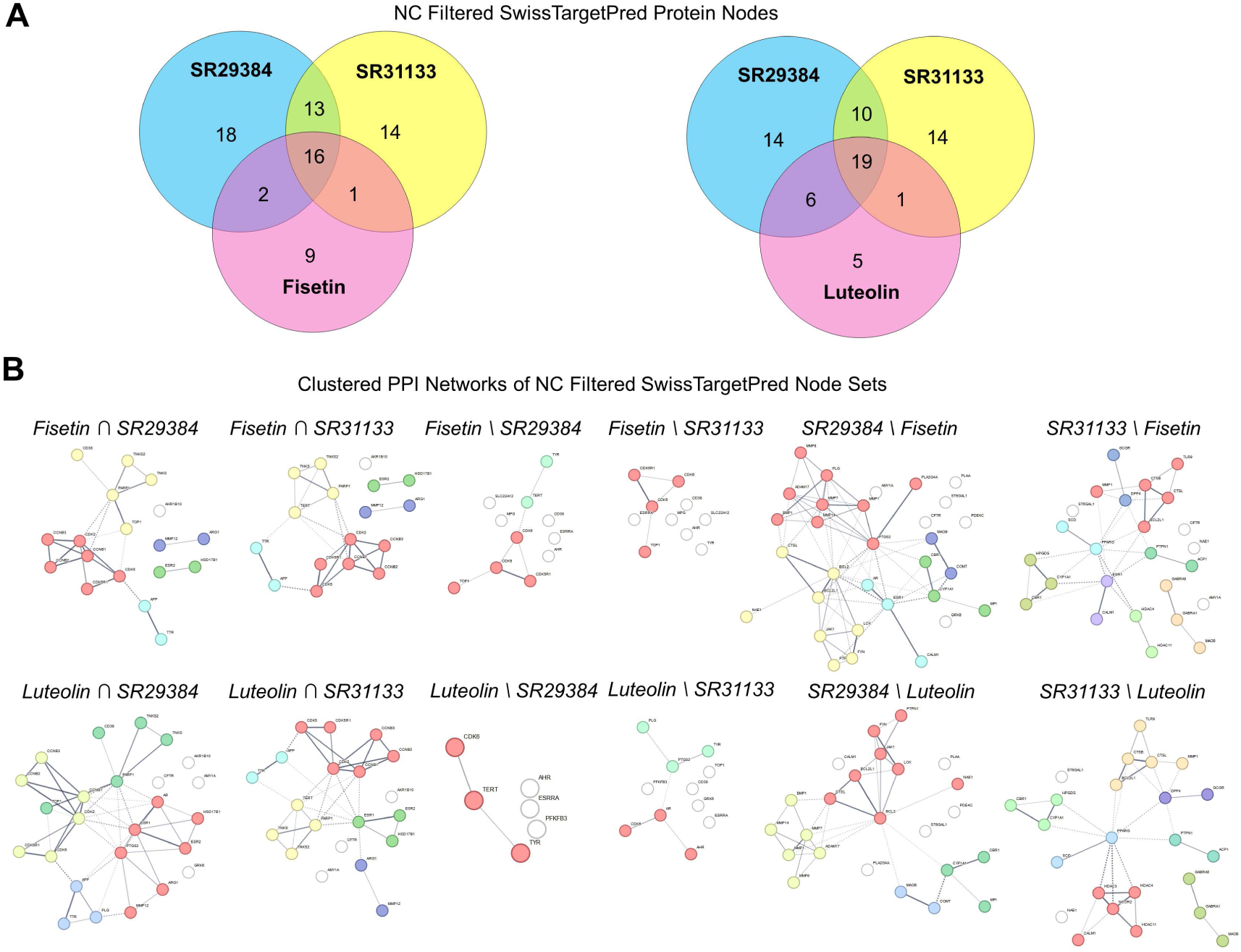
Shared and exclusive targets among FAs predicted based on ligand structure similarity. (A) Venn diagram depicting overlaps among targets for SR29384, SR31133, Fisetin, and Luteolin based on SwissTargetPrediction scores. Target sets were filtered to exclude targets shared with negative control FAs (NC). (B) k-means clustered STRING PPI maps constructed from shared and exclusive NC filtered target proteins for different sub-sets of targets. Nodes are clustered based on degree centrality.

**Supplementary Figure S8.**
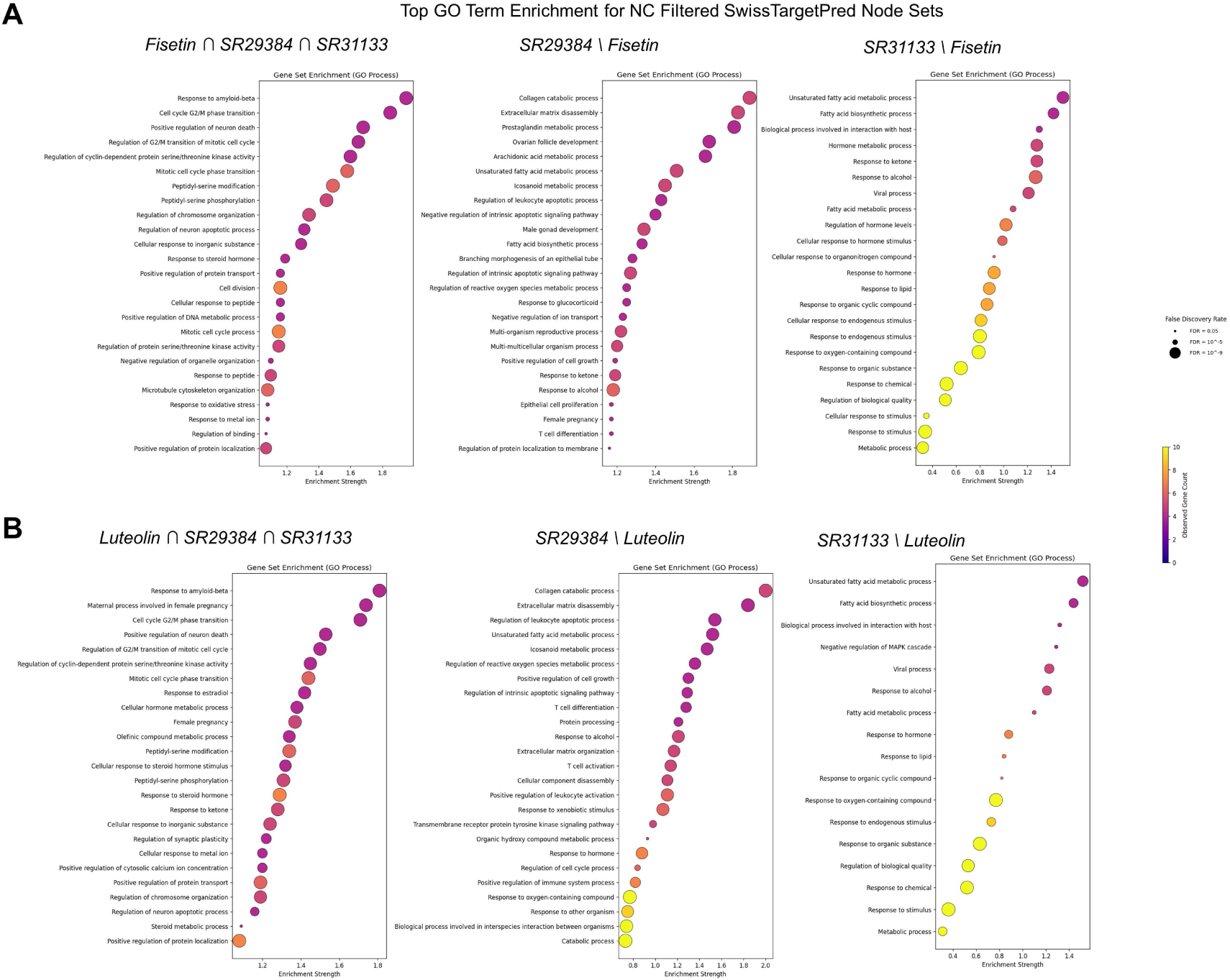
Top drug-pathway interactions based on ligand structure similarity for shared and exclusive targets among FAs. Ranked bubble plot of pathway (GO Biological Process) enrichment for constructed interaction network node sets of shared and exclusive targets for SR29384, SR31133, with (A) Fisetin, and (B) Luteolin. Bubble color and sizes correspond to number of observed genes in GO term, and FDR values (< 0.05).

**Supplementary Figure S9.**
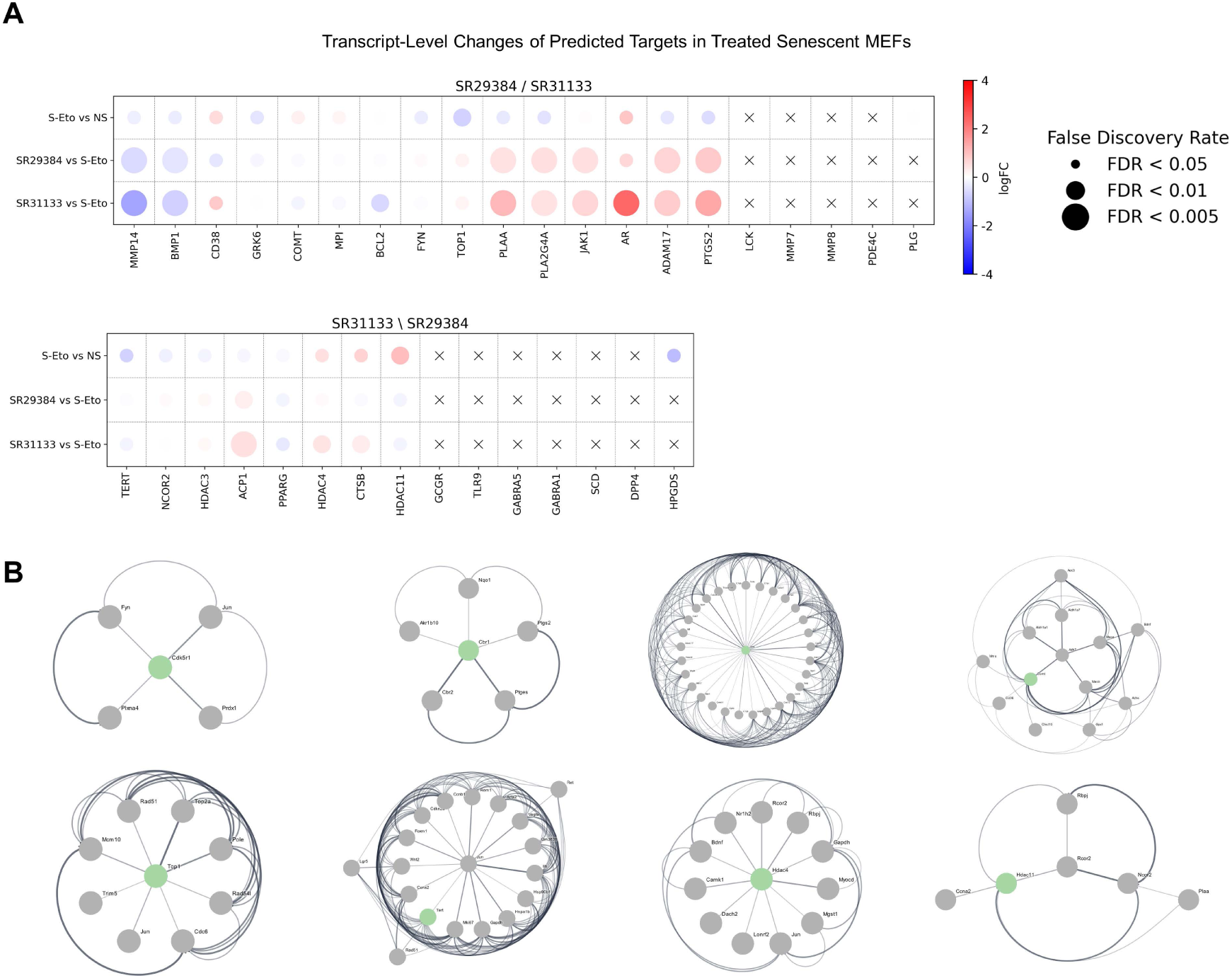
Expression and subnetworks of exclusive molecular targets of top candidate FAs. (A) Bubble plot grid depicting changes in mRNA expression of predicted exclusive targets in etoposide induced senescence in MEFs, and senescent MEFs treated with SR29384, and SR31133. Genes are sorted by log_2_FC values in the SR29384 vs S-Eto dataset. Color and sizes of bubbles correspond to a set scale for log_2_FC and FDR values; “X” symbols represent genes not detected in dataset (CPM ≤ 0.5). (B) Radial diagram of 1^st^ neighbor node DEGs of CDK5R1, CBR1, GRK6, COMT, TOP1, HDAC4 and HDAC11. The target nodes are represented in green; hub genes within the sub-networks are in the center. Connecting edges are defined as level of interaction (probability > 0.9).

## Notes

https://www.ncbi.nlm.nih.gov/geo/query/acc.cgi?acc=GSE279349

### Competing Interest Statement

LJN and PDR are cofounders of Itasca Therapeutics, developing senotherapeutics for aging and age-related diseases. LJZ, LJN, PDR and the University of Minnesota have filed a provisional patent on the application of flavonoid analogs, including SR29384 and SR31133, as a strategy to treat age-related diseases.

## References

1. Lopez-Otin, C., Blasco, M.A., Partridge, L., Serrano, M., and Kroemer, G. (2013). The hallmarks of aging. Cell 153, 1194–1217.

2. Kennedy, B.K., Berger, S.L., Brunet, A., Campisi, J., Cuervo, A.M., Epel, E.S., Franceschi, C., Lithgow, G.J., Morimoto, R.I., Pessin, J.E., et al. (2014). Geroscience: linking aging to chronic disease. Cell 159, 709–713.

3. López-Otín, C., Blasco, M.A., Partridge, L., Serrano, M., and Kroemer, G. (2023). Hallmarks of aging: An expanding universe. Cell 186, 243–278.

4. Hayflick, L., and Moorhead, P.S. (1961). The serial cultivation of human diploid cell strains. Exp Cell Res 25, 585–621.

5. Coppe, J.P., Desprez, P.Y., Krtolica, A., and Campisi, J. (2010). The senescence-associated secretory phenotype: the dark side of tumor suppression. Annu Rev Pathol 5, 99–118.

6. Di Micco, R., Krizhanovsky, V., Baker, D., and d’Adda di Fagagna, F. (2021). Cellular senescence in ageing: from mechanisms to therapeutic opportunities. Nat Rev Mol Cell Biol 22, 75–95.

7. Birch, J., and Gil, J. (2020). Senescence and the SASP: many therapeutic avenues. Genes Dev 34, 1565–1576.

8. Gorgoulis, V., Adams, P.D., Alimonti, A., Bennett, D.C., Bischof, O., Bishop, C., Campisi, J., Collado, M., Evangelou, K., Ferbeyre, G., et al. (2019). Cellular Senescence: Defining a Path Forward. Cell 179, 813–827.

9. Wiley, C.D., and Campisi, J. (2021). The metabolic roots of senescence: mechanisms and opportunities for intervention. Nat Metab 3, 1290–1301.

10. Baker, D.J., Wijshake, T., Tchkonia, T., LeBrasseur, N.K., Childs, B.G., van de Sluis, B., Kirkland, J.L., and van Deursen, J.M. (2011). Clearance of p16Ink4a-positive senescent cells delays ageing-associated disorders. Nature 479, 232–236.

11. Baker, D.J., Childs, B.G., Durik, M., Wijers, M.E., Sieben, C.J., Zhong, J., A. Saltness, R., Jeganathan, K.B., Verzosa, G.C., Pezeshki, A., et al. (2016). Naturally occurring p16Ink4a-positive cells shorten healthy lifespan. Nature 530, 184–189.

12. Wang, B., Wang, L., Gasek, N.S., Zhou, Y., Kim, T., Guo, C., Jellison, E.R., Haynes, L., Yadav, S., Tchkonia, T., et al. (2021). An inducible p21-Cre mouse model to monitor and manipulate p21-highly-expressing senescent cells in vivo. Nature Aging 1, 962–973.

13. Robbins, P.D., Jurk, D., Khosla, S., Kirkland, J.L., LeBrasseur, N.K., Miller, J.D., Passos, J.F., Pignolo, R.J., Tchkonia, T., and Niedernhofer, L.J. (2021). Senolytic Drugs: Reducing Senescent Cell Viability to Extend Health Span. Annu Rev Pharmacol Toxicol 61, 779–803.

14. Prasnikar, E., Borisek, J., and Perdih, A. (2020). Senescent cells as promising targets to tackle age-related diseases. Ageing Res Rev 66, 101251.

15. Kirkland, J.L., and Tchkonia, T. (2020). Senolytic drugs: from discovery to translation. J Intern Med 288, 518–536.

16. Zhang, L., Pitcher, L.E., Prahalad, V., Niedernhofer, L.J., and Robbins, P.D. (2021). Recent advances in the discovery of senolytics. Mech Ageing Dev 200, 111587.

17. Zhang, L., Pitcher, L.E., Prahalad, V., Niedernhofer, L.J., and Robbins, P.D. (2023). Targeting cellular senescence with senotherapeutics: senolytics and senomorphics. FEBS J 290, 1362–1383.

18. Zhang, L., Pitcher, L.E., Yousefzadeh, M.J., Niedernhofer, L.J., Robbins, P.D., and Zhu, Y. (2022). Cellular senescence: a key therapeutic target in aging and diseases. J Clin Invest 132, e158450.

19. Yousefzadeh, M.J., Zhu, Y., McGowan, S.J., Angelini, L., Fuhrmann-Stroissnigg, H., Xu, M., Ling, Y.Y., Melos, K.I., Pirtskhalava, T., Inman, C.L., et al. (2018). Fisetin is a senotherapeutic that extends health and lifespan. EBioMedicine 36, 18–28.

20. Zhu, Y., Doornebal, E.J., Pirtskhalava, T., Giorgadze, N., Wentworth, M., Fuhrmann-Stroissnigg, H., Niedernhofer, L.J., Robbins, P.D., Tchkonia, T., and Kirkland, J.L. (2017). New agents that target senescent cells: the flavone, fisetin, and the BCL-XL inhibitors, A1331852 and A1155463. Aging (Albany NY) 9, 955–963.

21. Jo, J.H., Jo, J.J., Lee, J.M., and Lee, S. (2016). Identification of absolute conversion to geraldol from fisetin and pharmacokinetics in mouse. J Chromatogr B Analyt Technol Biomed Life Sci 1038, 95–100.

22. Seguin, J., Brulle, L., Boyer, R., Lu, Y.M., Ramos Romano, M., Touil, Y.S., Scherman, D., Bessodes, M., Mignet, N., and Chabot, G.G. (2013). Liposomal encapsulation of the natural flavonoid fisetin improves bioavailability and antitumor efficacy. Int J Pharm 444, 146–154.

23. Proshkina, E., Koval, L., Platonova, E., Golubev, D., Ulyasheva, N., Babak, T., Shaposhnikov, M., and Moskalev, A. (2024). Polyphenols as Potential Geroprotectors. Antioxidants & Redox Signaling 40, 564–593.

24. Ravula, A.R., Teegala, S.B., Kalakotla, S., Pasangulapati, J.P., Perumal, V., and Boyina, H.K. (2021). Fisetin, potential flavonoid with multifarious targets for treating neurological disorders: An updated review. Eur J Pharmacol 910, 174492.

25. Rahmani, A.H., Almatroudi, A., Allemailem, K.S., Khan, A.A., and Almatroodi, S.A. (2022). The Potential Role of Fisetin, a Flavonoid in Cancer Prevention and Treatment. Molecules 27, 9009.

26. Sundarraj, K., Raghunath, A., and Perumal, E. (2018). A review on the chemotherapeutic potential of fisetin: In vitro evidences. Biomed Pharmacother 97, 928–940.

27. Gurkar, A.U., and Niedernhofer, L.J. (2015). Comparison of mice with accelerated aging caused by distinct mechanisms. Exp Gerontol 68, 43–50.

28. Yousefzadeh, M.J., Zhao, J., Bukata, C., Wade, E.A., McGowan, S.J., Angelini, L.A., Bank, M.P., Gurkar, A.U., McGuckian, C.A., Calubag, M.F., et al. (2020). Tissue specificity of senescent cell accumulation during physiologic and accelerated aging of mice. Aging Cell 19, e13094.

29. van de Waterbeemd, H., Camenisch, G., Folkers, G., Chretien, J.R., and Raevsky, O.A. (1998). Estimation of blood-brain barrier crossing of drugs using molecular size and shape, and H-bonding descriptors. J Drug Target 6, 151–165.

30. Brand-Williams, W., Cuvelier, M.E., and Berset, C. (1995). Use of a free radical method to evaluate antioxidant activity. LWT - Food Science and Technology 28, 25–30.

31. Saul, D., Kosinsky, R.L., Atkinson, E.J., Doolittle, M.L., Zhang, X., LeBrasseur, N.K., Pignolo, R.J., Robbins, P.D., Niedernhofer, L.J., Ikeno, Y., et al. (2022). A new gene set identifies senescent cells and predicts senescence-associated pathways across tissues. Nat Commun 13, 4827.

32. Jiménez, J., Sabbadin, D., Cuzzolin, A., Martínez-Rosell, G., Gora, J., Manchester, J., Duca, J., and De Fabritiis, G. (2019). PathwayMap: Molecular Pathway Association with Self-Normalizing Neural Networks. J Chem Inf Model 59, 1172–1181.

33. Gfeller, D., Grosdidier, A., Wirth, M., Daina, A., Michielin, O., and Zoete, V. (2014). SwissTargetPrediction: a web server for target prediction of bioactive small molecules. Nucleic Acids Res 42, W32–38.

34. Moffat, J.G., Vincent, F., Lee, J.A., Eder, J., and Prunotto, M. (2017). Opportunities and challenges in phenotypic drug discovery: an industry perspective. Nat Rev Drug Discov 16, 531–543.

35. Berg, E.L. (2021). The future of phenotypic drug discovery. Cell Chem Biol 28, 424–430.

36. Vincent, F., Nueda, A., Lee, J., Schenone, M., Prunotto, M., and Mercola, M. (2022). Phenotypic drug discovery: recent successes, lessons learned and new directions. Nat Rev Drug Discov 21, 899–914.

37. Zhu, R.Z., Li, B.S., Gao, S.S., Seo, J.H., and Choi, B.M. (2021). Luteolin inhibits H(2)O(2)-induced cellular senescence via modulation of SIRT1 and p53. Korean J Physiol Pharmacol 25, 297–305.

38. Xie, T., Yuan, J., Mei, L., Li, P., and Pan, R. (2022). Luteolin suppresses TNF-α- induced inflammatory injury and senescence of nucleus pulposus cells via the Sirt6/NF- κB pathway. Exp Ther Med 24, 469.

39. Zumerle, S., Sarill, M., Saponaro, M., Colucci, M., Contu, L., Lazzarini, E., Sartori, R., Pezzini, C., Rinaldi, A., Scanu, A., et al. (2024). Targeting senescence induced by age or chemotherapy with a polyphenol-rich natural extract improves longevity and healthspan in mice. Nature Aging.

40. Annunziata, C., Castoldi, F., Jan, S., Ang, H.X., Ristovska, M., Melini, S., Welch, R., Riedel, C.G., and Pietrocola, F. (2023). A versatile method for the identification of senolytic compounds. Cell Stress 7, 105–111.

41. Lee, J.Y., Reyes, N.S., Ravishankar, S., Zhou, M., Krasilnikov, M., Ringler, C., Pohan, G., Wilson, C., Ang, K.K.-H., Wolters, P.J., et al. (2024). An in vivo screening platform identifies senolytic compounds that target p16INK4a+ fibroblasts in lung fibrosis. The Journal of Clinical Investigation 134.

42. Rauf, A., Abu-Izneid, T., Imran, M., Hemeg, H.A., Bashir, K., Aljohani, A.S.M., Aljohani, M.S.M., Alhumaydhi, F.A., Khan, I.N., Bin Emran, T., et al. (2023). Therapeutic Potential and Molecular Mechanisms of the Multitargeted Flavonoid Fisetin. Curr Top Med Chem 23, 2075–2096.

43. Kołacz, K., and Robaszkiewicz, A. (2024). PARP1 at the crossroad of cellular senescence and nucleolar processes. Ageing Res Rev 94, 102206.

44. Fleury, H., Malaquin, N., Tu, V., Gilbert, S., Martinez, A., Olivier, M.A., Sauriol, A., Communal, L., Leclerc-Desaulniers, K., Carmona, E., et al. (2019). Exploiting interconnected synthetic lethal interactions between PARP inhibition and cancer cell reversible senescence. Nat Commun 10, 2556.

45. Softah, A., Alotaibi, M.R., Alhoshani, A.R., Saleh, T., Alhazzani, K., Almutairi, M.M., AlRowis, R., Alshehri, S., Albekairy, N.A., Harada, H., et al. (2023). The Combination of Radiation with PARP Inhibition Enhances Senescence and Sensitivity to the Senolytic, Navitoclax, in Triple Negative Breast Tumor Cells. Biomedicines 11.

46. Nehme, J., Mesilmany, L., Farfariello, V., Varela-Eirin, M., Lin, Y., Costa, M.G., Seelen, M., Hillebrands, J.-L., Goor, H.v., Saab, R., et al. (2023). PARP1 inhibition mediates a switch from necrosis to senescence that favors repair from acute oxidative injury. Research Square.

47. Zhu, H., Wei, M., Xu, J., Hua, J., Liang, C., Meng, Q., Zhang, Y., Liu, J., Zhang, B., Yu, X., et al. (2020). PARP inhibitors in pancreatic cancer: molecular mechanisms and clinical applications. Mol Cancer 19, 49.

48. Samuelsson, M.K., Pazirandeh, A., and Okret, S. (2002). A pro-apoptotic effect of the CDK inhibitor p57(Kip2) on staurosporine-induced apoptosis in HeLa cells. Biochem Biophys Res Commun 296, 702–709.

49. Jiang, J., Matranga, C.B., Cai, D., Latham, V.M., Jr., Zhang, X., Lowell, A.M., Martelli, F., and Shapiro, G.I. (2003). Flavopiridol-induced apoptosis during S phase requires E2F-1 and inhibition of cyclin A-dependent kinase activity. Cancer Res 63, 7410–7422.

50. Martin, N., Popgeorgiev, N., Ichim, G., and Bernard, D. (2023). BCL-2 proteins in senescence: beyond a simple target for senolysis? Nat Rev Mol Cell Biol 24, 517–518.

51. Yu, C.Y., Yeung, T.K., Fu, W.K., and Poon, R.Y.C. (2024). BCL-XL regulates the timing of mitotic apoptosis independently of BCL2 and MCL1 compensation. Cell Death Dis 15, 2.

52. Dutta, C., Day, T., Kopp, N., van Bodegom, D., Davids, M.S., Ryan, J., Bird, L., Kommajosyula, N., Weigert, O., Yoda, A., et al. (2012). BCL2 suppresses PARP1 function and nonapoptotic cell death. Cancer Res 72, 4193–4203.

53. Rello-Varona, S., Fuentes-Guirado, M., López-Alemany, R., Contreras-Pérez, A., Mulet-Margalef, N., García-Monclús, S., Tirado, O.M., and García Del Muro, X. (2019). Bcl-x(L) inhibition enhances Dinaciclib-induced cell death in soft-tissue sarcomas. Sci Rep 9, 3816.

54. Zhu, T., Zheng, J.Y., Huang, L.L., Wang, Y.H., Yao, D.F., and Dai, H.B. (2023). Human PARP1 substrates and regulators of its catalytic activity: An updated overview. Front Pharmacol 14, 1137151.

55. Adachi, S., Ito, H., Tamamori-Adachi, M., Ono, Y., Nozato, T., Abe, S., Ikeda, M., Marumo, F., and Hiroe, M. (2001). Cyclin A/cdk2 activation is involved in hypoxia-induced apoptosis in cardiomyocytes. Circ Res 88, 408–414.

56. Chen, N., Chen, X., Huang, R., Zeng, H., Gong, J., Meng, W., Lu, Y., Zhao, F., Wang, L., and Zhou, Q. (2009). BCL-xL is a target gene regulated by hypoxia-inducible factor-1alpha. J Biol Chem 284, 10004–10012.

57. Gardner, L.B., Li, Q., Park, M.S., Flanagan, W.M., Semenza, G.L., and Dang, C.V. (2001). Hypoxia inhibits G1/S transition through regulation of p27 expression. J Biol Chem 276, 7919–7926.

58. Fuhrmann, D.C., and Brüne, B. (2017). Mitochondrial composition and function under the control of hypoxia. Redox Biol 12, 208–215.

59. Yan, J., Sun, C.L., Shin, S., Van Gilst, M., and Crowder, C.M. (2021). Effect of the mitochondrial unfolded protein response on hypoxic death and mitochondrial protein aggregation. Cell Death Dis 12, 711.

60. Marin, I., Serrano, M., and Pietrocola, F. (2023). Recent insights into the crosstalk between senescent cells and CD8 T lymphocytes. NPJ Aging 9, 8.

61. Mavrogonatou, E., Pratsinis, H., Papadopoulou, A., Karamanos, N.K., and Kletsas, D. (2019). Extracellular matrix alterations in senescent cells and their significance in tissue homeostasis. Matrix Biol 75-76, 27–42.

62. Zhang, J., and Zhong, Q. (2014). Histone deacetylase inhibitors and cell death. Cell Mol Life Sci 71, 3885–3901.

63. Bahl, S., Ling, H., Acharige, N.P.N., Santos-Barriopedro, I., Pflum, M.K.H., and Seto, E. (2021). EGFR phosphorylates HDAC1 to regulate its expression and anti-apoptotic function. Cell Death Dis 12, 469.

64. Daina, A., Michielin, O., and Zoete, V. (2017). SwissADME: a free web tool to evaluate pharmacokinetics, drug-likeness and medicinal chemistry friendliness of small molecules. Sci Rep 7, 42717.

65. Banerjee, P., Kemmler, E., Dunkel, M., and Preissner, R. (2024). ProTox 3.0: a webserver for the prediction of toxicity of chemicals. Nucleic Acids Res 52, W513–w520.

66. Wu, T., Hu, E., Xu, S., Chen, M., Guo, P., Dai, Z., Feng, T., Zhou, L., Tang, W., Zhan, L., et al. (2021). clusterProfiler 4.0: A universal enrichment tool for interpreting omics data. Innovation (Camb) 2, 100141.

67. Yu, G., Wang, L.G., Han, Y., and He, Q.Y. (2012). clusterProfiler: an R package for comparing biological themes among gene clusters. Omics 16, 284–287.

68. Subramanian, A., Tamayo, P., Mootha, V.K., Mukherjee, S., Ebert, B.L., Gillette, M.A., Paulovich, A., Pomeroy, S.L., Golub, T.R., Lander, E.S., et al. (2005). Gene set enrichment analysis: a knowledge-based approach for interpreting genome-wide expression profiles. Proc Natl Acad Sci U S A 102, 15545–15550.

69. Mootha, V.K., Lindgren, C.M., Eriksson, K.F., Subramanian, A., Sihag, S., Lehar, J., Puigserver, P., Carlsson, E., Ridderstråle, M., Laurila, E., et al. (2003). PGC-1alpha-responsive genes involved in oxidative phosphorylation are coordinately downregulated in human diabetes. Nat Genet 34, 267–273.

70. Jiménez, J., Doerr, S., Martínez-Rosell, G., Rose, A.S., and De Fabritiis, G. (2017). DeepSite: protein-binding site predictor using 3D-convolutional neural networks. Bioinformatics 33, 3036–3042.

71. Santos, K.B., Guedes, I.A., Karl, A.L.M., and Dardenne, L.E. (2020). Highly Flexible Ligand Docking: Benchmarking of the DockThor Program on the LEADS-PEP Protein-Peptide Data Set. J Chem Inf Model 60, 667–683.

