## Supporting Information for "Development of novel flavonoid senolytics through phenotypic drug screening and drug design"

### Chemistry

All solvents and chemicals were reagent grade. Unless otherwise mentioned, all reagents and solvents were purchased from commercial vendors and used as received. Flash column chromatography was carried out on a Teledyne ISCO CombiFlash Rf system using prepacked columns. Solvents used include hexanes, ethyl acetate (EtOAc), dichloromethane (DCM) and methanol. Purity and characterization of compounds were established by a combination of HPLC, TLC, mass spectrometry, and NMR analyses. $^1$H NMR spectra were recorded on a Bruker Avance DPX-400 (400 MHz) spectrometer and were determined in chloroform-*d* or DMSO-$d^6$ with solvent peaks as the internal reference. Chemical shifts are reported in ppm relative to the reference signal and coupling constant (J) values are reported in hertz (Hz). Multiplicities are given as singlet (s), doublet (d), doublet of doublets (dd), triplet (t), quartet (q), multiplet (m), and broad (br). Thin layer chromatography (TLC) was performed on EMD precoated silica gel 60 F254 plates, and spots were visualized with UV light or iodine staining. Low resolution mass spectra were obtained using a Thermo Scientific ultimate 3000/ LCQ Fleet system (ESI). All compounds containing a stereogenic center are racemic. All test compounds were greater than 95% pure as determined by reverse-phase Shimadzu analytical HPLC using YMC-Pack Pro C18 50cm x 4.6 mm, 5 μm column.

### Chemical Synthesis and Characterization

**Example 1. Preparation of 2-(3,4-dihydroxyphenyl)-6-isopropyl-4H-chromen-4-one (SR29384).**

**Scheme 1. Synthesis of SR29384.**

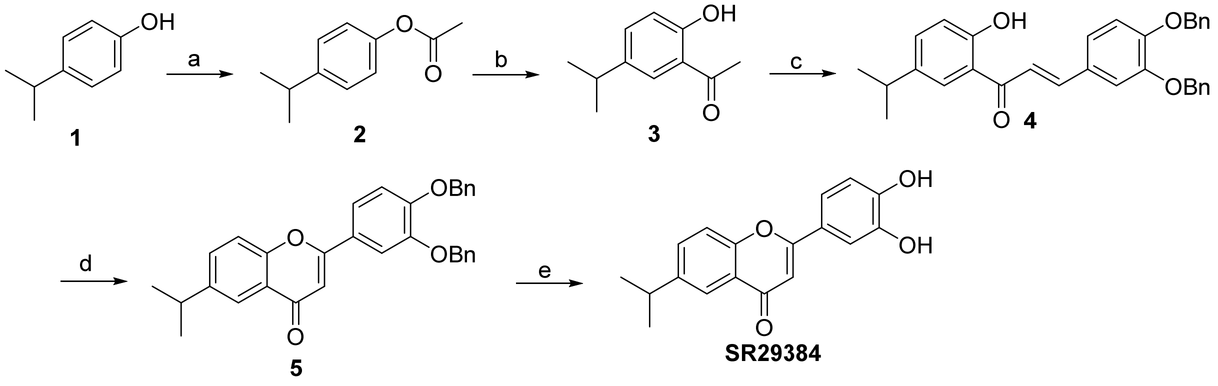


Reagents and conditions: (a) conc. $H_2SO_4$, acetic anhydride, rt; (b) $CF_3SO_3H$, 0 °C; (c) 3,4-

bis(benzyloxy)benzaldehyde/EtOH, NaOH, rt, 2 d; (d) $I_2$, 130 °C; (e) Pd/C, $H_2$, EtOAc/EtOH, overnight.

Step 1: 4-Isopropylphenyl acetate (**2**).

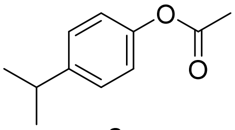

**2**

To the mixture of 4-isopropylphenol (**1**) (5.0 g, 36.91 mmol) in acetic anhydride (5.2 mL, 55.37 mmol) was added 2 drops of concentrated $H_2SO_4$ at room temperature. The completion of the reaction was monitored by analytical HPLC. The reaction mixture was concentrated, redissolved with EtOAc, washed with saturated $NaHCO_3$, brine and dried over $Na_2SO_4$ to obtain the title compound with no further purification. $^1$H NMR (400 MHz, $CDCl_3$): δ 7.22 (d, $J$ = 8.4 Hz, 2H), 6.99 (d, $J$ = 8.4 Hz, 2H), 2.99-2.87 (m, 1H), 2.29 (s, 3H), 1.24 (d, $J$ = 6.9 Hz, 6H).

Step 2: 1-(2-Hydroxy-5-isopropylphenyl)ethan-1-one (**3**).

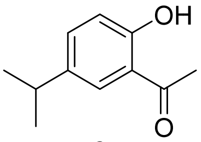

**3**

Intermediate 4-isopropylphenyl acetate **2** (10.17 g, 57.06 mmol) was added slowly to trifluoromethanesulfonic acid (20 mL) at ice bath. The completion of reaction was monitored by analytical HPLC (2 h). The reaction was poured into ice and extracted with EtOAc, washed with water, saturated $NaHCO_3$, and brine, and dried over $Na_2SO_4$. The solvent was removed to obtain the crude which was purified via silica gel flash chromatography to obtain the title compound. $^1$H NMR (400 MHz, $CDCl_3$): δ 12.29 (s, 1H), 7.64 (d, $J$ = 8.2 Hz, 1H), 6.83 (d, $J$ = 1.6 Hz, 1H), 6.77 (dd, $J$ = 8.2 Hz, 1.6 HZ, 1H), 2.99-2.87 (m, 1H), 2.60 (s, 3H), 1.24 (d, $J$ = 6.9 Hz, 6H).

Step 3: (*E*)-3-(3,4-Bis(benzyloxy)phenyl)-1-(2-hydroxy-5-isopropylphenyl)prop-2-en-1-one (**4**).

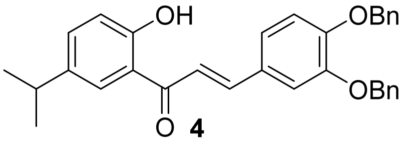

**4**

To a mixture of compound **3** (0.28 g, 1.57 mmol) and 3,4-bis(benzyloxy)benzaldehyde (0.5 g, 1.57 mmol) in EtOH (10 mL) was added NaOH solution (1.6 mL, 5 M in water, 8.0 mmol) and the reaction was stirred at room temperature for 2 days. The mixture was poured into ice water, extracted with EtOAc, washed with water, saturated $NaHCO_3$, and brine, and dried over $Na_2SO_4$. The solvent was removed to obtain the crude which was purified via silica gel flash chromatography to obtain the title compound. $^1$H NMR (400 MHz, $CDCl_3$): δ 7.47-7.26 (m, 14H), 7.14 (d, *J* = 2.0 Hz, 1H), 6.09 (dd, *J* = 8.2 Hz, 2.0 HZ, 1H), 6.92 (d, *J* = 8.4 Hz, 1H), 6.54 (d, *J* = 16.0 Hz, 1H), 5.21 (s, 2H), 5.19 (s, 2H), 2.24-2.16 (m, 1H), 0.96 (d, *J* = 6.9 Hz, 6H).

Step 4: 2-(3,4-Bis(benzyloxy)phenyl)-6-isopropyl-4H-chromen-4-one (**5**).

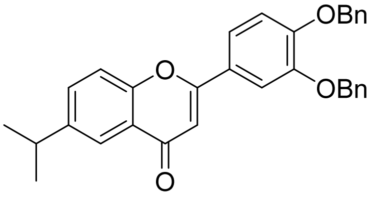


**5**

To a mixture of compound **4** (0.18 g, 0.376 mmol) in DMSO (3 mL) was added iodine (0.01 g, 0.04 mmol), the reaction was heated at 130 °C. The completion of reaction was monitored by analytical HPLC. The reaction was cooled to room temperature and water was added, then extracted with EtOAc, washed with water, saturated $NaHCO_3$, and brine, and dried over $Na_2SO_4$. The solvent was removed to obtain the crude which was purified via silica gel flash chromatography to obtain the title compound. $^1$H NMR (400 MHz, $CDCl_3$): δ 8.05 (d, *J* = 2.2 Hz, 1H), 7.56 (dd, *J* = 8.2 Hz, 2.0 Hz, 1H), 7.49-7.31 (m, 13H), 7.02 (d, *J* = 8.4 Hz, 1H), 6.68 (s, 1H), 5.25 (s, 4H), 3.06-3.00 (m, 1H), 1.30 (d, *J* = 6.9 Hz, 6H).

Step 5: 2-(3,4-Dihydroxyphenyl)-6-isopropyl-4H-chromen-4-one (**SR29384**).

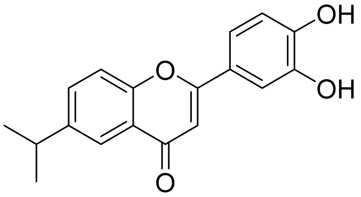


**SR29384**

The mixture of compound **5** (0.05 g) and Pd/C (0.01g, 10% Pd on carbon) in EtOAc/EtOH (1:1, 1 mL) was degassed and stirred under $H_2$ balloon overnight. The completion of reaction was monitored by analytical HPLC. The reaction mixture was filtered through filter paper and the filtrate was concentrated to obtain the crude which was purified by preparative HPLC to obtain the title compound. $^1$H NMR (400 MHz, $CDCl_3$ and $CD_3OD$): δ 7.99 (d, *J* = 2.2 Hz, 1H), 7.61 (dd, *J* = 8.2

Hz, 2.0 Hz, 1H), 7.02 (d, *J* = 8.0 Hz, 1H), 7.45-7.40 (m, 2H), 6.92 (d, *J* = 8.9 Hz, 1H), 6.70 (s, 1H), 3.08-3.00 (m, 1H), 1.30 (d, *J* = 6.9 Hz, 6H).

**Example 2. Preparation of 6-(tert-butyl)-2-(3,4-dihydroxyphenyl)-4H-chromen-4-one (SR32170).**

**Scheme 2. Synthesis of SR32170.**

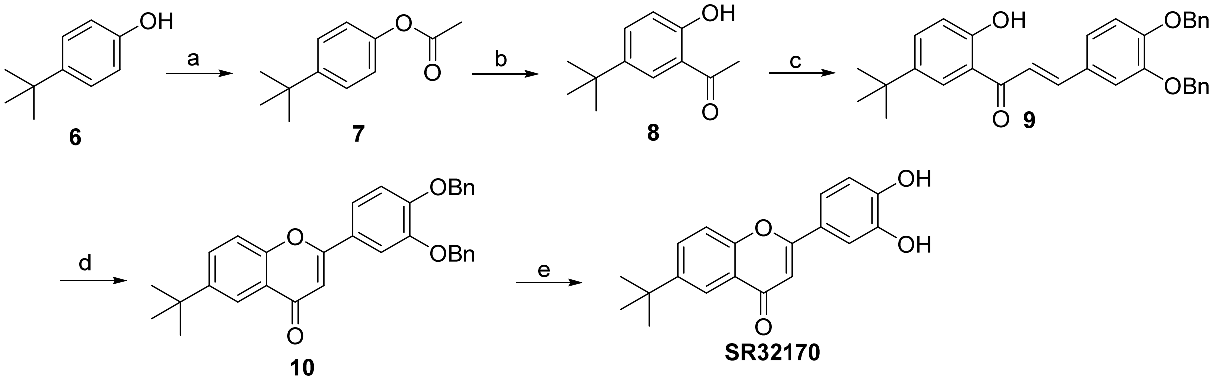

Reagents and conditions: (a) conc. $H_2SO_4$, acetic anhydride, rt; (b) $CF_3SO_3H$, 0 °C; (c) 3,4-bis(benzyloxy)benzaldehyde/EtOH, NaOH, rt, 2 d; (d) $I_2$, 130 °C; (e) Pd/C, $H_2$, EtOAc/EtOH, overnight.

Step 1: 4-(Tert-butyl)phenyl acetate (**7**).

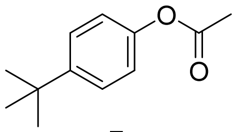

**7**

The title compound was prepared following the same general protocol as described for Step 1, Example 1, using 4-(tert-butyl)phenol **6**. $^1H$ NMR (400 MHz, $CDCl_3$): δ 7.42 (dd, *J* = 6.7 Hz, 2.0Hz, 2H), 7.04 (dd, *J* = 6.7 Hz, 2.0Hz, 2H), 2.31 (s, 3H), 1.34 (s, 9H).

Step 2: 1-(5-(Tert-butyl)-2-hydroxyphenyl)ethenone (**8**).

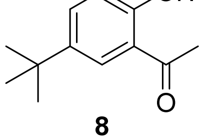

**8**

The title compound was prepared following the same general protocol as described for Step 2, Example 1, using compound **7**. $^{1}$H NMR (400 MHz, $CDCl_3$): δ 12.05 (s, 1H), 7.61 (dd, *J* = 8.7Hz, 2.4 Hz, 1H), 6.86 (d, *J* = 8.7 Hz, 1H), 2.57 (s, 3H), 1.25 (s, 9H).

Step 3: (*E*)-3-(3,4-Bis(benzyloxy)phenyl)-1-(5-(tert-butyl)-2-hydroxyphenyl)prop-2-enone (**9**).

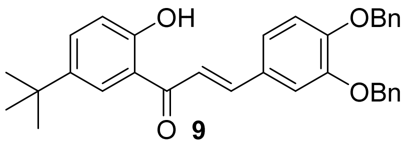

**9**

The title compound was prepared following the same general protocol as described for Step 3, Example 1, using compound **8**. $^{1}$H NMR (400 MHz, $CDCl_3$): δ 12.35 (s, 1H), 7.96 (d, *J* = 2.6Hz, 1H), 7.80 (d, *J* = 8.8Hz, 1H), 7.63-7.59 (m, 2H), 7.49-7.31 (m, 12H), 7.16 (d, 8.4Hz, 1H), 8.95 (d, 8.4Hz, 1H), 5.26 (s, 2H), 5.23 (s, 2H), 1.35 (s, 9H).

Step 4: 2-(3,4-Bis(benzyloxy)phenyl)-6-(tert-butyl)-4H-chromen-4-one (**10**).

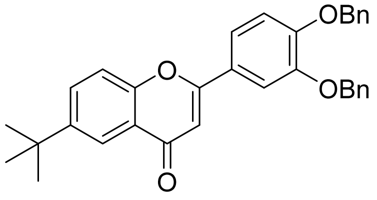

**10**

The title compound was prepared following the same general protocol as described for Step 4, Example 1, using compound **9**. $^{1}$H NMR (400 MHz, $CDCl_3$): δ 8.05 (d, *J* = 2.2 Hz, 1H), 7.64 (dd, *J* = 8.2 Hz, 2.0 Hz, 1H), 7.39-7.28 (m, 13H), 7.26 (d, *J* = 8.4 Hz, 1H), 6.95 (s, 1H), 5.16 (s, 4H), 1.30 (s, 9H).

Step 5: 6-(Tert-butyl)-2-(3,4-dihydroxyphenyl)-4H-chromen-4-one (**SR32170**).

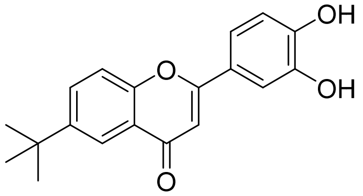

**SR32170**

The title compound was prepared following the same general protocol as described for Step 5, Example 1, using compound **10**. $^{1}$H NMR (400 MHz, DMSO-$d_6$): δ 7.97 (d, *J* = 2.4 Hz, 1H), 7.91 (dd, *J* = 8.2 Hz, 2.4 Hz, 1H), 7.67 (m, 2H), 6.91 (d, *J* = 8.9 Hz, 1H), 6.75 (s, 1H), 1.35 (s, 9H).

**Example 3. Preparation of 6-cyclopentyl-2-(3,4-dihydroxyphenyl)-4H-chromen-4-one (SR31133).**

**Scheme 3. Synthesis of SR31133.**

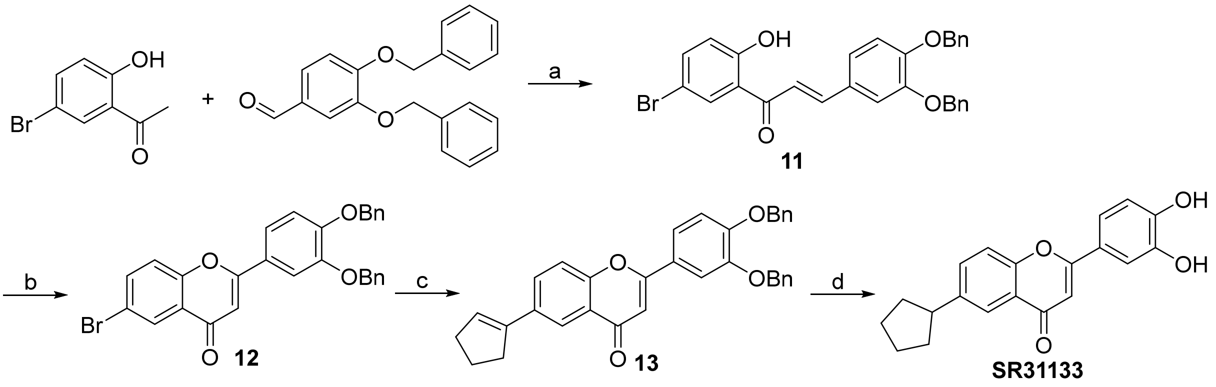


Reagents and conditions: (a) NaOMe/MeOH, 90 °C, 16 h; (b) $I_2$, 130 °C; (c) 2-(cyclopent-1-en-1-yl)-4,4,5,5-tetramethyl-1,3,2-dioxaborolane, $Pd(PPh_3)_4$, $K_2CO_3$, dioxane/$H_2O$, 140 °C, overnight; (d) Pd/C, $H_2$, EtOAc/EtOH, overnight.

Step1: (*E*)-3-(3,4-Bis(benzyloxy)phenyl)-1-(5-bromo-2-hydroxyphenyl)prop-2-en-1-one (**11**).

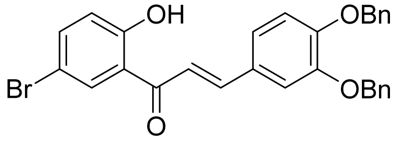


**11**

To a mixture of 1-(5-bromo-2-hydroxyphenyl)ethan-1-one (7.0 g, 32.55 mmol) and 3,4-bis(benzyloxy)benzaldehyde (10.36 g, 32.55 mmol) in MeOH (80 mL) was added freshly made NaOMe in MeOH (120 mL, 1.4 M, 168.0 mmol) and the reaction was stirred at 90 °C oil bath for 16 h. The mixture was cooled to room temperature and poured into ice water, and the resulting mixture was adjusted to pH 3-4 by addition of 5 N HCl. Filter the mixture to obtain the solid as the desired product with no further purification. ESI-MS (m/z): 514.8, 516.7 $[M+1]^+$.

Step 2: 2-(3,4-Bis(benzyloxy)phenyl)-6-bromo-4H-chromen-4-one (**12**).

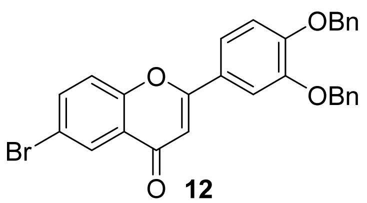

The title compound was prepared following the same general protocol as described for Step 4, Example 1, using compound **11**. ESI-MS (m/z): 512.8, 514.7 $[M+1]^+$. $^1$H NMR (400 MHz, DMSO-$d_6$): δ 8.14 (d, *J* = 4.0 Hz, 1H), 8.04 (dd, *J* = 8.0 Hz, 4.0 Hz, 1H), 7.84-7.82 (m, 2H), 7.76 (dd, *J* = 8.0 Hz, 4.0 Hz, 1H), 7.56-7.50 (m, 4H), 7.47-7.42 (m, 4H), 9.39-7.35 (m, 2H), 7.29 (d, *J* = 12.0 Hz, 1H), 7.13 (s, 1H), 5.33 (s, 2H), 5.30 (s, 2H).

Step 3: 2-(3,4-Bis(benzyloxy)phenyl)-6-(cyclopent-1-en-1-yl)-4H-chromen-4-one (**13**).

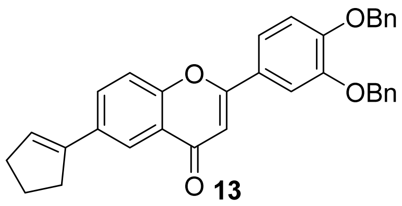

A mixture of compound **12** (0.15 g, 0.292 mmol), 2-(cyclopent-1-en-1-yl)-4,4,5,5-tetramethyl-1,3,2-dioxaborolane (0.085 g, 0.438 mmol), $Pd(PPh_3)_4$ (0.017 g, 0.015 mmol) and $K_2CO_3$ (0.061 g, 0.441 mmol) in dioxane (1 mL) and water (0.2 mL) was stirred at 140 °C overnight under nitrogen. The mixture was added water and EtOAc, the mixture was stirred and separated, the aqueous phase was extracted with EtOAc twice, the combined organic phases were washed with brine and dried with $Na_2SO_4$, filtered through silica gel to get oil, which was purified by silica gel column to obtain the title compound. ESI-MS (m/z): 501.0 $[M+1]^+$.

Step 4: 6-Cyclopentyl-2-(3,4-dihydroxyphenyl)-4H-chromen-4-one (**SR31133**).

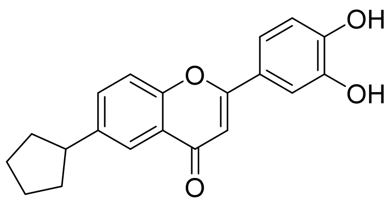

**SR31133**

The title compound was prepared following the same general protocol as described for Step 5, Example 1, using compound **13**. ESI-MS (m/z): 322.6 $[M+1]^+$. $^1$H NMR (400 MHz, DMSO-$d_6$): δ 9.90 (s, 1H), 9.43 (s, 1H), 7.89 (d, *J* = 4.0 Hz, 1H), 7.76 (dd, *J* = 8.0 Hz, 4.0 Hz, 1H), 7.68 (d, *J* =

8.0 Hz, 1H), 7.52-7.46 (m, 2H),6.95 (d, *J* = 16.0 Hz, 1H), 6.77 (s, 1H), 3.20-3.12 (m, 1H)m, 2.15-2.08 (m, 2H), 1.88-1.79 (m, 2H), 1.77-1.69 (m, 2H), 1.66-1.55 (m, 2H).

**Example 4. Preparation of 6-cyclohexyl-2-(3,4-dihydroxyphenyl)-4H-chromen-4-one (SR31872)**

Step 1: 6-Cyclohexyl-2-(3,4-dihydroxyphenyl)-4H-chromen-4-one

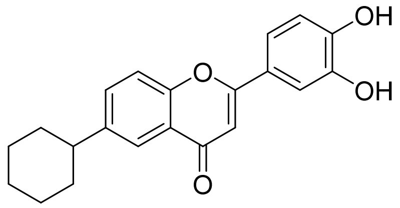


**SR31872**

The title compound was prepared following the same general protocol as described for Step 5, Example 1, using 6-(cyclohex-1-en-1-yl)-2-(3,4-dihydroxyphenyl)-4H-chromen-4-one. $^1$H NMR (400 MHz, DMSO-$d_6$): δ 7.84 (d, *J* = 2.1 Hz, 1H), 7.71-7.63 (m, 2H), 7.45 -7.43 (m, 2H), 6.91 (d, *J* = 8.20, 1H), 6.74 (s, 1H), 2.67-2.64 (m, 1H), 1.86-1.51 (m 5H), 1.49-1.24 (m, 5H).

**Example 5. Preparation of 2-(3,4-bis(benzyloxy)phenyl)-6-(cyclohex-1-en-1-yl)-4H-chromen-4-one (SR31871).**

Step1: 2-(3,4-Bis(benzyloxy)phenyl)-6-(cyclohex-1-en-1-yl)-4H-chromen-4-one.

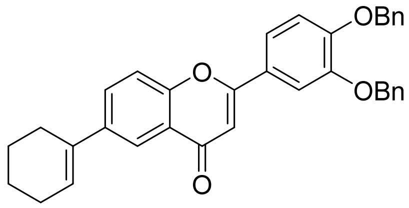


The title compound was prepared following the same general protocol as described for Step 3, Example 3, using 2-(1-cyclohexenyl)-4,4,5,5-tetramethyl-1,3,2-dioxaborolane. $^1$H NMR (400 MHz, $CDCl_3$): δ 8.08 (d, *J* = 2.3 Hz, 1H), 7.37 (dd, *J* = 5.2 Hz, 2.4 Hz, 2H), 7.33 -7.17 (m, 12H), 6.94 (d, *J* = 8.28, 1H), 6.58 (s, 1H), 6.17 (q, *J* = 2.32 Hz, 1H), 5.16 (s, 4H), 2.40-2.37 (m, 2H), 2.17-2.13 (m, 2H), 1.75-1.72 (m, 2H), 1.69-1.62 (m, 2H).

Step 2: 6-(Cyclohex-1-en-1-yl)-2-(3,4-dihydroxyphenyl)-4H-chromen-4-one (**SR31871**).

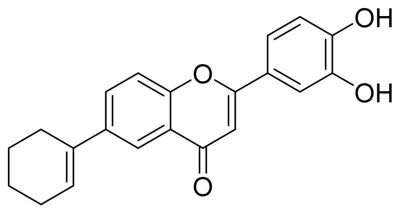

**SR31871**

To the mixture of 2-(3,4-bis(benzyloxy)phenyl)-6-(cyclohex-1-en-1-yl)-4H-chromen-4-one (0.11 g, 0.22 mmol) in DCM (3 mL) at room temperature was added $BBr_3$ (0.87 mL, 1M in DCM, 0.87 mmol) dropwise. The completion of the reaction was monitored by analytical HPLC. The mixture was slowly poured into cold saturated $NaHCO_3$, extracted with DCM, washed with brine and dried over $Na_2SO_4$. The solvent was removed to obtain the crude which was purified by analytical HPLC to obtain the title compound. $^1$H NMR (400 MHz, DMSO-$d_6$): δ 9.88 (br, 1H), 9.47 (br, 1H), 7.95 (s, 1H), 6.94 (dd, $J$ = 8.20 Hz, 2.10 Hz, 1H), 7.68 (d, $J$ = 8.20, 1H), 7.46 (m, 2H), 6.92 (d, $J$ = 8.2Hz, 2.1 Hz, 1H), 6.75 (s, 1H), 6.33 (t, $J$ = 4 Hz, 1H), 2.44-2.43 (m, 2H), 2.24-2.22 (m, 2H), 1.80-1.74 (m, 2H), 1.66-1.60 (m, 2H).

**Example 6. Preparation of 2-(3,4-dihydroxyphenyl)-6-phenyl-4H-chromen-4-one (SR31131).**

Step1: 2-(3,4-Bis(benzyloxy)phenyl)-6-phenyl-4H-chromen-4-one.

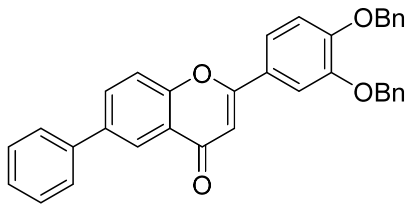

The title compound was prepared following the same general protocol as described for Step 3, Example 3, using phenylboronic acid. ESI-MS (m/z): 511.0 $[M+1]^+$.

Step 2: 2-(3,4-dihydroxyphenyl)-6-phenyl-4H-chromen-4-one (**SR31131**).

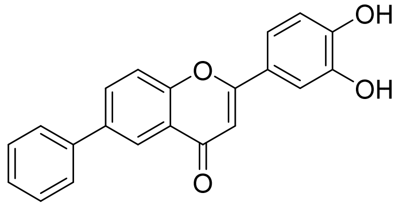

**SR31131**

The title compound was prepared following the same general protocol as described for Step 2, Example 5, using 2-(3,4-bis(benzyloxy)phenyl)-6-phenyl-4H-chromen-4-one. ESI-MS (m/z):

330.5 [M+1]$^+$. $^1$H NMR (400 MHz, DMSO-$d_6$): δ 9.95 (s, 1H), 9.45 (s, 1H), 8.27 (d, *J* = 4.0 Hz, 1H), 8.16 (dd, *J* = 8.0 Hz, 4.0 Hz, 1H), 7.87 (d, *J* = 8.0 Hz, 1H), 7.82-7.79 (m, 2H), 7.58-7.44 (m, 5H), 6.96 (d, *J* = 8.0 Hz, 1H), 6.77 (s, 1H).

**Example 7. Preparation of 2-(3,4-dihydroxyphenyl)-6-(2-hydroxyphenyl)-4H-chromen-4-one (SR31127).**

Step1: 2-(3,4-Bis(benzyloxy)phenyl)-6-(2-hydroxyphenyl)-4H-chromen-4-one.

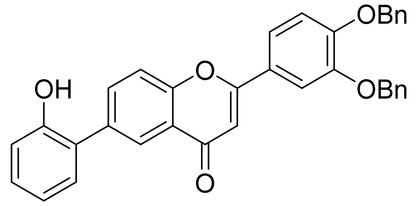


The title compound was prepared following the same general protocol as described for Step 3, Example 3, using (2-hydroxyphenyl)boronic acid. ESI-MS (m/z): 527.0 [M+1]$^+$.

Step 2: 2-(3,4-Dihydroxyphenyl)-6-(2-hydroxyphenyl)-4H-chromen-4-one (**SR31127**).

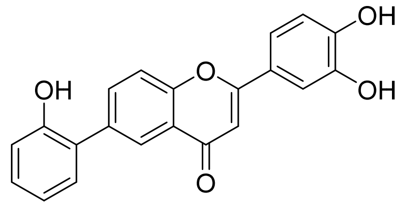


**SR31127**

The title compound was prepared following the same general protocol as described for Step 2, Example 5, using 2-(3,4-bis(benzyloxy)phenyl)-6-(2-hydroxyphenyl)-4H-chromen-4-one. ESI-MS (m/z): 346.6 [M+1]$^+$. $^1$H NMR (400 MHz, DMSO-$d_6$): δ 9.77 (s, 2H), 8.22 (d, *J* = 4.0 Hz, 1H), 8.03 (dd, *J* = 8.0 Hz, 4.0 Hz, 1H), 7.78 (d, *J* = 8.0 Hz, 1H), 7.52-7.50 (m, 2H), 7.49 (dd, *J* = 8.0 Hz, 4.0 Hz, 1H), 7.26 (m, 1H), 7.03 (d, *J* = 8.0 Hz, 1H), 7.04-6.95 (m, 2H), 6.82 (s, 1H).

**Example 8. Preparation of 2-(3,4-dihydroxyphenyl)-6-(3-hydroxyphenyl)-4H-chromen-4-one (SR31130).**

Step1: 2-(3,4-Bis(benzyloxy)phenyl)-6-(3-hydroxyphenyl)-4H-chromen-4-one.

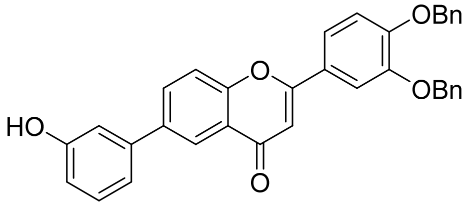

The title compound was prepared following the same general protocol as described for Step 3, Example 3, using (3-hydroxyphenyl)boronic acid. ESI-MS (m/z): 527.0 $[M+1]^+$.

Step 2: 2-(3,4-Dihydroxyphenyl)-6-(3-hydroxyphenyl)-4H-chromen-4-one (**SR31130**).

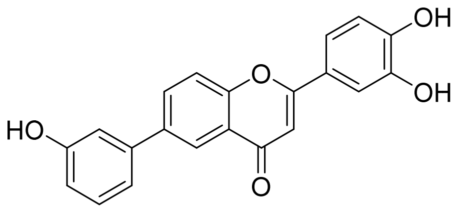


**SR31130**

The title compound was prepared following the same general protocol as described for Step 2, Example 5, using 2-(3,4-bis(benzyloxy)phenyl)-6-(3-hydroxyphenyl)-4H-chromen-4-one. ESI-MS (m/z): 346.6 $[M+1]^+$. $^1$H NMR (400 MHz, DMSO-$d_6$): δ 9.94-9.64 (m, 3H), 8.20 (d, *J* = 4.0 Hz, 1H), 8.09 (dd, *J* = 8.0 Hz, 4.0 Hz, 1H), 7.84 (d, *J* = 8.0 Hz, 1H), 7.53-7.50 (m, 2H), 7.37-7.32 (m, 1H), 7.20 (d, *J* = 8.0 Hz, 1H), 7.15-7.12 (m, 1H),6.97-6.94 (m, 1H), 6.87-6.83 (m, 2H).

**Example 9. Preparation of 2-(3,4-dihydroxyphenyl)-6-(4-methoxyphenyl)-4H-chromen-4-one (SR31870).**

Step 1: 2-(3,4-Bis(benzyloxy)phenyl)-6-(4-methoxyphenyl)-4H-chromen-4-one.

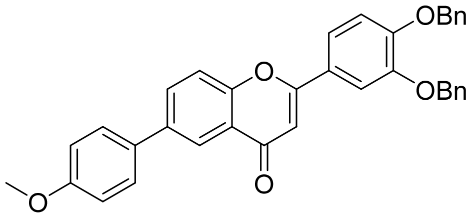


The title compound was prepared following the same general protocol as described for Step 3, Example 3, using (4-methoxyphenyl)boronic acid. $^1$H NMR (400 MHz, $CDCl_3$): δ 8.48 (d, *J* = 2.3 Hz, 1H), 7.70 (dd, *J* = 8.2 Hz, 2.1 Hz, 1H), 7.59 -7.28 (m, 17H), 7.07-7.01 (m, 1H), 6.73 (s, 1H), 5.32 (s, 2H), 5.29 (s, 2H), 3.89 (s, 3H).

Step 2: 2-(3,4-Dihydroxyphenyl)-6-(4-methoxyphenyl)-4H-chromen-4-one (**SR31870**).

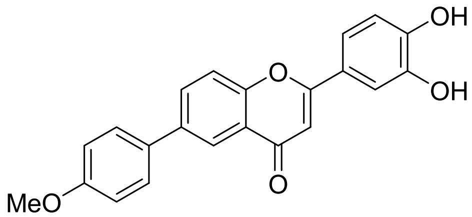


**SR31870**

The title compound was prepared following the same general protocol as described for Step 5, Example 1, using 2-(3,4-bis(benzyloxy)phenyl)-6-(4-methoxyphenyl)-4H-chromen-4-one. 1H NMR (400 MHz, $CD_3OD$): δ 8.31 (s, 1H), 8.07 (d, *J* = 8.1 Hz, 1H), 7.77-7.66 (m, 3H), 7.50 (d, *J* = 8.20 Hz, 2H), 7.08 (d, *J* = 8.1 Hz, 2H), 6.95 (d, *J* = 8.1 Hz, 1H), 6.79 (s, 1H), 3.87 (s, 3H).

**Example 10. Preparation of 2-(3,4-dihydroxyphenyl)-6-(3-(trifluoromethyl)phenyl)-4H-chromen-4-one (SR31865).**

Step 1: 2-(3,4-Bis(benzyloxy)phenyl)-6-(3-(trifluoromethyl)phenyl)-4H-chromen-4-one.

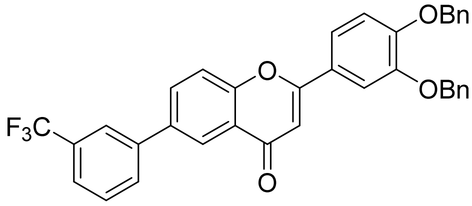


The title compound was prepared following the same general protocol as described for Step 3, Example 3, using (4-methoxyphenyl)boronic acid. $^1$H NMR (400 MHz, $CDCl_3$): δ 8.50 (d, *J* = 2.3 Hz, 1H), 7.55 (dd, *J* = 8.2 Hz, 2.1 Hz, 2H), 7.48 (dd, *J* = 8.1 Hz, 4.2Hz, 1H), 7.37 -7.17 (m, 15H), 7.95 (d, *J* = 8.2 Hz, 1H), 6.62 (s, 1H), 5.16 (d, *J* = 1.9 Hz, 4H).

Step 2: 2-(3,4-Dihydroxyphenyl)-6-(3-(trifluoromethyl)phenyl)-4H-chromen-4-one (**SR31865**).

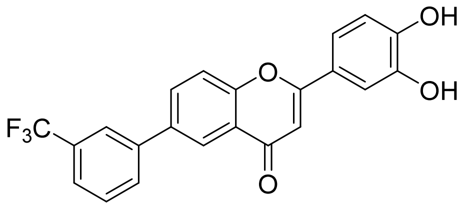


**SR31865**

The title compound was prepared following the same general protocol as described for Step 5, Example 1, using 2-(3,4-bis(benzyloxy)phenyl)-6-(3-(trifluoromethyl)phenyl)-4H-chromen-4-one. $^1$H NMR (400 MHz, $CD_3OD$): δ 8.39 (s, 1H), 8.15 (d, *J* = 8.2 Hz, 1H), 8.00 (s, 2H), 7.84-7.58 (m, 3H), 7.58-7.48 (m, 2H), 6.96 (d, *J* = 8.2 Hz, 1H), 6.80 (s, 1H).

**Example 11. Preparation of 6-bromo-2-(3,4-dihydroxyphenyl)-4H-chromen-4-one (SR31125).**

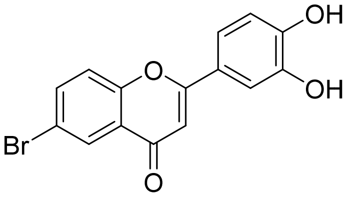


**SR31125**

The title compound was prepared following the same general protocol as described for Step 2, Example 5, using 2-(3,4-bis(benzyloxy)phenyl)-6-bromo-4H-chromen-4-one. ESI-MS (m/z): 332.5, 334.4 $[M+1]^+$. $^1H$ NMR (400 MHz, DMSO-$d_6$): δ 9.99 (s, 1H), 9.44 (s, 1H), 8.13 (d, *J* = 4.0 Hz, 1H), 8.01 (dd, *J* = 8.0 Hz, 4.0 Hz, 1H), 7.77 (d, *J* = 8.0 Hz, 1H), 7.52-7.48 (m, 2H), 6.94 (d, *J* = 8.0 Hz, 1H), 6.85 (s, 1H).

**Example 12. Preparation of 2-(3,4-dihydroxyphenyl)-6-hydroxy-4H-chromen-4-one (SR31128) and 2-(3,4-dihydroxyphenyl)-6-methoxy-4H-chromen-4-one (SR31129).**

Step 1: (*E*)-3-(3,4-Bis(benzyloxy)phenyl)-1-(2-hydroxy-5-methoxyphenyl)prop-2-en-1-one.

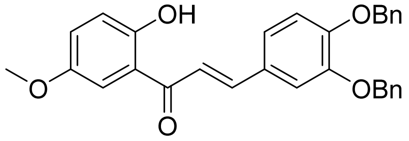


The title compound was prepared following the same general protocol as described for Step 3, Example 1, using 1-(2-hydroxy-5-methoxyphenyl)ethan-1-one. ESI-MS (m/z): 466.6 $[M+1]^+$.

Step 2: 2-(3,4-Bis(benzyloxy)phenyl)-6-methoxy-4H-chromen-4-one.

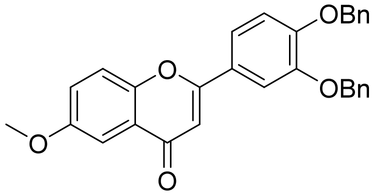


The title compound was prepared following the same general protocol as described for Step 4, Example 1, using (*E*)-3-(3,4-bis(benzyloxy)phenyl)-1-(2-hydroxy-5-methoxyphenyl)prop-2-en-1-one. ESI-MS (m/z): 464.6 $[M+1]^+$.

Step 3: 2-(3,4-Dihydroxyphenyl)-6-methoxy-4H-chromen-4-one (**SR31129**).

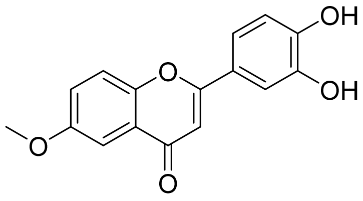


**SR31129**

The title compound was prepared following the same general protocol as described for Step 2, Example 5, using 2-(3,4-bis(benzyloxy)phenyl)-6-methoxy-4H-chromen-4-one. ESI-MS (m/z): 284.5 $[M+1]^+$. $^1$H NMR (400 MHz, DMSO-$d_6$): δ 9.89 (br s, 1H), 9.41 (br s, 1H), 7.73 (d, $J$ = 8.0 Hz, 1H), 7.50-7.41 (m, 4H), 6.94 (d, $J$ = 12.0 Hz, 1H), 6.75 (s, 1H), 3.90 (s, 3H).

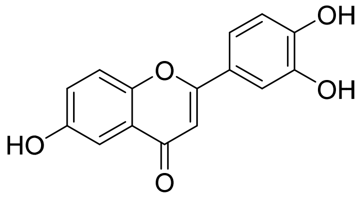


**SR31128**

2-(3,4-Dihydroxyphenyl)-6-hydroxy-4H-chromen-4-one (**SR31128**): ESI-MS (m/z): 270.5 $[M+1]^+$. $^1$H NMR (400 MHz, DMSO-$d_6$): δ 9.99 (br s, 1H), 9.85 (br s, 1H), 9.39 (br s, 1H), 7.62 (d, $J$ = 8.0 Hz, 1H), 7.46-7.43 (m, 2H), 7.33 (d, $J$ = 4.0 Hz, 1H), 7.26 (dd, $J$ = 8.0 Hz, 4.0 Hz, 1H), 6.93 (d, $J$ = 8.0 Hz, 1H), 6.70 (s, 1H).

**Example 13. Preparation of 2-(3,4-dihydroxyphenyl)-6-methyl-4H-chromen-4-one (SR31126).**

Step1: 2-(3,4-Bis(benzyloxy)phenyl)-6-methyl-4H-chromen-4-one.

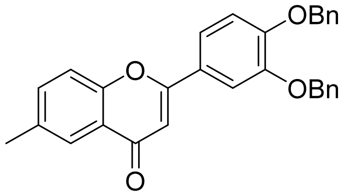


The title compound was prepared following the same general protocol as described for Step 3, Example 3, using 2,4,6-trimethyl-1,3,5,2,4,6-trioxatriborinane. ESI-MS (m/z): 448.6 $[M+1]^+$.

Step 2: 2-(3,4-Dihydroxyphenyl)-6-methyl-4H-chromen-4-one (**SR31126**).

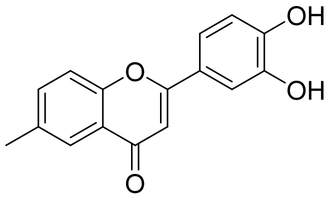

**SR31126**

The title compound was prepared following the same general protocol as described for Step 2, Example 5, using 2-(3,4-bis(benzyloxy)phenyl)-6-methyl-4H-chromen-4-one. ESI-MS (m/z): 268.5 $[M+1]^+$. $^1H$ NMR (400 MHz, DMSO-$d_6$): δ 9.91 (s, 1H), 9.43 (s, 1H), 7.86 (s, 1H), 7.60-7.57 (m, 2H), 7.48-7.46 (m, 2H), 6.94 (d, *J* = 8.0 Hz, 1H), 6.76 (s, 1H).

**Example 14. Preparation of 2-(3,4-dihydroxyphenyl)-6-(pyrrolidin-1-yl)-4H-chromen-4-one (SR31132).**

Step1: 2-(3,4-Bis(benzyloxy)phenyl)-6-methyl-4H-chromen-4-one.

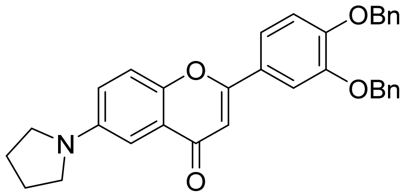

A mixture of 2-(3,4-bis(benzyloxy)phenyl)-6-bromo-4H-chromen-4-one (0.15 g, 0.292 mmol), pyrrolidine (0.031 g, 0.438 mmol), $Pd_2(dba)_3$ (0.027 g, 0.029 mmol) , Xantphos (0.051 g, 0.088 mmol) and $Cs_2CO_3$ (0.19 g, 0.584 mmol) in dioxane (1 mL) was degassed and stirred at 140°C overnight under nitrogen. The mixture was added water and EtOAc, the mixture was stirred and separated, the aqueous phase was extracted with EtOAc twice, the combined organic phases were washed with brine and dried with $Na_2SO_4$, filtered through silica gel to get the crude, which was purified by silica gel column to obtain the title compound. ESI-MS (m/z): 503.6 $[M+1]^+$.

Step2: 2-(3,4-Dihydroxyphenyl)-6-(pyrrolidin-1-yl)-4H-chromen-4-one (**SR31132**).

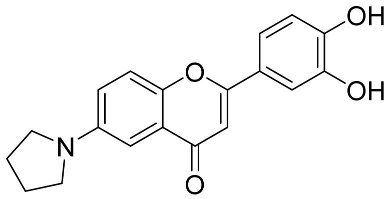

**SR31132**

The title compound was prepared following the same general protocol as described for Step 2, Example 5, using 2-(3,4-bis(benzyloxy)phenyl)-6-methyl-4H-chromen-4-one. ESI-MS (m/z):

323.6 [M+1]+. $^{1}H$ NMR (400 MHz, DMSO-$d_6$): δ 8.26 (br s, 2H), 7.62 (d, *J* = 12.0 Hz, 1H), 7.44-7.42 (m, 2H), 7.15-7.11 (m, 1H), 6.95-6.91 (m, 2H), 6.68 (s, 1H), 3.36-3.32 (m, 2H), 2.60-2.57 (m, 4H), 2.03-1.99 (m, 2H).

**Example 15. Preparation of 2-(3,4-dihydroxyphenyl)-7-hydroxy-4H-chromen-4-one (SR30998).**

Step 1: 1-(2-Hydroxy-4-(methoxymethoxy)phenyl)ethan-1-one.

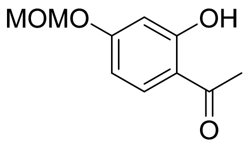


To 1-(2,4-dihydroxyphenyl)ethan-1-one (3.26 g, 21.42 mmol) and DIEA (9.3 mL, 53.55 mmol) in DCM (50 mL) was added slowly MOM-Cl (2.0 mL, 25.70 mmol). The mixture was stirred at room temperature for 1 h. The mixture was added water and DCM, the mixture was separated, and the aqueous phase was extracted with DCM twice, the combined organic phases were washed with brine and dried with $Na_2SO_4$, filtered through silica gel to get the crude, which was purified by silica gel column to obtain the title compound. $^{1}H$ NMR (400 MHz, $CDCl_3$): δ 12.61 (s, 1H), 7.65 (d, *J* = 8.0 Hz, 1H), 7.26 (s, 1H), 6.60 (d, *J* = 4.0 Hz, 1H), 6.54 (dd, *J* = 8.0 Hz, 4.0 Hz, 1H), 5.21 (s, 2H), 3.48 (s, 3H), 2.57 (s, 3H).

Step 2: (*E*)-3-(3,4-Bis(benzyloxy)phenyl)-1-(2-hydroxy-4-(methoxymethoxy)phenyl)prop-2-en-1-one.

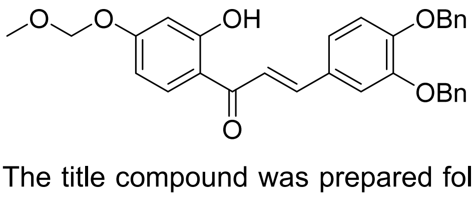


The title compound was prepared following the same general protocol as described for Step 1, Example 3, using 1-(2-hydroxy-4-(methoxymethoxy)phenyl)ethan-1-one. ESI-MS (m/z): 496.9[M+1]+.

Step 3: 2-(3,4-Bis(benzyloxy)phenyl)-7-hydroxy-4H-chromen-4-one.

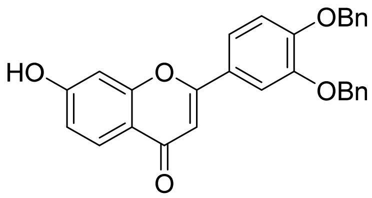

The title compound was prepared following the same general protocol as described for Step 4, Example 1, using (*E*)-3-(3,4-bis(benzyloxy)phenyl)-1-(2-hydroxy-4-(methoxymethoxy)phenyl)prop-2-en-1-one. ESI-MS (m/z): 450.9 $[M+1]^+$.

Step 4: 2-(3,4-Dihydroxyphenyl)-7-hydroxy-4H-chromen-4-one (**SR30998**).

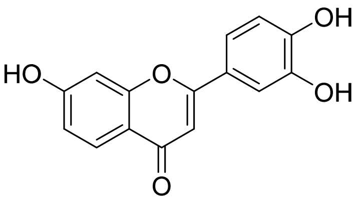


**SR30998**

The title compound was prepared following the same general protocol as described for Step 5, Example 1, using 2-(3,4-bis(benzyloxy)phenyl)-7-hydroxy-4H-chromen-4-one. ESI-MS (m/z): 270.7 $[M+1]^+$.

**Example 16. Preparation of 2-(3,4-dihydroxyphenyl)-7-methoxy-4H-chromen-4-one (SR29385).**

Step 1: (*E*)-3-(3,4-Bis(benzyloxy)phenyl)-1-(2-hydroxy-4-methoxyphenyl)prop-2-en-1-one.

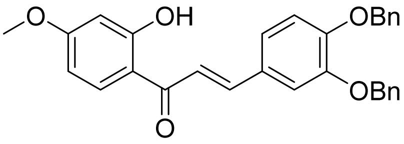


The title compound was prepared following the same general protocol as described for Step 3, Example 1, using 1-(2-hydroxy-4-methoxyphenyl)ethan-1-one. ESI-MS (m/z): 466.6 $[M+1]^+$. $^1$H NMR (400 MHz, $CDCl_3$): δ 7.78 (s, 1H), 7.76 (d, *J* = 4.0 Hz, 1H), 7.49-7.30 (m, 13H), 6.93 (d, *J* = 8.0 Hz, 1H), 6.52-6.45 (m, 2H), 5.22 (s, 4H), 3.86 (s, 3H).

Step 2: 2-(3,4-Bis(benzyloxy)phenyl)-7-methoxy-4H-chromen-4-one.

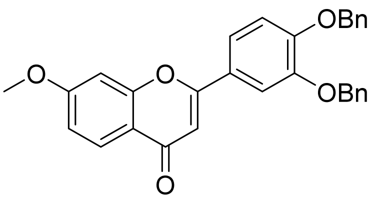


The title compound was prepared following the same general protocol as described for Step 4, Example 1, using (*E*)-3-(3,4-bis(benzyloxy)phenyl)-1-(2-hydroxy-4-methoxyphenyl)prop-2-en-1-one. ESI-MS (m/z): 464.5 $[M+1]^+$. $^1$H NMR (400 MHz, $CD_3CN$): δ 7.99 (d, *J* = 8.0 Hz, 1H), 7.63-

7.59 (m, 2H), 7.50-7.34 (m, 10H), 7.18-7.14 (m, 2H), 7.01 (dd, *J* = 8.0 Hz, 2.0 Hz, 1H), 6.66 (s, 1H), 5.25 (s, 2H), 5.22 (s, 2H), 3.94 (s, 3H).

Step 3: 2-(3,4-Dihydroxyphenyl)-7-methoxy-4H-chromen-4-one (**SR29385**).

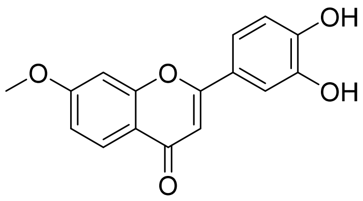


**SR29385**

The title compound was prepared following the same general protocol as described for Step 5, Example 1, using 2-(3,4-bis(benzyloxy)phenyl)-7-methoxy-4H-chromen-4-one. ESI-MS (m/z): 284.9 $[M+1]^+$. $^1H$ NMR (400 MHz, DMSO-$d_6$): δ 9.90 (s, 1H), 9.38 (s, 1H), 7.96 (d, *J* = 8.0 Hz, 1H), 7.48-7.45 (m, 2H), 7.28 (d, *J* = 4.0 Hz, 1H), 7.08 (dd, *J* = 8.0 Hz, 4.0 Hz, 1H), 6.94 (d, *J* = 12.0 Hz, 1H), 6.70 (s, 1H), 3.96 (s, 3H).

**Example 17. Preparation of 2-(3,4-dihydroxyphenyl)-6,7-dimethoxy-4H-chromen-4-one (SR29386).**

Step 1: (*E*)-3-(3,4-Bis(benzyloxy)phenyl)-1-(2-hydroxy-4,5-dimethoxyphenyl)prop-2-en-1-one.

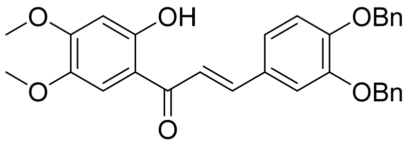


The title compound was prepared following the same general protocol as described for Step 1, Example 3, using 1-(2-hydroxy-4,5-dimethoxyphenyl)ethan-1-one. ESI-MS (m/z): 496.9$[M+1]^+$.

Step 2: 2-(3,4-Bis(benzyloxy)phenyl)-6,7-dimethoxy-4H-chromen-4-one.

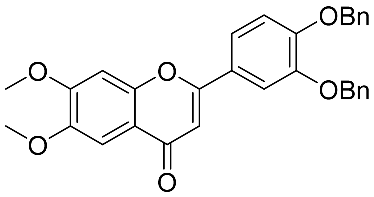


The title compound was prepared following the same general protocol as described for Step 4, Example 1, using (*E*)-3-(3,4-bis(benzyloxy)phenyl)-1-(2-hydroxy-4,5-dimethoxyphenyl)prop-2-en-1-one. ESI-MS (m/z): 494.9 $[M+1]^+$.

Step 3: 2-(3,4-Dihydroxyphenyl)-6,7-dimethoxy-4H-chromen-4-one (**SR29386**).

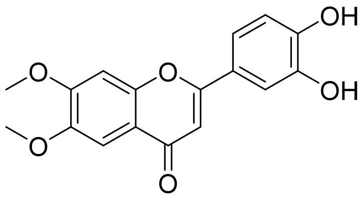

**SR29386**

The title compound was prepared following the same general protocol as described for Step 5, Example 1, using 2-(3,4-bis(benzyloxy)phenyl)-7-hydroxy-4H-chromen-4-one. ESI-MS (m/z): 314.6 $[M+1]^+$. $^1H$ NMR (400 MHz, DMSO-$d_6$): δ 7.46-7.44 (m, 2H), 7.39 (br s, 1H), 7.34 (br s, 1H), 6.93 (d, *J* = 8.0 Hz, 1H), 6.70 (s, 1H), 3.97 (s, 3H), 3.90 (s, 3H).

**Example 18. Preparation of 2-(3,4-dihydroxyphenyl)-6-(dimethylamino)-4H-chromen-4-one (SR29387).**

Step 1: (*E*)-3-(3,4-Bis(benzyloxy)phenyl)-1-(5-(dimethylamino)-2-hydroxyphenyl)prop-2-en-1-one.

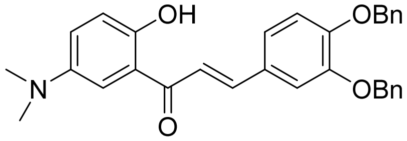

The title compound was prepared following the same general protocol as described for Step 1, Example 3, using 1-(5-(dimethylamino)-2-hydroxyphenyl)ethan-1-one. $^1H$ NMR (400 MHz, $CDCl_3$): δ 12.33 (s, 1H), 7.76 (d, *J* = 16.0 Hz, 1H), 7.48-7.34 (m, 13H), 7.26-7.22 (m, 2H), 6.99-6.91 (m, 2H), 5.24 (s, 4H), 2.92 (s, 6H).

Step 2: 2-(3,4-Bis(benzyloxy)phenyl)-6-(dimethylamino)-4H-chromen-4-one.

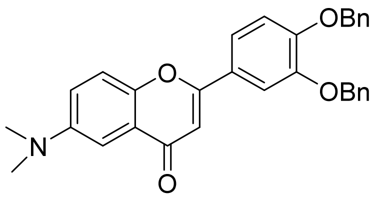

The title compound was prepared following the same general protocol as described for Step 4, Example 1, using (*E*)-3-(3,4-bis(benzyloxy)phenyl)-1-(5-(dimethylamino)-2-hydroxyphenyl)prop-2-en-1-one. $^1H$ NMR (400 MHz, $CDCl_3$): δ 7.51-7.31 (m, 14H), 7.16 (d, *J* = 8.0 Hz, 1H), 7.02 (d, *J* = 12.0 Hz, 1H), 6.67 (s, 1H), 5.25 (s, 4H), 3.04 (s, 6H).

Step 3: 2-(3,4-Dihydroxyphenyl)-6-(dimethylamino)-4H-chromen-4-one (**SR29387**).

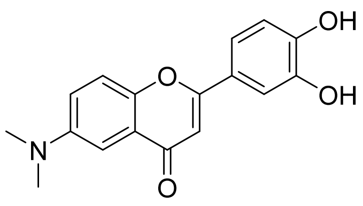

**SR29387**

The title compound was prepared following the same general protocol as described for Step 5, Example 1, using 2-(3,4-bis(benzyloxy)phenyl)-6-(dimethylamino)-4H-chromen-4-one. ESI-MS (m/z): 297.6 $[M+1]^+$. $^1$H NMR (400 MHz, DMSO-$d_6$): δ 9.84(s, 1H), 9.40 (s, 1H), 7.62 (d, *J* = 8.0 Hz, 1H), 7.45-7.41 (m, 2H), 7.32 (dd, *J* = 8.0 Hz, 4.0 Hz, 1H), 7.11 (d, *J* = 4.0 Hz, 1H), 6.93 (d, *J* = 12.0 Hz, 1H), 6.70 (s, 1H), 3.02 (s, 6H).

**Example 19. Preparation of N-(2-(3,4-dihydroxyphenyl)-4-oxo-4H-chromen-7-yl)acetamide (SR29388).**

Step 1: (*E*)-N-(4-(3-(3,4-bis(benzyloxy)phenyl)acryloyl)-3-hydroxyphenyl)acetamide.

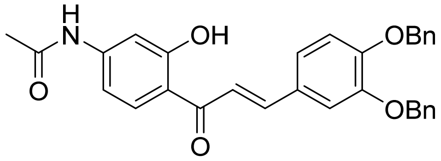

The title compound was prepared following the same general protocol as described for Step 1, Example 3, using N-(4-acetyl-3-hydroxyphenyl)acetamide. $^1$H NMR (400 MHz, $CDCl_3$): δ 13.20 (s, 1H), 7.82-7.76 (m, 2H), 7.49-7.31 (m, 13H), 7.24-7.19 (m, 2H), 7.07 (d, *J* = 4.0 Hz, 1H), 6.95 (d, *J* = 8.0 Hz, 1H), 5.23 (s, 4H), 2.20 (s, 3H).

Step 2: N-(2-(3,4-bis(benzyloxy)phenyl)-4-oxo-4H-chromen-7-yl)acetamide.

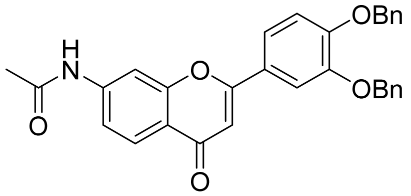

The title compound was prepared following the same general protocol as described for Step 4, Example 1, using (*E*)-N-(4-(3-(3,4-bis(benzyloxy)phenyl)acryloyl)-3-hydroxyphenyl)acetamide. ESI-MS (m/z): 492.0 $[M+1]^+$.

Step 3: N-(2-(3,4-dihydroxyphenyl)-4-oxo-4H-chromen-7-yl)acetamide.

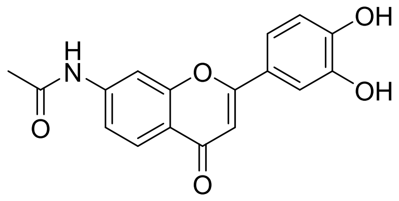

**SR29388**

The title compound was prepared following the same general protocol as described for Step 5, Example 1, using 2-(3,4-bis(benzyloxy)phenyl)-6-(dimethylamino)-4H-chromen-4-one. ESI-MS (m/z): 311.9 $[M+1]^+$.

**Example 20. Preparation of 2-(3,4-dihydroxyphenyl)-7-(dimethylamino)-4H-chromen-4-one (SR29389).**

Step 1: 7-Amino-2-(3,4-bis(benzyloxy)phenyl)-4H-chromen-4-one.

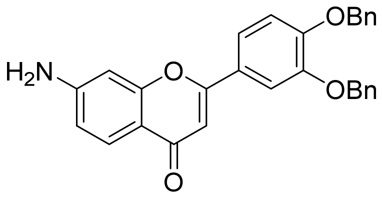

N-(2-(3,4-bis(benzyloxy)phenyl)-4-oxo-4H-chromen-7-yl)acetamide (0.4 g) in EtOH (20 mL)was added HCl (2 M, 5 mL) and then stirred at 100 °C oil bath overnight. The mixture was cooled and concentrated for the next step with no purification. ESI-MS (m/z): 450.9 $[M+1]^+$.

Step 2: 2-(3,4-Bis(benzyloxy)phenyl)-7-(dimethylamino)-4H-chromen-4-one.

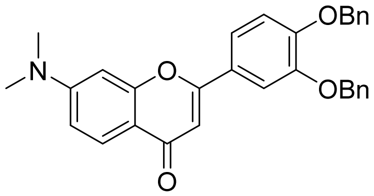

7-Amino-2-(3,4-bis(benzyloxy)phenyl)-4H-chromen-4-one (0.18 g, 0.4 mmol) and $K_2CO_3$ (0.166 g, 1.2 mmol) in DMF (2 mL) was added MeI (0.07 mL, 1.2 mmol). The mixture was stirred at 100 ℃ oil bath overnight. The mixture was added water and EtOAc, the mixture was separated, and the aqueous phase was extracted with EtOAc twice, the combined organic phases were washed with brine and dried with $Na_2SO_4$, filtered through silica gel to get the crude, which was purified by silica gel column to obtain the title compound. ESI-MS (m/z): 480.0 $[M+1]^+$.

Step 3: 2-(3,4-Dihydroxyphenyl)-7-(dimethylamino)-4H-chromen-4-one.

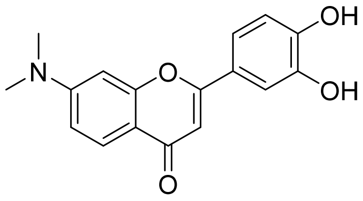


**SR29389**

The title compound was prepared following the same general protocol as described for Step 5, Example 1, using 2-(3,4-bis(benzyloxy)phenyl)-7-(dimethylamino)-4H-chromen-4-one. ESI-MS (m/z): 297.6 $[M+1]^+$. $^1$H NMR (400 MHz, DMSO-$d_6$): δ 9.82(br s, 1H), 9.37 (br s, 1H), 7.82 (d, *J* = 8.0 Hz, 1H), 7.44-7.40 (m, 2H), 6.91 (dd, *J* = 8.0 Hz, 4.0 Hz, 2H), 6.74 (d, *J* = 4.0 Hz, 1H), 6.56 (s, 1H), 3.11 (s, 6H).

**Example 21. Preparation of 2-(3,4-dihydroxyphenyl)-6-isopropoxy-4H-chromen-4-one (SR31124).**

Step 1: (*E*)-3-(3,4-Bis(benzyloxy)phenyl)-1-(2-hydroxy-5-isopropoxyphenyl)prop-2-en-1-one.

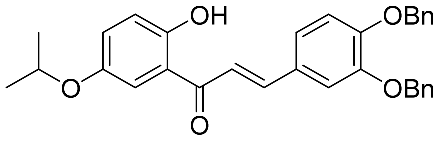


The title compound was prepared following the same general protocol as described for Step 1, Example 3, using 1-(2-hydroxy-5-isopropoxyphenyl)ethan-1-one. ESI-MS (m/z): 494.8 $[M+1]^+$.

Step 2: 2-(3,4-Bis(benzyloxy)phenyl)-6-isopropoxy-4H-chromen-4-one.

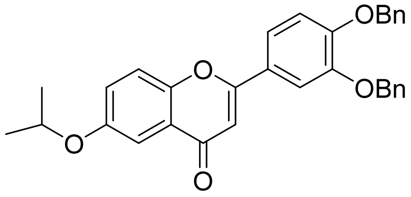


The title compound was prepared following the same general protocol as described for Step 4, Example 1, using (*E*)-3-(3,4-bis(benzyloxy)phenyl)-1-(2-hydroxy-5-isopropoxyphenyl)prop-2-en-1-one. ESI-MS (m/z): 492.9 $[M+1]^+$.

Step 3: 2-(3,4-Dihydroxyphenyl)-6-isopropoxy-4H-chromen-4-one (**SR31124**).

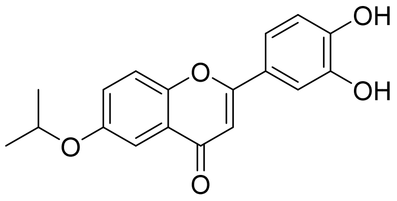

**SR31124**

The title compound was prepared following the same general protocol as described for Step 5, Example 1, using 2-(3,4-bis(benzyloxy)phenyl)-6-(dimethylamino)-4H-chromen-4-one. ESI-MS (m/z): 312.5 $[M+1]^+$. $^1$H NMR (400 MHz, DMSO-$d_6$): δ 9.90 (br s, 1H), 9.42 (br s, 1H), 7.70 (d, *J* = 8.0 Hz, 1H), 7.48-7.38 (m, 4H), 6.94 (d, *J* = 12.0 Hz, 1H), 6.76 (s, 1H), 4.78-4.72 (m, 1H), 1.34 (d, *J* = 4.0 Hz, 6H).

**Example 22. Preparation of 2-(3,4-dimethoxyphenyl)-6-isopropyl-4H-chromen-4-one (SR32171).**

Step 1: (*E*)-3-(3,4-Dimethoxyphenyl)-1-(2-hydroxy-5-isopropylphenyl)prop-2-en-1-one.

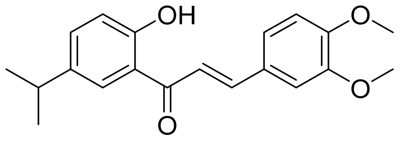

The title compound was prepared following the same general protocol as described for Step 3, Example 1, using 3,4-dimethoxybenzaldehyde. ESI-MS (m/z): 327.1 $[M+1]^+$.

Step2: 2-(3,4-Dimethoxyphenyl)-6-isopropyl-4H-chromen-4-one (**SR32171**).

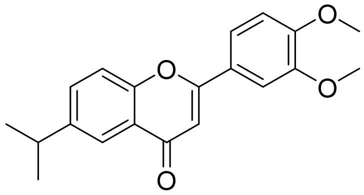

**SR32171**

The title compound was prepared following the same general protocol as described for Step 4, Example 1, using (*E*)-3-(3,4-dimethoxyphenyl)-1-(2-hydroxy-5-isopropylphenyl)prop-2-en-1-one. ESI-MS (m/z): 324.9 $[M+1]^+$.

**Example 24. Preparation of 2-(3,4-dihydroxyphenyl)-6-(3-methoxyphenyl)-4H-chromen-4-one (SR31864).**

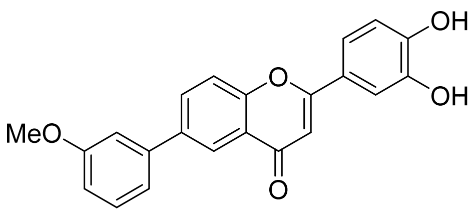

**SR31864**

The title compound was prepared following the same general protocols as described for Example 3. $^1$H NMR (400 MHz, DMSO-$d_6$): δ 9.95 (s, 1H), 8.99 (s, 1H), 8.29 (d, *J* = 4.0 Hz, 1H), 8.20 (dd, *J* = 8.0 Hz, 4.0 Hz, 1H), 7.88 (d, *J* = 12.0 Hz, 1H), 7.56-7.54 (m, 2H), 7.48 (d, *J* = 8.0 Hz, 1H), 7.41-7.37 (m, 1H), 7.34 (m, 1H), 7.05 (dd, *J* = 8.0 Hz, 4.0 Hz, 1H), 6.98 (d, *J* = 8.0 Hz, 1H), 6.87 (s, 1H), 3.92 (s, 3H).

**Example 25. Preparation of 2-(3,4-dihydroxyphenyl)-6-(pyridin-3-yl)-4H-chromen-4-one (SR31866).**

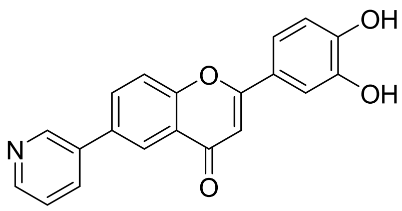

**SR31866**

The title compound was prepared following the same general protocols as described for Example 3. $^1$H NMR (400 MHz, $CD_3OD$): δ 9.45 (s, 1H), 8.99 (s, 1H), 8.63 (d, *J* = 4.0 Hz, 1H), 8.28 (d, *J* = 4.0 Hz, 1H), 8.21-8.18 (m, 2H), 7.88 (d, *J* = 8.0 Hz, 1H), 7.65-7.45 (m, 3H), 6.94 (d, *J* = 8.0 Hz, 1H), 6.83 (s, 1H).

**Example 26. Preparation of 6-(3,5-dichlorophenyl)-2-(3,4-dihydroxyphenyl)-4H-chromen-4-one (SR31867).**

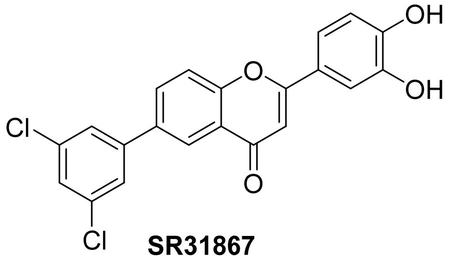

**SR31867**

The title compound was prepared following the same general protocols as described for Example 3. $^1$H NMR (400 MHz, $CD_3OD$): δ 8.28 (d, *J* = 4.0 Hz, 1H), 8.20 (dd, *J* = 8.0 Hz, 4.0 Hz, 1H), 8.0

(d, *J* = 4.0 Hz, 1H), 7.90-7.84 (m, 2H), 7.68-7.66 (m, 1H), 7.50-7.47 (m, 2H), 6.94 (d, *J* = 8.0 Hz, 1H), 6.84 (s, 1H).

**Example 27. Preparation of 2-(3,4-dihydroxyphenyl)-6-(pyridin-4-yl)-4H-chromen-4-one (SR31869).**

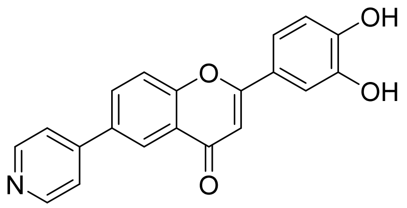

**SR31869**

The title compound was prepared following the same general protocols as described for Example 3. $^1$H NMR (400 MHz, $CD_3OD$): δ 8.69 (d, *J* = 8.0 Hz, 2H), 8.44 (s, 1H), 8.25 (d, *J* = 8.0 Hz, 1H), 7.89 (d, *J* = 8.0 Hz, 1H), 7.87-7.83 (m, 2H), 7.53-7.47 (m, 1H), 6.94 (d, *J* = 8.0 Hz, 1H), 6.87 (s, 1H).

**Example 28. Preparation of 6-cyclopropyl-2-(3,4-dihydroxyphenyl)-4H-chromen-4-one (SR31873).**

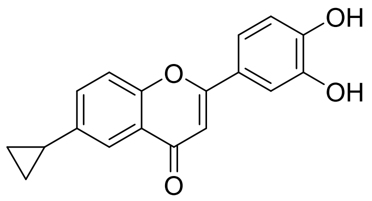

**SR31873**

The title compound was prepared following the same general protocols as described for Example 3. $^1$H NMR (400 MHz, $CD_3OD$): δ 8.70 (d, *J* = 4.0 Hz, 1H), 7.62 (d, *J* = 8.0 Hz, 1H), 7.52 (d, *J* = 8.0 Hz, 1H), 7.45-7.43 (m, 2H), 6.90 (d, *J* = 8.0 Hz, 1H), 6.73 (s, 1H), 2.14-2.07 (m, 1H), 1.06-1.00 (m, 2H), 0.77-0.73 (m, 2H).

**Example 29. Preparation of 2-(3,4-dihydroxyphenyl)-6-(tetrahydro-2H-pyran-4-yl)-4H-chromen-4-one (SR31884).**

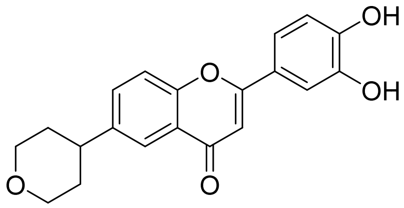

**SR31884**

The title compound was prepared following the same general protocols as described for Example 3. $^1$H NMR (400 MHz, $CD_3OD$): δ 8.21 (d, *J* = 8.0 Hz, 1H), 7.95 (t, *J* = 8.0 Hz, 1H), 7.86 (d, *J* = 8.0 Hz, 1H), 7.60 (d, *J* = 8.0 Hz, 1H), 7.57 (s, 1H), 7.03 (s, 1H), 6.99 (d, *J* = 8.0 Hz, 1H), 3.70-3.60 (m, 4H), 2.23-2.16 (m, 1H), 1.45-1.26 (m, 2H), 0.88-0.85 (m, 2H).

**Example 30. Preparation of 6-cyclobutyl-2-(3,4-dihydroxyphenyl)-4H-chromen-4-one (SR32164).**

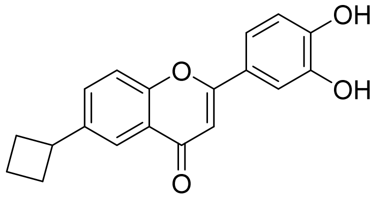

**SR32164**

The title compound was prepared following the same general protocols as described for Example 3. $^1$H NMR (400 MHz, $CD_3OD$): δ 8.21 (d, *J* = 4.0 Hz, 1H), 7.92 (dd, *J* = 8.0 Hz, 4.0 Hz, 1H), 7.66 (d, *J* = 8.0 Hz, 1H), 7.47-7.44 (m, 2H), 6.94 (d, *J* = 8.0 Hz, 1H), 6.77 (s, 1H), 3.71-3.54 (m, 1H), 1.35-1.27 (m, 2H), 0.93-0.88 (m, 4H).

**Example 31. Preparation of 2-(3,4-dihydroxyphenyl)-6-ethyl-4H-chromen-4-one (SR32165).**

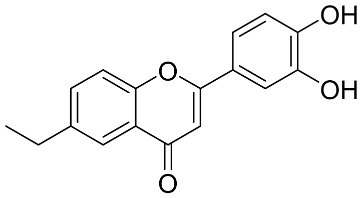

**SR32165**

The title compound was prepared following the same general protocols as described for Example 3. $^1$H NMR (400 MHz, $CD_3OD$): δ 7.56-7.55 (m, 2H), 7.47 (d, *J* = 8.0 Hz, 1H), 7.36-7.32 (m, 2H), 6.83 (d, *J* = 4.0 Hz, 1H), 6.64 (s, 1H), 2.71-2.65 (q, *J* = 8.0 Hz, 2H), 1.20 (t, *J* = 8.0 Hz, 3H).

**Example 32. Preparation of 2-(3,4-dihydroxyphenyl)-6-propyl-4H-chromen-4-one (SR32166).**

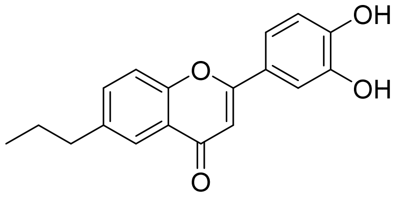

**SR32166**

The title compound was prepared following the same general protocols as described for Example 3. $^1H$ NMR (400 MHz, $CD_3OD$): δ 8.05 (d, *J* = 8.0 Hz, 2H), 7.95 (d, *J* = 12.0 Hz, 1H), 7.63-7.60 (m, 2H), 7.30 (d, *J* = 8.0 Hz, 1H), 6.83 (d, *J* = 8.0 Hz, 1H), 6.25 (s, 1H), 3.06-2.93 (m, 2H), 1.67-1.57 (m, 2H), 0.88 (t, *J* = 8.0 Hz, 3H).

**Example 33. Preparation of 2-(3,4-dihydroxyphenyl)-6-isobutyl-4H-chromen-4-one (SR32167).**

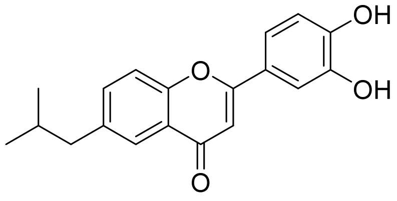

**SR32167**

The title compound was prepared following the same general protocols as described for Example 3. $^1H$ NMR (400 MHz, $CD_3OD$): δ 7.81(s, 1H), 7.55-7.48 (m, 2H), 7.46 (d, *J* = 8.0 Hz, 1H), 7.34 (m, 1H), 6.84 (d, *J* = 8.0 Hz, 1H), 6.71 (s, 1H), 2.54 (d, *J* = 8.0 Hz, 2H), 1.88-1.81 (m, 1H), 0.85 (d, *J* = 4.0 Hz, 6H).

**Example 34. Preparation of 6-(sec-butyl)-2-(3,4-dihydroxyphenyl)-4H-chromen-4-one (SR32168).**

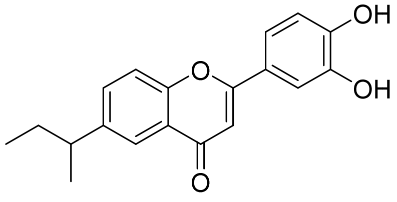

**SR32168**

The title compound was prepared following the same general protocols as described for Example 3. $^1H$ NMR (400 MHz, $CD_3OD$): δ 8.16 (d, *J* = 8.0 Hz, 1H), 7.86 (dd, *J* = 8.0 Hz, 4.0 Hz, 1H), 7.75

(d, $J$ = 8.0 Hz, 1H), 7.53-7.49 (m, 2H), 6.95 (d, $J$ = 8.0 Hz, 1H), 6.82 (s, 1H), 2.53-2.50 (m, 1H), 1.29-1.23 (m, 5H), 0.91 (t, $J$ = 8.0 Hz, 3H).

**Example 35. Preparation of 6-benzyl-2-(3,4-dihydroxyphenyl)-4H-chromen-4-one (SR32169).**

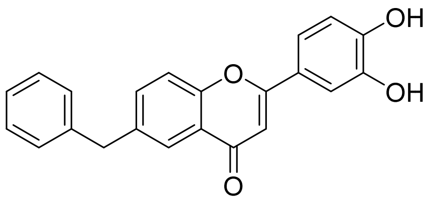


**SR32169**

The title compound was prepared following the same general protocols as described for Example 3. $^1H$ NMR (400 MHz, $CD_3OD$): δ 7.84 (br s, 1H), 7.61-7.49 (m, 2H), 7.41-7.30 (m, 3H), 7.22-7.08 (m, 4H), 6.83 (d, $J$ = 8.0 Hz, 1H), 6.64 (s, 1H), 5.39 (s, 2H).
